# Biosensor-Driven Strain Engineering Reveals Key Cellular Processes for Maximizing Isoprenol Production in *Pseudomonas putida*

**DOI:** 10.1101/2025.03.18.643695

**Authors:** Javier Menasalvas, Shawn Kulakowski, Yan Chen, Jennifer W. Gin, Emine Akyuz Turumtay, Nawa Raj Baral, Morgan A. Apolonio, Alex Rivier, Ian S. Yunus, Megan E. Garber, Corinne D. Scown, Paul D. Adams, Taek Soon Lee, Ian K. Blaby, Edward E. K. Baidoo, Christopher J. Petzold, Thomas Eng, Aindrila Mukhopadhyay

**Author notes:** correspondence: Thomas Eng; Aindrila Mukhopadhyay.

## Abstract

Synthetic biology tools have accelerated the generation of simple mutants, but combinatorial testing remains challenging. High-throughput methods struggle translating from proof-of-principle molecules to advanced bioproducts. We address this challenge with a biosensor-driven strategy for enhanced isoprenol production in *Pseudomonas putida*, a key precursor for sustainable aviation fuel and platform chemicals. This biosensor leverages *P. putida*’s native response to short-chain alcohols via a previously uncharacterized hybrid histidine kinase signaling cascade. Refactoring the biosensor for a conditional growth-based selection enabled identification of competing cellular processes with a ∼16,500-member CRISPRi-library. An iterative combinatorial strain engineering approach yielded an integrated *P. putida* strain producing ∼900 mg/L isoprenol in glucose minimal medium, a 36-fold increase. Ensemble -omics analysis revealed metabolic rewiring, including amino acid accumulation as key drivers of enhanced production. Techno-economic analysis elucidated the path to economic viability and confirmed the benefits of adding amino acids outweigh the additional costs. This study establishes a robust biosensor driven approach for optimizing other heterologous pathways, accelerating microbial cell factory development.

**GRAPHICAL ABSTRACT:** 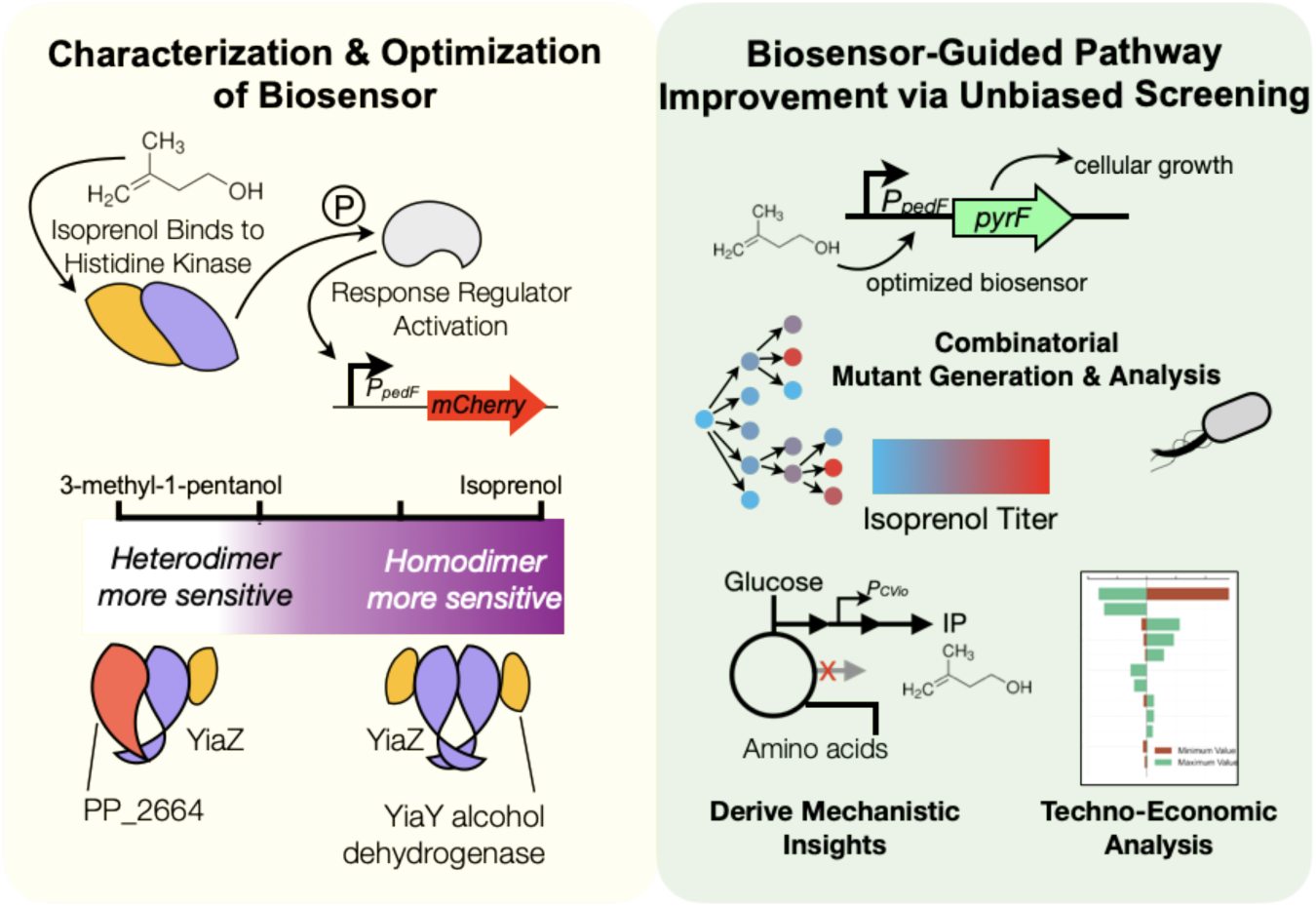

## INTRODUCTION

Microbially-derived fuels offer a strategic opportunity to enhance energy security, reduce the environmental footprint of industrialization, and drive innovation in biomanufacturing (*1*, *2*). Synthetic biology can produce advanced fuel molecules with superior combustion properties (*3–5*). However, the development of biofuel molecules (*6*, *7*), particularly short to medium chain alcohols, is hindered by several challenges. Unlike other products (*8*, *9*), alcohols have physical properties that impede development cycles from both analytical and biological standpoints. Analytically, alcohols are colorless and volatile molecules. These properties make their measurement challenging with high throughput methods requiring tradeoffs between speed, cost, and measurement accuracy for biological samples (*10*). For accurate quantification, the gold standard is gas chromatography (*11*, *12*), despite it being a serial analysis method prone to operator fatigue and error. Heterologous production of alcohols can be limited due to their inherent inhibitory nature, which causes growth defects (*13*, *14*). Expression of heterologous pathways can also lead to the accumulation of toxic intermediate pathway metabolites in the host organism (*15–17*). The additional burden in these strains often result in phenotypic drift (*18*) where production is lost after several generations of growth. Unpredictable performance increases the risk when evaluating a strain for industrial scaleup (*19*). Therefore, it is crucial to develop new tools, specific for the challenges of screening alcohols. This will enable the rapid development of new biofuel production processes that can meet the demands of emerging biorefinery configurations.

We evaluated the production of isoprenol, a platform commodity chemical (*20*) with attractive properties specifically as a diesel fuel blendstock (*21*) and precursor for the bio-jetfuel DMCO (*22*, *23*), across a number of microbial hosts (*24–27*). While each host has its strengths and weaknesses, *Pseudomonas putida* KT2440 was initially discouraging for use as a microbial chassis because it consumed the final product (*28*). The stepwise catabolism of isoprenol into acetyl-CoA was proposed using RB-TnSeq functional genomics data (*28*). However, we recognized that *P. putida*’*s* ability to catabolize isoprenol could be leveraged as an advantage, rather than a limitation: we could refactor the isoprenol-responsive signaling cascade based off the natural catabolic pathway and provide a readout for intracellular isoprenol levels. This approach could also enable screening methodologies providing alternatives to traditional strain engineering methods.

We adhere to a common definition for the term “biosensor”: a system in which a ligand is recognized by a transcriptional activator which then binds to a cognate DNA sequence, driving transcription of a downstream reporter gene (*29*). The resulting gene expression should increase linearly in response to the initial ligand concentration. There are undoubtedly other biosensor modalities, such as directly converting the ligand into a colored molecule (*30*), but the advantage of a biosensor that activates a genetic circuit is that the response can also be integrated back into cellular physiology for high throughput selection with growth based assays. For example, strains can be devised where cell growth is concomitant with the increased production of the ligand.

While understanding the molecular interactions between ligands and their partner proteins is important for biosensor design, crystallographic evidence for short chain alcohol binding is scarce (*31*). The absence of structural information necessitates alternative strategies for biosensor development. For example, Bahls *et. al.* recently developed an isopentanol (3-methyl-1-butanol) biosensor by error-prone PCR mutagenesis of *P. putida* GP01 *alkS* to broaden its substrate specificity in *E. coli*(*32*). Other reports on isopentanol using rational engineering approaches have also demonstrated isopentanol titers exceeding 9.5 g/L in batch mode cultures (*33*) gene targets. Biosensor-derived findings can be synergistic or additive with other approaches, providing a high-throughput route to such genome wide targets necessary for higher titers.

Here we show the development and characterization of an isoprenol biosensor, mediated by a native *P. putida* hybrid histidine kinase (abbreviated HHK) signaling cascade that is responsive to short chain alcohols including isoprenol and diols. After delineating its core components, we refactor the biosensor to identify bottleneck processes limiting heterologous isoprenol production using a pooled dCpf1/dCas12a gRNA library targeting nearly all ORFs in the genome. We generated and screened over 165 combinatorial mutants including 70 new gene loci targeted for deletion or overexpression, integrating known rational engineering targets with discoveries that emerged from our unbiased biosensor screen. An integrated -omics analysis of high producer strains revealed previously unknown intracellular amino acid carbon sinks enhancing isoprenol titer in production phase and confirmed experimentally with a medium analysis. Our modular methods provide a step towards commercial viability and are relevant to other analytically and biologically challenging molecules for biotechnology applications.

## RESULTS

### Development of a Dose-Dependent Isoprenol Biosensor

We developed a generalizable approach to building biosensors from RB-TnSeq fitness data (**Figure 1A, Supplemental Note 1**). This method builds upon an earlier report (*34*) that adds an RBS optimization step to tune the exogenous ligand concentration to a proportionate fluorescent response. Using our updated assay we searched for transcriptional regulators from existing RB-TnSeq datasets to identify genes necessary for growth in the presence of exogenously added isoprenol (*28*, *35*) (**Figure 1B**). We identified several promising signaling systems elements that hinted at an inducible gene expression profile using our co-fitness analysis. Cross-referencing the RB-TnSeq datasets with predicted ethanol degradation pathways in *Pseudomonads* (*36, 37*), we identified several two component signaling systems for analysis (**Figure 1B**), including a HHK *yiaZ* (or PP_2683), a response regulator PP_2665 and several metabolic enzymes including PP_2674, *pedF* and *yiaY* (or PP_2682) (**Figure 1B**). The specific protein components of a putative isoprenol signaling cascade were not clear as there were several probable kinases and regulators acting at any upstream promoter element for these encoded catabolic steps.

To examine a potential ligand-induced signaling cascade, we tested if the upstream *pedF* DNA sequence was responsive to isoprenol via a fluorescent reporter (**Figure 1C**). We selected a 254bp non-coding DNA sequence located between genes *pedF* and *pedE* to serve as the promoter sequence upstream of the *mCherry* sequence. The RBS placed here between the promoter and *mCherry* sequence was previously described (*38*). After transforming the *P. putida* wild-type strain with the *P_pedF_-RBS-mCherry* plasmid construct, strains were back-diluted into fresh media containing a range of isoprenol concentrations (0.62 to 1 g/L) to determine the fold isoprenol response (**Materials and Methods**). The lack of an RBS sequence impaired biosensor activation with poor dose-response, linearity, and reproducibility across replicates (**Figure 1C**). However with an optimized RBS sequence, the synthetic construct showed a 20x increase in signal over an uninduced control using 1 g/L exogenous isoprenol by 40 hours post exogenous isoprenol addition (**Figure 1C**). To demonstrate the general utility of this method, we include a second biosensor for the monomeric aromatic *p-*CA, where an optimized RBS improved the fluorescent activation from 5x to 200x (**Supplemental Note 1**). We concluded the dose-dependent response both constructs demonstrated the generalizability of this process.

**Figure 1:**
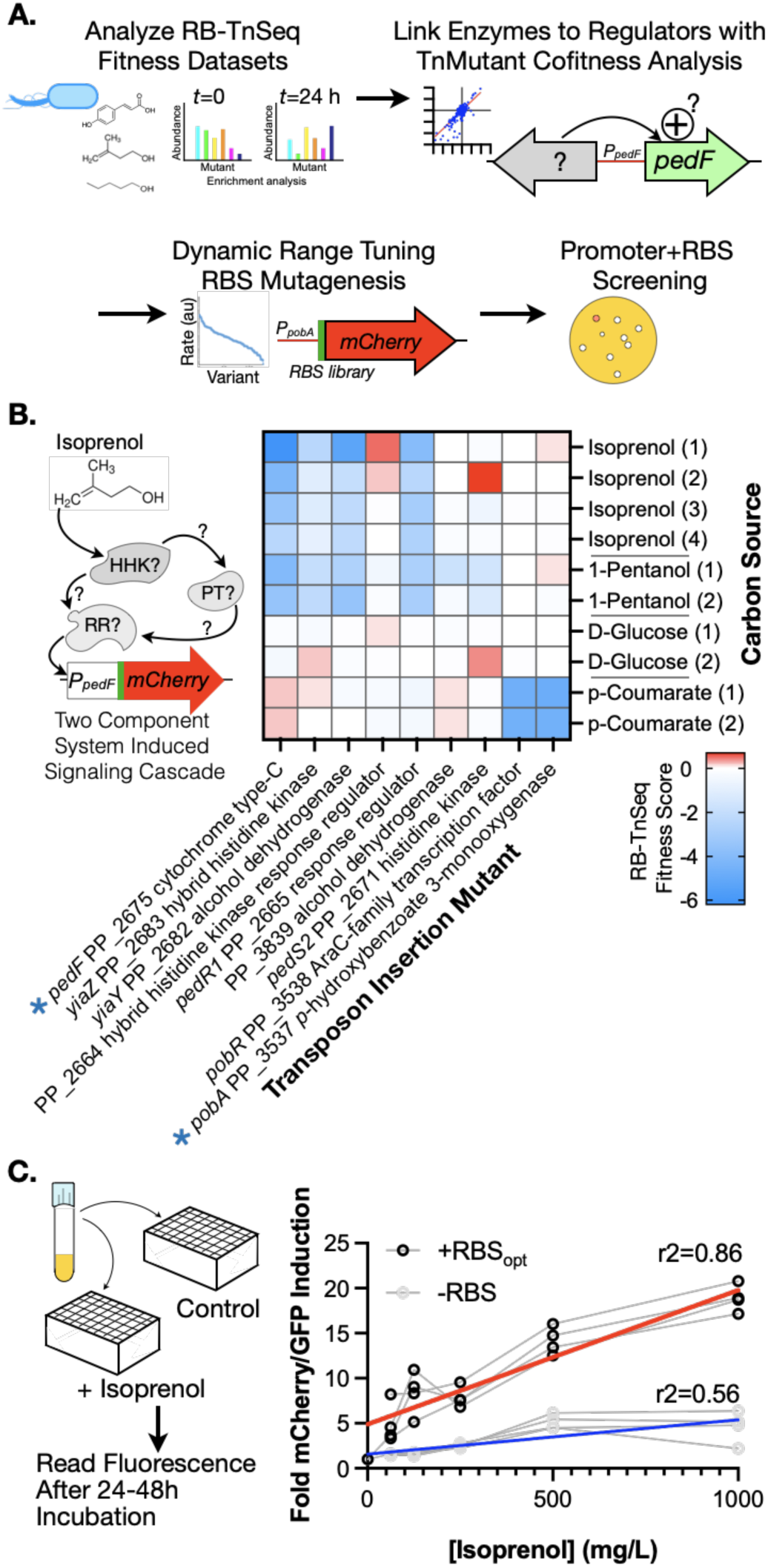
A Workflow for Developing Biosensors Using Functional Genomics and RBS Optimization. **(A)** Biosensor design pipeline sketch. RB-TnSeq fitness data is analyzed for the relevant growth conditions. Co-fitness analysis links genes with similar fitness trends in the selected condition. RBS sequences are computationally predicted and implemented via mutagenesis to optimize the ligand-driven expression of mCherry using the identified promoters in the co-fitness analysis. **(B)** RB-TnSeq fitness heatmap of the selected genes (X axis) from the co-fitness analysis on the indicated carbon sources (Y axis). The number in the parenthesis indicates multiple datasets from the RB-TnSeq library. The asterisk denotes promoter sequences used for biosensor development in this study. Left: proposed signaling cascade that activates the transcription of mCherry at the pedF promoter sequence. **(C)** Characterization of isoprenol linear dynamic range of the designed synthetic biosensors with or without an optimized RBS sequence. A linear regression and r^2^ value is shown. The fold change was calculated by dividing the fluorescent signal from samples grown in the presence of the inducer with control samples grown in the absence of the inducer.

### Elucidating the Isoprenol Biosensor Regulatory Network via Mutant Analysis

Our biosensor helped reveal the native regulatory mechanisms underlying isoprenol catabolism in *P. putida*. By leveraging our rapid, quantitative isoprenol activation assay we compared fold activation between mutants, enabling a facile assessment of which encoded genes were required vs dispensable for *pedF-RBS-mCherry* activation. We used recombineering (**Materials and Methods**) to generate isogenic gene deletions of HHKs and response regulators identified in the co-fit analysis in **Figure 1B**. Knockout strains were transformed with the *P_pedF_-RBS-mCherry* construct to examine differences in the activation of the isoprenol biosensor (**Figure 2A**).

First, we observed that deletion of PP_2665, encoding a response regulator, ablated the biosensor response, indicating this was the primary regulator involved in the isoprenol signaling cascade. Since response regulators work in tandem with upstream sensor histidine kinases, the three most promising candidate histidine kinases evaluated were, PedS2 (or PP_2671), PP_2664 and YiaZ (or PP_2683) due to their co-fitness profiles. Only the Δ*yiaZ* mutant strain showed loss of the biosensor activation, pinpointing the specific HHK required for the isoprenol-induced signaling cascade. Deletion of PP_2664 slightly increased the biosensor fold response compared to a WT control (**Figure 2A**), denoting a possible interaction that attenuates the response to isoprenol. Both YiaZ and PP_2664 encode hybrid histidine kinases, and contain receiver domains. Such HHKs can function directly via a response regulator (e.g. PP_2665) or have additional phosphorelay proteins involved (*40*). In our analysis, PP_2671 was the only other signaling cascade related candidate with a potential role in this cascade, however the ΔPP_2671 mutant did not disrupt the biosensor activation (**Figure 2A**). It is possible that other phopho-relay proteins not revealed in RB-TnSeq co-fitness analysis are involved in the signaling response to isoprenol. From our results, the minimal isoprenol signaling cascade that was able to activate the biosensor included the HHK YiaZ, which either directly or indirectly activates the PP_2665 response regulator, which in turn binds to and drives gene expression at the *pedF* promoter sequence.

Complementation analysis of PP_2665 was used to test genetic linkage with the observed phenotype. Reintroducing PP_2665 in the ΔPP_2665 strain on a plasmid resulted in constitutive *mCherry* fluorescence under all arabinose concentrations tested (**Supplemental Figure 2**). This indicates the construct’s sensitivity to PP_2665 gene stoichiometry for the optimal dynamic range of the system and confirms PP_2665 as a key component in the signaling cascade.

In contrast, transforming the Δ*yiaZ* strain with a plasmid expressing *yiaZ* did not restore biosensor activity (**Figure 2B**). Other *yiaZ* expression constructs were built using the native *yiaYZ* promoter and a constitutive PJ23119 Anderson promoter, but they also failed to complement biosensor activity. Proteomics analysis confirmed that in all plasmid-borne constructs YiaZ was expressed at comparable levels and well above native levels in WT, which was below the detection limit (**Supplemental Figure 3B**). The failure to complement the *yiaZ* deletion with plasmid-based reintroduction was unexpected. However, *yiaZ* is encoded in an overlapping frame with an alcohol dehydrogenase, *yiaY.* We hypothesized that a truncated gene sequence of *yiaZ* could have been subcloned if the annotated start codon was incorrect, likely resulting in the expression of a nonfunctional protein. To address this possibility, we tested the complementation of Δ*yiaZ* with a larger genomic 1.5kb upstream sequence including *yiaY* to eliminate uncertainty regarding the YiaZ coding sequence. This sequence includes PP_2681, *yiaZ* and *yiaY* in a new plasmid construct and now provided successful complementation of Δ*yiaZ*. This indicated that the larger sequence was sufficient to activate the biosensor (**Figure 2B**, **Supplemental Figure 3A**). Inactivating the *yiaY* start codon in the the PP_2681-yiaZ-yiaY DNA sequence failed to complement, indicating the *yiaY* and *yiaZ* gene products were both required for biosensor activity and ruling out any contribution from PP_2681 (**Supplemental Figure 3A**). Since there is an existing genomic copy of *yiaY* in the Δ*yiaZ* strain, tuning of expression levels of *yiaY* and *yiaZ* may be important for the biosensor activation.

To contextualize the unexpected requirement for both YiaY and YiaZ in restoring biosensor activity, we further characterized YiaY. The reciprocal *yiaY* deletion strain showed similar requirements for complementation. While plasmid expression of *yiaY* failed to restore the biosensor activity, the reintroduction of both *yiaY* and *yiaZ* together showed complementation (**Figure 2B**). Domain analysis with PFAM revealed the YiaY protein contains an iron-dependent alcohol dehydrogenase domain. YiaY is 41% identical to an iron-dependent alcohol dehydrogenase from *Zymomonas mobilis* ZM4 (PDB: 3OWO (*41*)). Iron-dependent alcohol dehydrogenases oxidize short chain alcohols to aldehydes, but the product generated by YiaY from isoprenol and the role of an alcohol dehydrogenase in the signaling cascade is uncommon. Bacterial alcohol dehydrogenases form multimeric homocomplexes that participate in multi-layer regulatory functions (reviewed in (*42*, *43*), but are not known to be stoichiometric constituents of two-component signaling systems. Closely related homologs of *yiaY* and *yiaZ* are only detected in 20 *Pseudomonad* genomes (*44*) and no other *genera* (**Supplementary Figure 4, Supplementary Data 4**) indicating functionally similar homologs (*45*) of this HHK and alcohol dehydrogenase operon are phylogenetically restricted.

**Figure 2:**
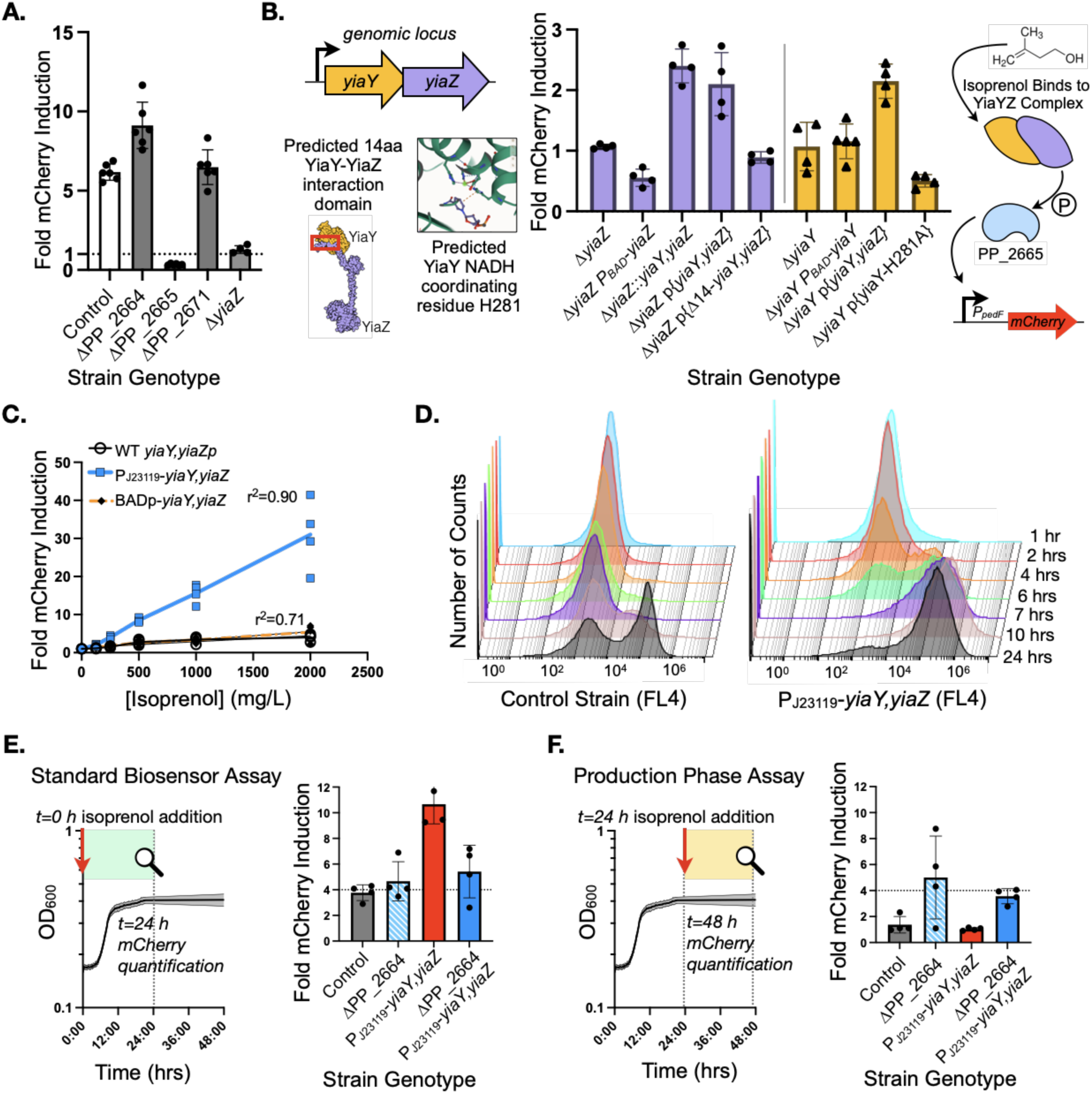
YiaZ/PP_2683, YiaY, and PP_2665 Form the Core Isoprenol Biosensor Signaling Cascade, with PP_2664 as an Inhibitory Modulator. **(A)** Fold mCherry signal detected in deletion stains of the indicated genotypes harboring the *PpedF-RBS-mCherry* plasmid-based isoprenol biosensor in the presence of 1g/L isoprenol compared to the control media without isoprenol. Samples were analyzed for mCherry fluorescence ∼25 hours post inoculation. **(B)** Left: Top. Genomic locus of *yiaZ* and *yiaY* showing their overlapping sequence. Bottom left. AlphaFold (*39*) predicted the physical interaction between YiaY and YiaZ through a 14 amino acid domain. Bottom right. AlphaFold3 analysis of coordinating NADH residues indicated a potential active site of YiaY for residue H281. See Supplementary Figure 5 for structure confidence scores for interacting residues. Middle: complementation assay to restore the lack of function caused by the deletions of yiaZ (purple) and yiaY (yellow). The *yiaY, yiaZ* plasmid contains a 1.5 kb genomic sequence that also includes PP_2681. Right: The known signaling cascade with abbreviated phospho-relay. **(C)** Isoprenol biosensor linear dynamic range in a control strain with the native promoter of *yia* (black), the high-expression constitutive Anderson promoter J23119 (blue) and a minimal BAD promoter with constitutive expression (orange). **(D)** m-Cherry positive population analysis by flow cytometry. The time of sampling is labeled to the right of each axis. 30,000 events (cells) were collected for each timepoint. **(E)** The standard biosensor assay. Isoprenol is added at the same time the strains are inoculated into the fresh media and mCherry is quantified 24 hrs post strain inoculation. **(F)** The production phase biosensor assay. Isoprenol is added to saturated cultures 24 hrs after the strains are inoculated in fresh media. mCherry is quantified 48 hrs post strain inoculation. For all panels, all datapoints are shown and error bars indicate standard deviation from the mean.

YiaY may play a potential structural or enzymatic role in the isoprenol signaling cascade. Neither functions are mutually exclusive. Since biosensor activation was only observed when *yiaY* and *yiaZ* were both genomically or episomally expressed in the *cis* configuration, we hypothesized a model in which both would be constituent subunits of a multimeric complex with a specific stoichiometry for activity (**Figure 2B**). AlphaFold3 multimer analysis (*39*) provided the potential interaction domains between YiaY and YiaZ (**Supplementary Figure 5A**, **Supplementary Data 3**). The leading N-terminal 14 amino acids of YiaY were predicted to interact with the N terminus of YiaZ (**Supplementary Figure 4, 5A)** and truncating these residues from YiaY blocked biosensor activation, but did not destabilize YiaY protein levels (**Figure 2B, Supplementary Figure 3C**). In addition, when YiaY and YiaZ are coexpressed in *E. coli* they co-sediment by gel filtration analysis, consistent with the formation of a protein complex via a direct physical interaction (**Supplemental Figure 6A**). We also used AlphaFold3 ligand analysis (*39*) to determine the potential active site by identifying where the NAD+ cofactor would reside in YiaY, as NAD is generally required for enzyme activity (**Supplementary Figure 5B, Supplementary Data 3**). One of the key residues that could form a coordinating salt bridge with NAD+ in YiaY was mutated (YiaY-H281A) and we noted no biosensor activation in this resulting point mutant (**Figure 2B**). Together, these separation of function mutants indicate that YiaY is an integral accessory factor in the YiaZ/PP_2665 two component signaling pathway, and that the YiaY and YiaZ complex formation as well as its catalytic activity is essential for isoprenol to activate the regulatory cascade.

We reevaluated the isoprenol signaling cascade to identify how the biosensor response could be optimized. With the native *yiaYZ* operon, detection of fluorescent signal required 24-48 hours of incubation for signal maturation and the variation between biological replicates was high (**Figure 1C**, **Figure 2A & 2C**). Constitutive expression of the *yiaYZ* operon may lead to a more rapid isoprenol detection by priming the two key signaling components for activation. We generated a *P*_J23119_-*yiaYZ* or *P_BAD_-yiaYZ* variant strains via recombineering; without introducing *araC* into the *P_BAD_-yiaYZ* strain, the *BAD* promoter should also show constitutive expression. The linear dynamic range of these constructs compared to a WT control strain (**Figure 2C**) showed that the kinetics of detectable fluorescence and linear dynamic range improved in the *P*_J23119_-*yiaYZ* promoter variant. In contrast the *BAD* promoter variant had only a negligible improvement (**Figure 2C**). Differential proteomics analysis of the *P*_J23119_-*yiaYZ* promoter variant identified increased cellular protein expression at multiple gene operons, suggesting the YiaYZ signaling cascade activated a diverse range of processes including many *ped*-family operons (i.e. PP_2663-PP_2680 & PP_4064-PP_4067); glucose uptake proteins and an uncharacterized transporter (KguDTKE and PP_3210); and PQQ synthase proteins (PP_0375-PP_0379) (**Supplemental Data 2-2**).

Flow cytometry of the activated biosensor strains revealed kinetic improvements in signal maturation (**Materials and Methods**). In the P_J23119_-*yiaYZ* promoter variant strain, a distinct bimodal population emerged approximately 4 hours post exogenous isoprenol addition with 25% of the population expressing at least 10-fold more *mCherry* and, by 7 hours, nearly 80% of cells were mCherry positive (**Figure 2D, Supplemental Figure 7**). In contrast, only 20% of the WT control strain population was *mCherry* positive 24-hour post-isoprenol addition, and had a maximum of 5-fold *mCherry* induction (**Supplemental Figure 7**). Overall, the synthetic constitutive Anderson promoter upstream of *yiaYZ* had successfully improved biosensor performance in dynamic range, kinetics, and %CV between biological replicates.

While the biosensor effectively detected isoprenol in log-phase (**Figure 2E**), the signal was lost in stationary-phase when using the WT control or the *P_J23119_-yiaYZ* biosensor strain (**Figure 2F**). However, deleting PP_2664 restored biosensor activity in stationary-phase cultures (**Figure 2F**) suggesting that PP_2664 acts as a competitive inhibitor, potentially forming heterodimers with YiaZ that are less efficient in activating the signaling cascade, especially in stationary phase. The ΔPP_2664 strain, as well as the *P_J23119_-yiaYZ* ΔPP_2664 strain, showed response to isoprenol in stationary phase, although the native promoter showed more variability (**Figure 2F**). The formation and persistence of YiaZ homodimers is likely critical for enabling biosensor activity in stationary phase.

The previous genetic analysis suggested that both YiaY and PP_2664 can modulate YiaZ HHK activity. While histidine kinases are known to form homodimers, the analogous interaction of any histidine kinase with an alcohol dehydrogenase has not been reported elsewhere. We confirmed several biochemical aspects of the YiaY-YiaZ complex. As mentioned earlier, the full length YiaY-YiaZ complex elutes from size exclusion chromatography as a hetero-multimer at a very large apparent mobility (>670 kDa, YiaY monomer: 41 kDa; YiaZ monomer: 61 kDa) of approximately 1:1 molar ratio (**Supplemental Figure 6A**). Expressing the full length YiaZ in *E. coli* is not stable without coexpression with YiaY, but a truncated YiaZ PAS domain protein was successfully purified in *E. coli.* This PAS domain eluted as a homodimer (**Supplemental Figure 6B**), suggesting the full length YiaZ would be competent to behave similarly *in vitro* to form homo-dimers.

We used AlphaFold3 multimer analysis to predict if YiaZ and PP_2664 could interact and observed that the full length YiaZ and PP_2664 proteins could form both heterodimers along contacts between their C terminal response regulator head domains (**Supplementary Figure 5C, Supplementary Data 3**). Moreover, we provide evidence of a physical interaction between YiaZ and PP_2664 using an affinity pulldown & detection by mass spectrometry assay (**Materials and Methods**). An N-terminal in-frame 6HIS epitope tagged allele of PP_2664 was co-expressed with YiaZ in *P. putida* (**Supplementary Figure 6C**). Affinity purification of induced cells showed that YiaZ was enriched by 3.6 fold on nickel beads when 6HIS-PP_2664 was present in the sample over the YiaZ-only control, indicating that these two HHKs could stably interact consistent with the formation of a larger complex. This evidence of protein affinity between PP_2664 and YiaZ supports a potential PP_2664-YiaZ heterodimer as a functional unit in the cell that can exist alongside the PP_2664 and YiaZ homodimers.

Our genetic and biochemical experiments shed insight into the mechanism underlying the isoprenol signaling cascade driving downstream expression of the larger transcriptional response induced by YiaZ and we combine our findings in **Supplementary Figure 8**. In this model, *P. putida* expresses both PP_2664 and YiaZ HHKs under the presence of compatible ligands, which are able to form either homo-or heterodimers at an unknown dynamic rate. YiaY binds to YiaZ and might enzymatically reduce isoprenol to an aldehyde (ie, 3-methyl-3-butenal) and help physically stabilize the modified ligand for recognition by YiaZ. In turn, activated YiaZ autophosphorylation or trans-phosphorylation would cause the subsequent phosphorylation of PP_2665 (its active conformation) and allow it to bind to several genomic loci, facilitating the gene expression in conjunction with a sigma factor. The degree of isoprenol pathway activation in a population is variable as both the monomer protein expression level and ratio of homo-to heterodimer can differ, both contributing to pathway activation heterogeneity (**Supplementary Figure 8**). Since we previously observed that the deletion of *yiaZ* abolishes the biosensor response (**Figure 2A**) we concluded that the PP_2664 homodimer is not activated by isoprenol. In addition, we note that cells restricted to expressing the *yiaZ* homodimer have an improved response to isoprenol (**Figure 2A, 2E & 2F**).

### Alcohol Signaling Regulation is Tuned by HHK Dimer Configuration

We next determined the specificity of this biosensor to other potential ligands, surveying a broader range of short chain alcohols in WT, Δ*yiaZ* and ΔPP_2664 strains. We included these mutants to determine if alcohol activation was modulated by preferential formation of YiaZ or PP_2664 homodimers (**Figure 3**). Using equimolar alcohol concentrations, we observed that branched chain alcohols with the branch in the 1 position (isoprenol and isopentanol) showed strong, 25-fold induction in the ΔPP_2664 strain and was approximately 2-fold stronger than in WT (**Figure 3**). Branched molecules with the branch at the 2 position like 3-methyl-1-pentanol had the opposite preference, showing greater activation in WT over the ΔPP_2664 strain background. Linear alcohols like ethanol and hexanol had variable fold responses ranging from 4-fold up to 40-fold and were preferentially activated in the WT strain background. The branched 2 carbon alcohol, isopropanol, had similar fold activation levels like isoprenol, but with the opposite dimer preference where activation in WT was preferred. In all cases, the absence of YiaZ blocked downstream biosensor activation, indicating PP_2664 homodimers were not able to activate PP_2665 to drive expression of the *P_pedF_-RBS-mCherry* readout with other potential ligands. Additionally, based on YiaY’s CLEAN re-annotation (*46*) as a propanediol oxidoreductase, we confirmed that the biosensor can also be activated by both 1,2-propanediol and 1,5-pentanediol **(Supplemental Figure 9B).** Activation of the sensor by the tested diols showed little preference for either homo or heterodimer forms in the deletion mutants **(Supplemental Figure 9B)**. Outside of these alcohols and diols, amino acids or carbon sources including lysine, succinate, valine, acetate, and glycerol showed no biosensor activation **(Supplemental Figure 9A)**. We include a summary of the WT to ΔPP_2664 response ratio for alcohols as an inset panel in **Figure 3**.

**Figure 3:**
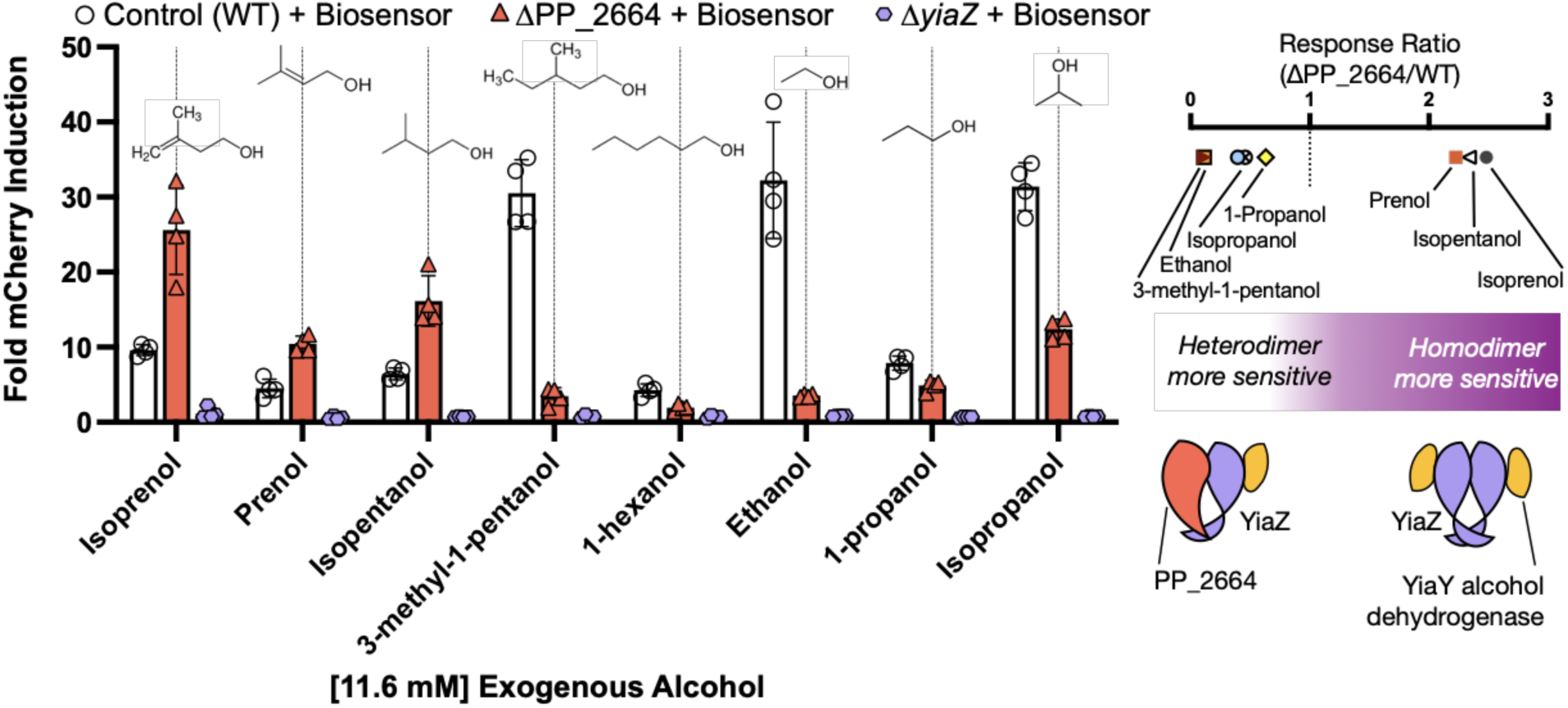
YiaZ and PP_2664 Drive Biosensor Alcohol Specificity in *P. putida*. Left: biosensor fold activation response in the presence of different alcohols. Control with WT *P. putida* strain (white), single ΔPP_2664 knockout mutant (red) and single Δ*yiaZ* knockout mutant (purple) harboring the biosensor. Fluorescent intensity was measured ∼30 hours post addition of the exogenous alcohol. Right: biosensor sensitivity summary sketch to different alcohols depending on the HHK dimer composition. All datapoints are shown and the error bars indicate standard deviation from the mean.

### A Unified Isoprenol Biosensor and Producer Strain

The fluorescent readout from the YiaZ/PP_2665-based *P_pedF_-RBS-mCherry* biosensor was developed using exogenous addition of isoprenol. Next we examined how *P. putida* cells encoding this sensor respond to heterologous isoprenol production, so as to be useful in developing a bioconversion host. Isoprenol production in *P. putida* was previously described (*25*) with a 5-step plasmid-borne pathway (**Figure 4A**). This pathway converts acetyl-CoA into 3-hydroxy-3-methylglutaryl-CoA (HMG-CoA) which is then subsequently metabolized into the mevalonate (MVA) with three additional pathway reactions generating isoprenol. We integrated both the isoprenol pathway and the *P_pedF_-RBS-mCherry* biosensor construct into the genome to bypass the limited availability of antibiotic markers optimized for *P. putida* and to move away from plasmid-borne production systems. As outlined in **Figure 4B**, genomic integration of both modules was attempted with the following strain construction strategy. As a first step, we integrated the biosensor and tested several synthetic promoters to constitutively express either *yiaYZ* and/or PP_2665 for increased biosensor sensitivity. Contrary to our observations testing the plasmid-based biosensor, we found that the best signal to noise isoprenol response was with the constitutive J23100 promoter for the PP_2665, PP_2666 operon. We then replaced the original arabinose inducible *BAD* promoter in the isoprenol pathway with a more economical crystal violet inducible promoter (*47*) and split the pathway across two integration loci as shown in **Figure 4B**. An additional copy of the biosensor was integrated at a second locus to increase sensitivity. Finally, PP_2664 and PP_2675 were deleted to enable stationary phase biosensor function and to block isoprenol catabolism, respectively. At several intermediate stages of strain development we assessed both isoprenol productivity and biosensor sensitivity (**Figure 4C, 4G, Supplemental Figure 10)**. While we observed a linear dynamic response in the lower range of isoprenol concentrations, the signal was generally saturated concentrations above 1 g/L (**Figure 4C**). This genomic strain provided a starting baseline for further isoprenol optimization.

To have a better understanding of this integrated production-biosensor strain (TEAM-2595), we used proteomics to evaluate cellular response under production conditions. Aeration differences from headspace can impact strain productivity and thus we assayed isoprenol titers in both deep-well plates and culture tube formats. TEAM-2595 grown in 5 mL culture tubes produced up to 200 mg/L of isoprenol, whereas the same strain grown in the deep-well plate format only produced approximately 25 mg/L (**Figure 4D**) suggesting cell physiology was favorable for isoprenol production due to aeration differences in the test tube format. Samples were harvested from these two formats to compare differentially expressed proteins at log phase and stationary phase (**Figure 4E**). With a Bonferroni adjusted *p-*value, roughly 2% of detectable proteins showed differential abundance at the 8-, 24-, and 48-hour timepoints, with about twice as many proteins exhibiting significant expression changes at the 24-hour timepoint (71 proteins) compared to the 8-hour timepoint (37 proteins). The proteomics dataset revealed differential expression across a range of cellular functions, including those involved in aromatic catabolism, alcohol dehydrogenases, membrane transporters, and the cold-shock response (**Supplemental Data 2-3**). We also observed differential abundance of several uncharacterized transcriptional regulators including PP_1697, PP_4197 & PP_4100 from the GntR and Cro/Cl family, with no fitness defects by RB-TnSeq analysis on isoprenol-containing M9 medium. While PP_4100 expression was only statistically significant at the 24-hour timepoint, PP_1697 and PP_4197 were significantly overexpressed in the 5 mL format at all timepoints (**Figure 4E**, **Supplemental Data 2-3**). We postulated that overexpressing a global regulator of multiple processes might have a greater impact on isoprenol titer than targeting a single metabolic reaction. To test this, we compared the above uncharacterized global regulators along with PP_2211, a regulator of acyl-CoA synthesis, for overexpression.

**Figure 4:**
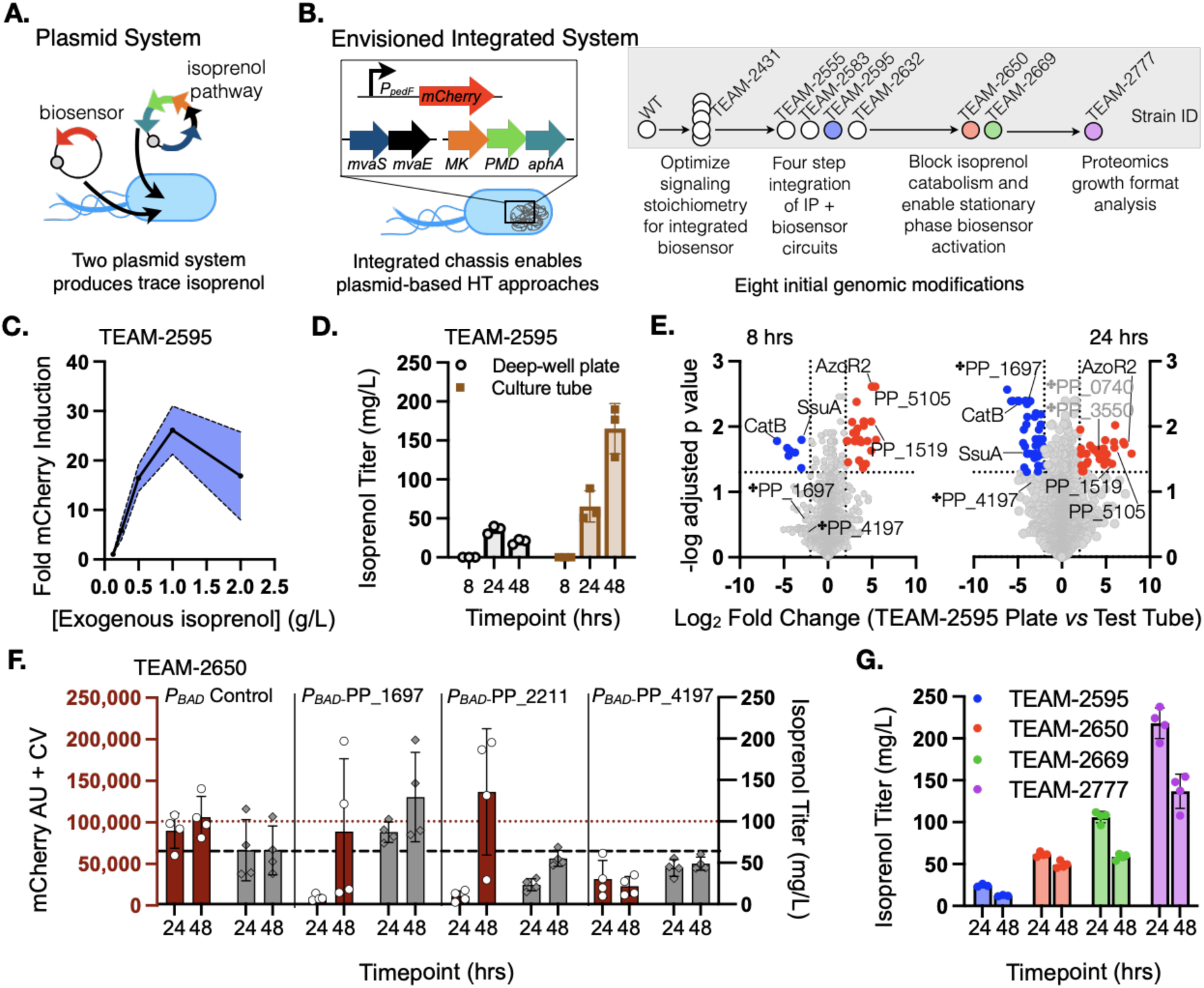
Using an Isoprenol Biosensor to Monitor and Enhance Heterologous Production. **(A)** Sketch of the initial isoprenol production system where the strain harbored the biosensor and the isoprenol production pathway in two independent plasmids. **(B)** Left. Sketch of our proposed isoprenol production system integrating the biosensor and the isoprenol production pathway in different genomic loci. Right. Summary of the strain lineage as genomic modifications were done sequentially. Significant strains that were used and displayed in subsequent figures are color-coded. **(C)** Integrated biosensor fold response to exogenous isoprenol in strain TEAM-2595. The blue shaded area surrounding the solid black line indicates the standard deviation from the mean. n=4 biological replicates. **(D)** Isoprenol productivity of the strain TEAM-2595 in different culture formats, culture tubes (brown) and deep-well plates (grey) at the indicated timepoints. **(E)** Volcano plots representing the ∼3,000 proteins detected by LC-MS/MS in deep-well plate format compared to the test tube format. Statistical significance and fold change cutoffs are marked with dotted lines. Proteins selected for overexpression in F are highlighted with the ✤ symbol. **(F)** Isoprenol productivity (grey) and biosensor fold activation (red) were assayed in deep-well plates using the strain TEAM-2650 harboring several arabinose-inducible overexpression constructs compared to an empty vector control. 0.1% arabinose was used to induce the plasmid-based overexpression constructs. **(G)** Several strains from the strain engineering lineage were assayed for isoprenol production in the deep-well plate format at 24 and 48 hrs. The colors used in this panel correspond to the colors indicated in panel B. In panels D, E, F, and G, all datapoints are shown and the error bar indicates standard deviation from the mean.

Overexpression of the uncharacterized global regulators under *P_BAD_* in the TEAM-2650 was analyzed to compare isoprenol titers directly and *mCherry* fluorescence. For *P_BAD_*-PP_2211 the increased fluorescence over the empty vector control did not correspond to improved isoprenol titers, indicating false positives (**Figure 4F**). *P_BAD_*-PP_4197 overexpression showed good correlation between fluorescence and isoprenol titer, but with an undesirable phenotype: both fluorescence as well as isoprenol titer were dampened. However, the isoprenol biosensor showed both improved fluorescence and isoprenol titer with the *P_BAD_*-PP_1697 construct (**Figure 4F**), indicating a true hit for improved titer in the deep well plate format. We incorporated a constitutive P_J23119_-PP_1697 promoter modification into TEAM-2669, generating TEAM-2777, and this strain increased titers again to 220 mg/L in the deep well plate format, a 7x increase over the initial titer (**Figure 4G**). These experiments demonstrate the complexity of monitoring dynamically produced isoprenol from a biological system compared to measuring a fixed concentration of isoprenol added exogenously into culture media at the start of a time course, and also indicates the potential of using global regulators as drivers for improved isoprenol titer.

### CRISPRi-Based Selection Using a Growth-Coupled Isoprenol Biosensor

One of the major challenges in optimizing heterologous pathways is that their interactions with native metabolism are not yet fully predictable. We used the optimized biosensor system to develop an efficient and unbiased selection method for interrogating if cellular processes were rate-limiting for isoprenol production. We coupled higher concentrations of isoprenol to cell growth by replacing *mCherry* with an *pyrF*, a *URA3* homolog from *Teredinibacter turnerae* T7901 under the *pedF* promoter allowing us to link activation of the biosensor to growth (instead of fluorescence). Via a CRISPRi library, we aimed to select candidate gene knockdowns that improved *pedF*-induced growth as a proxy for higher isoprenol titers (**Figure 5A**).

The newly designed synthetic *pedF-pyrF* construct was sufficient to complement viability in a Δ*pyrF* strain in M9 minimal medium, even without any added isoprenol. This indicated leaky expression was sufficient for growth. In order to identify a retuned RBS sequence that was isoprenol-dose responsive, we mutagenized the *pedF-pyrF* RBS sequence as before (**Materials and Methods**) screening for clones that demonstrated the desired isoprenol-response growth phenotype (**Figure 5B, Supplemental Figure 11**). Two constructs from this screen were selected for validation. The new RBS mutants were reassayed for growth in media containing either 300 mg/L and 1 g/L of isoprenol and identified two optimal RBS sequences, named RBSmut-1 and RBSmut-2, with different isoprenol sensitivity profiles. While the RBSmut-1 had a comparatively lower isoprenol threshold and supported growth in the presence of both concentrations, RBSmut-2 only displayed robust growth in the presence of the higher isoprenol concentration (**Figure 5B**). In addition to the new RBS sequences, the mutant plasmids contained additional modifications in the *pyrF* coding sequence that suggested additional mutations were necessary to reach the desired inducible range (**Supplemental Figure 11**).

We used triparental conjugation to transform a plasmid-born dCpf1/gRNA CRISPRi library in the isoprenol-producing Δ*pyrF* strain (**Materials and Methods)**. The dCpf1/gRNA CRISPRi library contains ∼16,500 gRNAs targeting nearly all genes with 3 unique gRNAs per gene (**Supplemental Data 1-2**). Analysis of the library confirmed expected insert size and distribution (**Supplemental Figure 12, Supplemental Data 1-3**), and that all variants but 40 were detected (**Supplemental Data 1-4**). Clones would show higher fitness for growth if they produced more isoprenol, in turn becoming enriched in abundance by sequencing. As CRISPRi knockdowns may not be fully penetrant (*38*, *48*), the selection for improved growth lessens the risk for false positives. Nonfunctional gRNAs are unlikely to dampen gene expression sufficient to increase isoprenol titers at the thresholds necessary to activate expression of *pyrF*.

**Figure 5:**
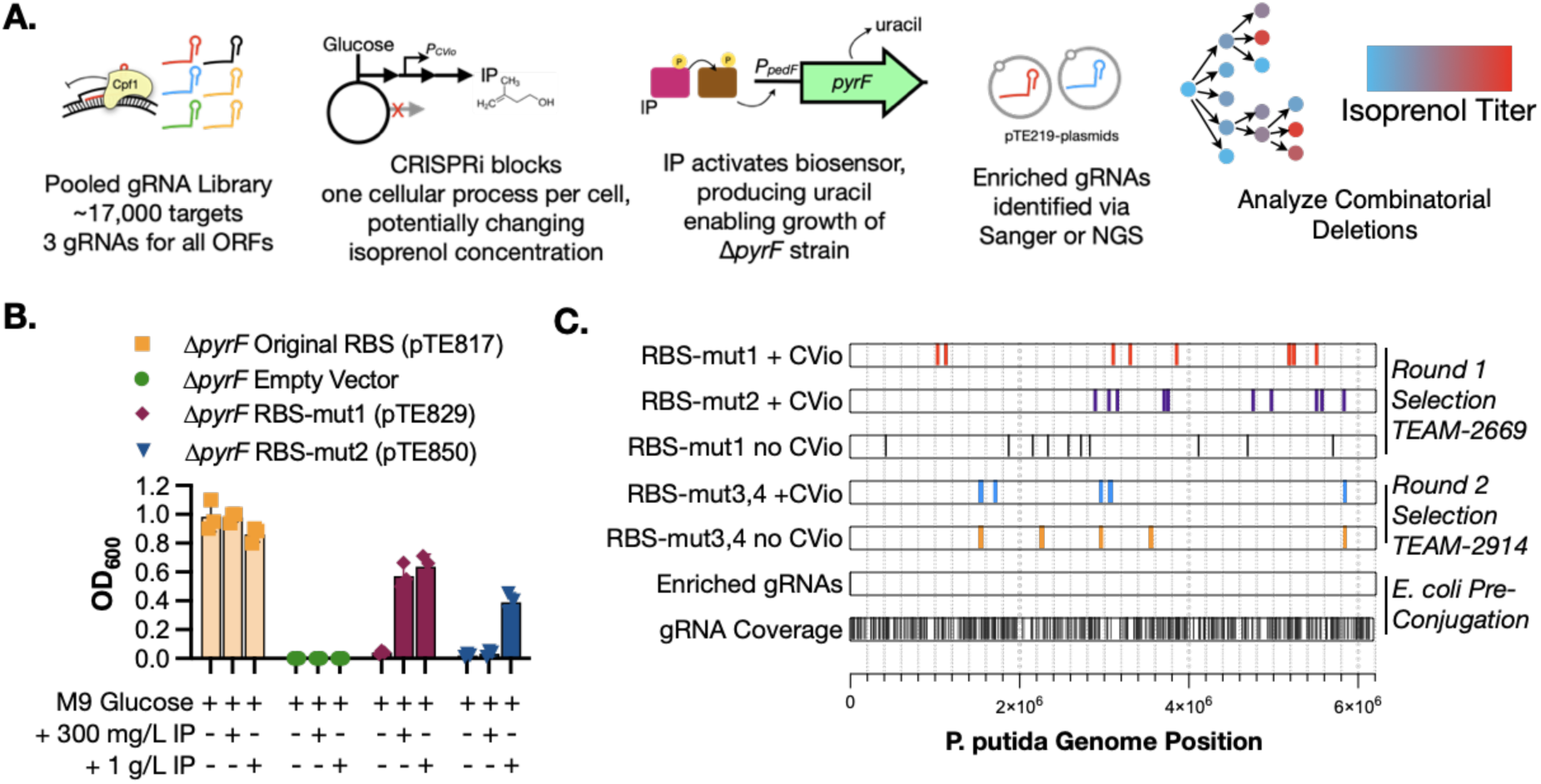
A Growth-Coupled Isoprenol Biosensor for High-Throughput Screening and Targeted Strain Engineering (A) Biosensor repurposing pipeline sketch to enhance heterologous isoprenol production. A pool of ∼16,500 gRNA targets were generated to use a dCas12a/CRISPRi system targeting nearly all genes in *P. putida*. The isoprenol biosensor was modified to replace the pedF-inducible mCherry gene by pyrF to enable the growth on M9 glucose of a ΔpyrF strain in the presence of isoprenol. The *PpedF-RBS-pyrF* Δ*pyrF* strain containing the gRNA library was grown on M9 minimal medium with 2% of glucose and assayed for isoprenol production. Enriched gRNAs at the end of the experiment were amplified, sequenced, and verified by generating the corresponding deletion mutant to role of the deleted gene on isoprenol titer. (B) RBS mutagenesis was performed to optimize the pedF-inducible expression of pyrF to constrain growth to higher concentrations of isoprenol. The response of the construct was evaluated measuring the growth of the strain under different isoprenol concentrations on M9 minimal medium with glucose as the carbon source. All datapoints are shown and the error bars indicate standard deviation from the mean. (C) The enriched gRNA sequencing results were compared between all growth assays performed using multiple optimized RBS and two strains with different isoprenol production productivities. The coverage of the gRNA library as detected with the rapid ONT sequencing method at the start of the experiment was also included. Each line indicates a different enriched gRNA detected from a representative sample and is plotted against its location in the *P. putida* KT2440 genome.

This screen identified pathways that indirectly influenced isoprenol production. The pooled CRISPRi-Δ*pyrF* selection regime was applied in two sequential rounds using four *P_pedF_-RBS-pyrF* variants and four different strains, each with varied base isoprenol titers and isoprenol activation thresholds (**Materials and Methods**). To identify the enriched gRNAs, we amplified the plasmid-borne gRNA sequence and subjected purified amplicons to nanopore transposase-based amplicon sequencing (**Figure 5C**). Overall, 61 gene deletions were selected from these biosensor-linked selection assays for analysis and spanned functional categories including membrane efflux pumps, uncharacterized transcriptional regulators, unrelated metabolic reactions, or had unknown function. It appeared that these pathways indirectly influenced isoprenol production by affecting competing pathways or regulatory networks. Their impact on isoprenol production will be described in the next section.

### Targets from the CRISPRi-Δ*pyrF* Enrichment Screen Guide Strain Engineering

We evaluated the genes identified from the CRISPRi screen on isoprenol production by testing the corresponding loss-of-function deletion mutant (**Figure 6A**, **Supplemental Table 2**). Gene deletions were introduced into the background isoprenol producer strain TEAM-2777 and assayed for isoprenol production in deep well plates (**Materials and Methods**). In the first isoprenol production run comparing 10 single deletion mutants, we identified four different deletion strains that increased titer from 150 mg/L to 250 mg/L (**Supplemental Figure 13**). We next investigated if the identified mutations had synergistic interactions with respect to isoprenol titer. As deletions were chosen from disparate functional categories, we selected the highest producing strain to modify further. We observed both additive, neutral, and inhibitory interactions as deletions were combined (**Figure 6A**).

The targets recovered from this unbiased method had little overlap with known isoprenol-improving reactions from the literature. This method identifies ∼2,000 gRNA reads out of 16,500 unique gRNAs under conditions where a single plasmid may dominate the population. To determine if the absence of known reactions recovered from the screen were false negatives, we selected four known genes from the literature (*25*, *49*) for deletion. If these were not false negatives, these deletions should improve isoprenol titer. Gene targets included the PHA operon (PP_5003-PP_5006/phaAZC-IID) and PP_5007/*phaF* PHA regulator, which generates PHA using Ac-CoA. Additionally, PP_1649/*ldhA*, and PP_3540/*mvaB* were included as hits from genome scale metabolic models. These two genes encode reactions that divert flux away from isoprenol: LdhA converts pyruvate into lactate, and MvaB converts HMG-CoA into Acc-CoA (*25*). We also included a catalase (CatCBA-1) identified from the format-dependent proteomics assay (refer to **Figure 4E**). Of all these rational engineering targets, only the deletion of *mvaB* increased isoprenol production in the strain background tested (**Figure 6A**, **Supplemental Figure 13, 14**), explaining the lack of enrichment for these three gRNAs. Additional rounds of this screen may also reduce false negatives such as the *mvaB* case.

While the CRISPRi screen did not identify *mvaB* as a hit, its deletion proved effective and increased isoprenol titer to ∼500 mg/L in the stacked deletion strain. We chose this specific strain for the second gRNA enrichment analysis starting with generation of the *pedF-RBS-pyrF* with a higher isoprenol activation threshold and subsequent transformation with the pooled gRNA library for selection and sequencing. Analysis of gRNAs by sequencing showed no overlap with the first set, as expected (**Figure 5**, **Supplemental Table 3**). We built more recombineering plasmids and oligos to test more deletion strains as before.

**Figure 6:**
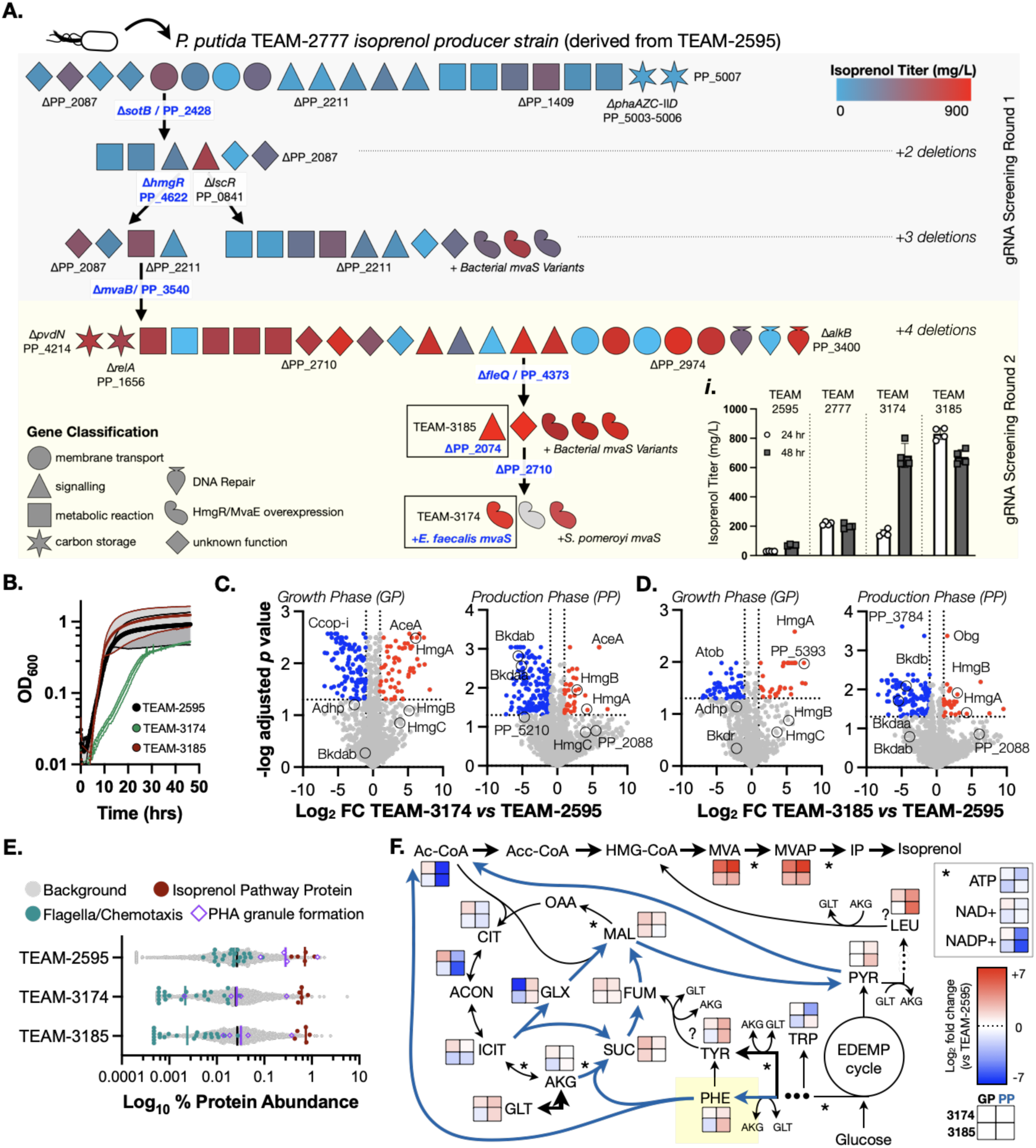
Stepwise Optimization of Isoprenol Production in *P. putida*. **(A)** Strain lineage sketch. This graphic describes the combinatorial modifications evaluated to increase the heterologous production of isoprenol. Over 165 deletions and overexpression constructs were tested and a representative strain lineage is shown. Deletion candidate targets were identified in the dCpf1/gRNA *PpedF-RBS-pyrF* growth assay and isoprenol productivity was evaluated after generating each modification. Candidate modifications were classified based on known gene annotations and are represented with different icons (refer to legend in the bottom left corner). Genes were also color-coded to indicate the isoprenol productivity of such modification (refer to heat-map legend in the top right corner). *i:* Isoprenol production timecourse of selected strains. Production strains were grown in M9 2% glucose and prepared for isoprenol titer analysis and sampled at the indicated timepoints. All datapoints are shown and the error bar indicates standard deviation from the mean. **(B)** Growth curve of two of the best isoprenol producing strains, TEAM-3174 and TEAM-3185, compared to the control strain TEAM-2595 in M9 2% glucose and 1µM crystal violet as inducer. n=4 biological replicates and the shaded area indicates standard deviation from the mean. **(C and D)** Volcano plots representing the ∼2,500 proteins detected by LC-MS/MS in TEAM-3185 (C) and TEAM-3174 (D) compared to the control strain, TEAM-2595. The proteome was analyzed in both growth and production phase. **(E)** Scatter plot indicating the percentage abundance of the selected proteins overlaid on all detected proteins in strains TEAM-2595, TEAM-3174, and TEAM-3185. Selected proteins were grouped according to the indicated functions and are colored according to the legend. **(F)** Metabolic map representing the differential log2 fold abundance change of the indicated metabolites in TEAM-3174 and TEAM-3185 compared to the control strain, TEAM-2595. The squares represent the log2 fold abundance of the metabolites in TEAM-3174 and TEAM-3185 in growth and production phase. Blue arrows indicate the predicted flux to isoprenol based on the variations of the log2 fold metabolite concentrations in production phase samples (24 hour timepoint). Mean values from 3 biological replicates for each sample are reported. A key production phase metabolite, phenylalanine, is highlighted in yellow. Abbreviations: GP: growth phase samples. PP: production phase samples. Pyruvate (PYR), leucine (LEU), tryptophan (TRP), phenylalanine (PHE), tyrosine (TYR), fumarate (FUM), malate (MAL), oxalacetate (OAA), citrate (CIT), aconitate (ACON), isocitrate (ICIT), alpha-ketoglutarate (AKG), glutamate (GLT), succinate (SUC), glyoxylate (GLX), acetyl-coenzyme A (Ac-CoA), acetoacetyl-coenzyme A (Acc-CoA), 3-hydroxy-3-methylglutaryl coenzyme A (HMG-CoA), mevalonate (MVA), mevalonate monophosphate (MVAP), isopentenyl monophosphate (IP), Entner-Doudoroff-Embden-Meyerhof-Parnas cycle (EDEMP).

While single deletions from the second round of gRNA targets enhanced isoprenol titer, combinatorial modifications showed minimal additive effects, suggesting the isoprenol pathway could be rate-limited in activity. MvaS is oftentimes the rate-limiting reaction for isoprenol (or isoprenoids in general) (*24*, *50*, *51*). Switching the NADPH to NADH cofactor used in *mvaS* could alleviate the metabolic cofactor imbalance (*50*). We tested a small set of *mvaS* variants (**Figure 6A**) and identified overexpression of *E. faecalis mvaS* as the most successful for isoprenol production improvement (**Figure 6A**). We concluded the combinatorial strain engineering workflow as observed titers now exceeded 850-900 mg/L in strains, a 5.6x increase from the base strain (TEAM-2777) in isolates TEAM-3185 and TEAM-3174 sampled in two production phase timepoints (**Figure 6Ai**). These titers are comparable to other high producing batch-mode plasmid-borne isoprenol systems in *C. glutamicum* (*24*), *E. coli* (*27*) and *P. putida* (*25*).

Our two highest producer strains TEAM-3185 and TEAM-3174 contain gene deletions in PP_2428, PP_4622, PP_3540/*mvaB* and PP_4373/*fleQ*. TEAM-3174 also overexpresses *mvaS* and ΔPP_2710. TEAM-3185 lacks the mvaS overexpression and ΔPP_2710 but PP_2074 is deleted. See **Table 1** for a summary of putative encoded functions. The overexpression of *mvaS* in TEAM-3174 leads to a kinetic growth delay upon isoprenol pathway induction (**Figure 6B, Supplemental Figure 15A**) that may be related to its ability to produce isoprenol during growth phase, which is not observed in TEAM-3185 nor in the parental TEAM-2595 (**Figure 4D**, **Supplemental Figure 16**). Enhancing isoprenol titer did not sensitize the strains to exogenous isoprenol concentrations relative to the base strain (**Supplemental Figure 15B**). Whole genome sequencing identified 74 additional non-synonymous SNPs of unknown function in both improved producer strains, and a 26.1 kb deletion surrounding the fleQ/PP_4373 locus containing 24 flagella related genes. A comprehensive tabulation of all polymorphisms is described in **Supplemental Data 1-5**.

**Table 1.**
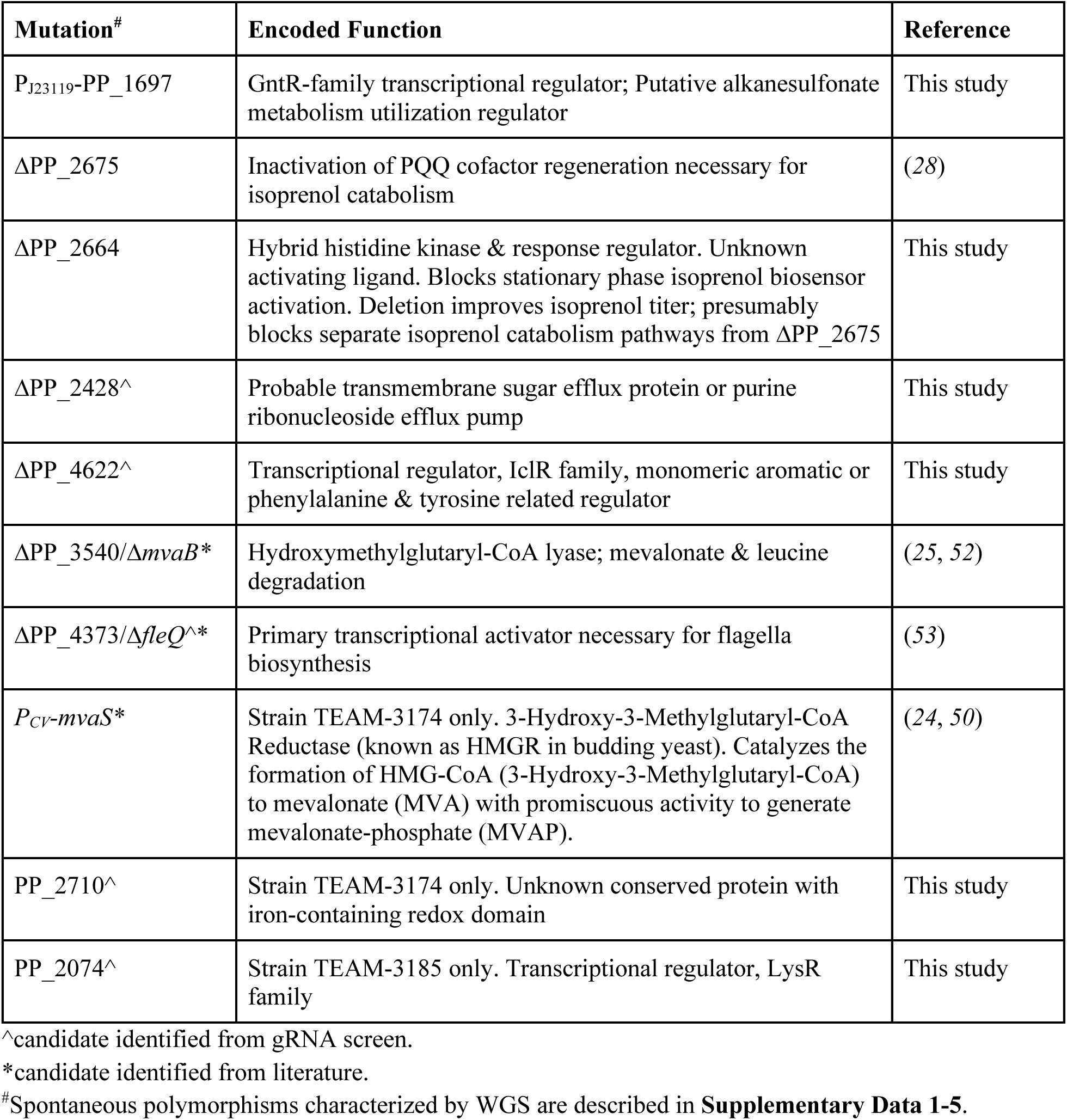
Genes Identified in Isoprenol Production Optimization.

| Mutation<sup>#</sup> | Encoded Function | Reference |
| --- | --- | --- |
| P<sub>J23119</sub>-PP_1697 | GntR-family transcriptional regulator; Putative alkanesulfonate metabolism utilization regulator | This study |
| ΔPP_2675 | Inactivation of PQQ cofactor regeneration necessary for isoprenol catabolism | (28) |
| ΔPP_2664 | Hybrid histidine kinase & response regulator. Unknown activating ligand. Blocks stationary phase isoprenol biosensor activation. Deletion improves isoprenol titer; presumably blocks separate isoprenol catabolism pathways from ΔPP_2675 | This study |
| ΔPP_2428<sup>^</sup> | Probable transmembrane sugar efflux protein or purine ribonucleoside efflux pump | This study |
| ΔPP_4622<sup>^</sup> | Transcriptional regulator, IclR family, monomeric aromatic or phenylalanine & tyrosine related regulator | This study |
| ΔPP_3540/Δ*mvaB*<sup>*</sup> | Hydroxymethylglutaryl-CoA lyase; mevalonate & leucine degradation | (25, 52) |
| ΔPP_4373/Δ*fleQ*<sup>*</sup> | Primary transcriptional activator necessary for flagella biosynthesis | (53) |
| P<sub>CV</sub>-*mvaS*<sup>*</sup> | Strain TEAM-3174 only. 3-Hydroxy-3-Methylglutaryl-CoA Reductase (known as HMGR in budding yeast). Catalyzes the formation of HMG-CoA (3-Hydroxy-3-Methylglutaryl-CoA) to mevalonate (MVA) with promiscuous activity to generate mevalonate-phosphate (MVAP). | (24, 50) |
| PP_2710<sup>^</sup> | Strain TEAM-3174 only. Unknown conserved protein with iron-containing redox domain | This study |
| PP_2074<sup>^</sup> | Strain TEAM-3185 only. Transcriptional regulator, LysR family | This study |

<sup>^</sup>candidate identified from gRNA screen.<sup>\*</sup>candidate identified from literature.<sup>#</sup>Spontaneous polymorphisms characterized by WGS are described in **Supplementary Data 1-5**.**-Omics Analysis Reveals Metabolic Shifts in High Isoprenol Producers**

### Omics Analysis Reveals Metabolic Shifts in High Isoprenol Producers

To characterize the cellular changes in these enhanced isoprenol producer strains, we used an -omics approach with proteomics and metabolomics. We assessed the most optimized final strains TEAM-3174 and TEAM-3185 to the starting strain TEAM-2595 to characterize cellular changes as isoprenol production increased from 50 mg/L to 850 mg/L (**Figure 6A**). We collected parallel samples from growth phase (OD_600_ ∼1) and the 24-hour timepoint in production phase for analysis (**Materials and Methods**). While there are several aspects of the proteomics dataset that merit discussion, the central conclusion from this analysis indicated a broad metabolic shift beyond increasing isoprenol pathway expression or reducing product degradation, hinting instead a new role in amino acid metabolism.

First we examined downregulated isoprenol-degrading reactions which could indicate greater stability of isoprenol (**Figure 6C**, **Figure 6D**). Of the nine alcohol dehydrogenases present in the proteomics dataset, PP_5210, PP_3839/AdhP, and PP_1816, were downregulated, suggesting the remaining five were irrelevant to isoprenol. The other isoprenol degradative steps are shared with leucine catabolism and we broadened our search to amino acid metabolism. This was supported by many enriched amino acid proteins from a ShinyGO analysis (*54*) of GO terms (**Supplemental Figure 17**, **Supplementary Data 1-6**). Specifically, an *hmgABC* operon (PP_4621, PP_4620, and PP_4619) was upregulated. This change in regulation was likely due to the targeted deletion of *hmgR*/PP_4622, an adjacent transcriptional regulator adjacent to the *hmgABC* operon. PP_4621 is a homogentisate deoxygenase implicated in the degradation of phenylalanine by RB-TnSeq mutant analysis (*55*), indicating a relationship between increased phenylalanine degradation and improved isoprenol titers. Next, as amino acids can be used for both carbon and nitrogen sources, we observed that a gene implicated in nitrogen utilization was upregulated. This gene (PP_2088, SigX, RNA polymerase sigma-70 factor) was upregulated during the production phase (34-fold in TEAM-3185 and 20-fold in TEAM-3174 (**Figure 6C**, **Figure 6D**), suggesting a nitrogen starvation response (*55*). While the fold change did not meet the higher adjusted *p-*value threshold for statistical significance, it provided corroborating biological evidence that the ammonium sulfate was rate-limiting in the high isoprenol producer strains, and that the activation of amino acid catabolism pathways could be used as the nitrogen source. Consistent with this idea, we see high (but unchanged) protein expression of PP_3511/IlvE, a branched-chain amino acid aminotransferase that catalyzes the release of glutamate (a common nitrogen source) from leucine and 2-ketoglutarate. Conversely, the downregulation of BKDC proteins (PP_4401, PP_4403) suggests that the formation of acyl-CoA from leucine (or other branch chain amino acids) for carbon utilization is reduced.

Additional proteomics analysis ruled out that the isoprenol pathway expression was rate limiting as previously hypothesized. When proteomics samples were visualized as a fraction of the total proteome for each sample, the abundance of isoprenol pathway proteins was unchanged despite producing 16-fold more isoprenol (refer to **Figure 6ai**), even when overexpression of *mvaS* led to a modest 1.4x protein increase in TEAM-3174 (**Figure 6E**). This indicates that pathway expression is not the rate-limiting step for isoprenol production, and is corroborated by the consistently high metabolite concentrations for mevalonate and mevalonate monophosphate (**Figure 6F**) described in further detail in the next section. We further confirmed this observation by increasing pathway expression with the plasmid-based isoprenol pathway; however, the additional plasmid expression decreased isoprenol titers, even though pathway protein abundance was increased (**Supplemental Figure 18**). These results indicate plasmid burden was detrimental rather than beneficial for isoprenol production.

PHA granule-related proteins also exhibited a significant decrease in abundance in both high-producing strains compared to the control strain (**Figure 6E**). However, we did not follow this target due to our earlier results showing that they did not increase isoprenol production but only variability (**Supplemental Figure 13**) in contrast to plasmid-borne systems (*25*, *49*). However, differences in both amino acid and PHA activity suggested a shift in carbon flux, emphasizing the importance of directly examining metabolite concentrations using metabolomics analysis on the samples collected in parallel.

The key finding from the metabolomics analysis was that leucine, phenylalanine, and tyrosine accumulated during the production phase (and not growth phase) in the enhanced isoprenol producer strains. We calculated relative metabolite concentrations for both growth and production phase and visualized them using a metabolic map (**Figure 6F**). While there were some striking changes with regards to the activation of the TCA glyoxylate shunt in production phase, we were drawn to changes in amino acid concentrations corroborating the enriched amino acid pathways from proteomics as an attractive explanation for the metabolic changes we observed.

Leucine, phenylalanine and tryptophan are all derived from the pentose phosphate pathway (PPP) through the chorismate biosynthesis pathway (*56*). We observed a 2-fold increase in phenylalanine and 15 fold increase in leucine concentrations comparing TEAM-3174 and TEAM-3185 to the base strain (**Figure 6F, Supplemental Data 1-1**). In contrast, tryptophan concentrations showed inconsistent changes between the two producers, as TEAM-3185 had no change while TEAM-3174 showed a 7x decrease (**Figure 6F, Supplemental Data 1-1**). *P. putida* encodes both known catabolic routes for phenylalanine: it can both utilize a CoA-thioester intermediates (*57*) to generate succinyl-coA and acetyl-coA (the *pheBA* gene cluster), as well as the more conventional homogentisate route (*58*) (the *paa* gene cluster) to release fumarate and acetoacetate. Fitness data for *P. putida* grown on phenylalanine indicates both pathways are not functionally redundant; mutations in either pathway is sufficient to compromise fitness on phenylalanine as a carbon source (*35*). Additionally, the proteomics data indicated an increase in the homogentisate pathway proteins. Though most of the Paa operon is not detectable by this proteomics method, the Paa transcriptional regulator, PaaX, was upregulated 8-9x in both strains, suggesting the CoA-thioester pathway is also active. These results indicate that phenylalanine is likely being routed into the TCA cycle with potentially either or both enzymatic pathways. Either leucine and phenylalanine could be used as the replacement nitrogen or carbon source after ammonium sulphate was depleted, consistent with PP_2088 (Sigma-70 Factor, SigX) activation and the nitrogen starvation response identified by proteomics.

We combined our proteomics and metabolomics observations into a testable model: amino acid pools enable specific metabolic states that boost isoprenol production. Specific amino acids could be used as a nitrogen source for growth versus a carbon source for production. Depending on which growth phase the supplement was added, we might see changes to isoprenol titer. In this model, phenylalanine could act as a carbon source used to generate isoprenol after glucose depletion. Leucine, on the other hand, would serve as the nitrogen source to primarily support cell growth. We tested this hypothesis by supplementing the media with phenylalanine or leucine at different points in cultivation - at the start of the timecourse or at the 24-hour timepoint (when glucose was depleted) and monitored isoprenol titer afterwards. The results strongly supported our model. Leucine added at the start of the production timecourse increased isoprenol titer by 10%, but had no impact on isoprenol when added after glucose exhaustion in production phase (the 24 hour timepoint) (**Supplemental Figure 19**). Above 3mM, leucine supplementation reduced titers, indicating a sensitivity to supplied amounts. In contrast, phenylalanine supplementation added at the start of the timecourse did not change isoprenol titers; in contrast, when it was added after glucose depletion, isoprenol titers increased by 15% in a dose dependent manner (**Supplemental Figure 19**). This change in isoprenol titers confirms a specific metabolic state is required for phenylalanine to enhance isoprenol titers, and leucine supplementation is a consequence of increased biomass formation, with no impact when added after glucose is exhausted and primarily acting as a nitrogen source. The decreased isoprenol titers upon initial phenylalanine supplementation indicate a more complex interaction, possibly influenced by the metabolic state of the cells as they transition into production phase. These media amendments had no impact on titer in the base TEAM-2595 strain (**Supplemental Figure 19**). In summary, the media amendment experiment suggests a new strategy for improving production phase titers by revealing carbon and nitrogen sinks that can be targeted for additional metabolic engineering to improve isoprenol production.

### Technoeconomic Parameters for Isoprenol Scaleup

The state-of-the-art *P. putida* isoprenol chassis generated here provides a baseline for future improvement towards economic viability. This study addresses factors that complicate near-term scale-up, but are often overlooked in idealized “*n*^th^ plant” technoeconomic analyses (TEAs). For example, removing the need for antibiotics relieves a barrier to industrial scaleup both in raw reagent cost and the environmental impact (*59*), as government permitting and wastewater disposal can have separate challenges. To better understand the impacts of titer, rates, yields, and amino acid addition on costs we completed a TEA based on a lignocellulosic feedstock (forage sorghum) conversion to isoprenol. This updated analysis indicates a baseline selling price of isoprenol of $31.9/L-gasoline equivalent (LGE), solely using glucose at the yield demonstrated in this study (**Figure 7**). If *P. putida* is engineered to co-utilize xylose at a yield equal to 90% of the demonstrated yield from glucose, the result would fall to $22.4/LGE. It is worth noting that *P. putida* must still be further optimized to achieve parity with the best results in *E. coli* (*60*).

**Figure 7:**
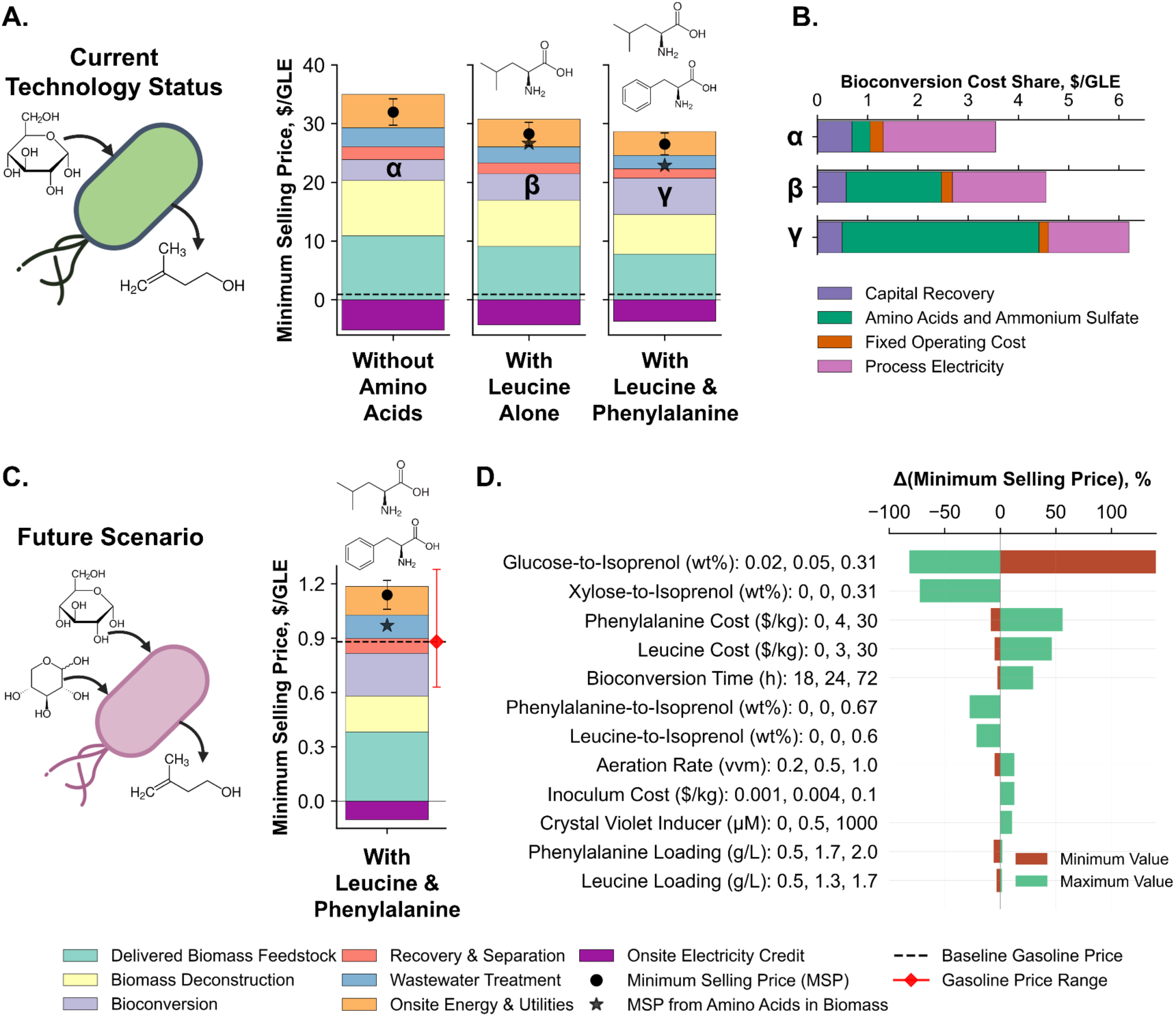
Techno-Economic Analysis of Isoprenol Production in *P. Putida*. **(A)** Minimum selling price of isoprenol with or without amino acid boosters using yield from glucose, as demonstrated in this study. **(B)** Bioconversion stage cost contributions to the total minimum selling price, where α, β, and γ represent bioconversion stages with or without amino acid boosters, as presented in Figure A. **(C)** Minimum selling price at optimal future case considering 90% of the theoretical biomass to isoprenol conversion rate. **(D)** Key parameters influencing the minimum selling price of isoprenol. GLE represents gasoline liter equivalent.

To understand the impact of using amino acid supplements (leucine and phenylalanine) to increase isoprenol titer, we ran a scenario that incorporates the additional input costs and resulting increases in titer (refer to **Supplemental Figure 19**). The results indicate that leucine can reduce the baseline selling price of $31.9/L-LGE (for glucose utilization only) by 11.7%, while leucine and phenylalanine together achieved a net 17.1% reduction in the baseline selling price. Biorefineries can minimize the cost of amino acids by selecting lower purity inputs (in the range of 95–99%) or extracting them from biomass such as seaweed (*61*) or other low-cost biomass/residues (*62*, *63*). For example, selecting a feedstock that can be deconstructed to both sugars and amino acids can eliminate the additional cost of amino acids altogether, leading to a 28.5% reduction in the baseline selling price without amino acids. A sensitivity analysis indicates that, in a large-scale commercial facility, isoprenol yield from all carbon sources, including glucose, xylose, phenylalanine, and leucine, has the greatest impact on minimum selling price (**Figure 7D**). *P. putida* has already been shown to use xylose in other catabolic pathways but the rate of glucose and xylose utilization could vary (*64–67*).

## DISCUSSION

Understanding how microbes perceive and respond to their environment is critical for leveraging their metabolic potential. Our modular workflows leverage functional genomics to discover biosensors, refining their activity with rational approaches. Growth-coupled biosensors can be paired with any mutant library beyond CRISPRi (ie, chemical mutagenesis (*68*) or Tn-Seq (*69*)) to enable rapid scanning of a large search space for a desired phenotype. The isoprenol biosensor in this study (summarized in **Supplemental Figure 8**) recruits a non-cognate pair of sensor HHK (YiaZ) and downstream response regulator (PP_2665). The sensor-response function was modulated with both the alcohol dehydrogenase YiaY and HHK PP_2664, a relatively uncommon configuration but one that effectively enabled the development of a highly responsive isoprenol biosensor in *P. putida* KT2440.

We revealed three key mechanisms that dictate high isoprenol production. First, isoprenol production is an emergent cellular property (*70*) arising from complex, unpredictable interactions. Although individual gene deletions often boosted isoprenol titers, combining deletions did not reliably produce additive effects and sometimes resulted in lower titers (refer to **Figure 6A**). While this property has been reported in other engineered systems (*71*), our panel of systematically generated isogenic mutants offer a unique opportunity to elucidate regulatory network connections relevant to terpenes using the -omics methods described here. Second, during growth phase, our high producer isoprenol strains express all isoprenol pathway enzymes at comparable levels, but only mevalonate (and not isoprenol) accumulates. Only when glucose is exhausted does a “switch” occur, unlocking isoprenol accumulation. It is possible that the accumulation of MVAP beyond a certain concentration threshold causes inhibition of MK and PMD activity during growth phase. Isoprenol production appears to be restored during the production phase after MVAP levels drop below this concentration threshold. This is exemplified with *mvaS* overexpression in TEAM-3174 (refer to **Supplemental Figure 16**). The cost of this phenotype is severe: cell growth is slowed and kinetically delays isoprenol formation in the production phase by 24 hours. Mutants which improve the rate of isoprenol production are rare but valuable in the context of fed-batch production regimes (*72*), but the decreased growth rate for this strain suggest computational methods for rate improvements (*73*) should be compared. Finally, phenylalanine and leucine accumulated in production phase, but only phenylalanine supplementation enhanced isoprenol titers in production phase. This metabolic rewiring favors leucine as a nitrogen source rather than use as a carbon source. Medium amendments do not modulate isoprenol production in the parental strain TEAM-2595, implying metabolic rewiring is specific to genetic modifications generated here and may not be general properties of the amino acids themselves. The underlying change in stationary phase physiology may be related to feedback inhibition (*74*) from higher extracellular concentrations, or perhaps different metabolic profiles as hinted from fitness datasets with comparing leucine as a nitrogen versus carbon source (*55*).

Our strains require further optimization, but have the potential to play an important role in the burgeoning bioeconomy by producing isoprenol as a useful end product and platform chemical for upgrading to fuels and commodity chemicals. We demonstrated production in economical M9 minimal salt media without the use of antibiotics or plasmids with the first genomically integrated design, which sets our strains apart from other microbial isoprenol hosts. These advances in strain engineering lay the groundwork for process-focused improvements required for optimal strain behavior in stirred tank, fed-batch production modes as we aim for cost-competitiveness in fuel and commodity chemical markets.

In conclusion, the use of the refined biosensor in conjunction with our straightforward workflow (RBS tuning & CRISPRi-mediated selection) identified competing cellular functions limiting heterologous isoprenol generation, linking peripheral cellular processes back to central metabolism. With two rounds of screening with different activation thresholds for *pedF-RBS-pyrF,* we increased isoprenol titers stepwise first from ∼25 mg/L to 400 mg/L and then again to 900 mg/L in the second round. Repeating this workflow is trivial to account for other process parameters. For example, utilization of mixed carbon sources containing amino acids under microaerobic conditions (*75*) could benefit from this unbiased optimization strategy. In tandem, machine learning (*76*) or constraint-based metabolic modeling methods (*73*) refined from -omics information (*77*) could suggest modifications that force growth coupling. As all of these methods can generate a near limitless set of predicted modifications, high throughput methods to evaluate designs could use either the growth or fluorescence modality to accelerate strain engineering.

## Supporting information

Menasalvas et al biosensor 2025 supplemental figures

## Abbreviations

HHK: hybrid histidine kinase
RB-TnSeq: Random Barcode Transposon Sequencing
RBS: Ribosome Binding Site
PCR: polymerase chain reaction
WT: wild type
CV: coefficient of variation
dCpf1/dCas12a: deactivated Cfp1/Deactivated Cas12a (CRISPRi system)
gRNA: guide
RNAORF: open reading frame
LC-MS/MS: liquid chromatography tandem Mass Spectrometry
PHA: polyhydroxyalkanoate
SNP: Single Nucleotide Polymorphism
GO: Gene Ontology
ONT: Oxford Nanopore Technologies

## Supplementary Data 1

**Sheet 1**. Metabolite concentrations from selected isoprenol producer strains.

**Sheet 2**. Pooled gRNA targeting sequences synthesized for the library.

**Sheet 3**. Distribution of gRNA pooled library gRNA sequences.

**Sheet 4**. Expected gRNA sequences missing from the library.

**Sheet 5**. Illumina Genome Resequencing and Polymorphism Analysis of Selected Isoprenol Clones.

**Sheet 6.** ShinyGO Enrichment Analysis of High Producer Isoprenol Strains.

## Supplementary Data 2

Proteomics samples. Each individual sheet in this file contains one separate experimental dataset.

**Sheet 1.** Proteomics Analysis of Δ*yiaY* Δ*yiaZ* Complementation by Varied Plasmid Constructs

**Sheet 2.** Analysis of P_J23119_-*yiaY,yiaZ* Constitutive Expression

**Sheet 3.** Proteomics Culture Format Evaluation of Isoprenol Producer Strains

**Sheet 4.** Growth/Production phase Samples of High Isoprenol Producer Strains TEAM-3174 & 3185 Compared to TEAM-2595

**Sheet 5.** Plasmid-born augmentation of Isoprenol pathway overexpression in genomically integrated producer strains

## Supplementary Data 3

AlphaFold3 Multimer analysis of YiaZ + YiaY. AlphaFold3 Ligand Analysis of YiaZ + NAD. AlphaFold3 Multimer analysis of YiaZ + PP_2664.

## Supplementary Data 4

*fast.genomics* analysis of yiaY and yiaZ homolog co-occurrence in microbial genomes.

## Supplementary Data 5

Flow cytometry raw data for mcherry timecourse analysis. Representative Plasmidsaurus ONT gRNA amplicon sequencing reads.

## Acknowledgements

We thank David Carruthers and Aparajitha Srinivasan (LBNL) for technical assistance and constructive feedback. We are deeply appreciative for contributions from Robert Evans (JGI, LBNL), Jan-Fang Cheng (JGI, LBNL) Andria Rodrigues (LBNL), Andy deGiovanni (LBNL), Sudeep Agarwala (Ginkgo Bioworks) and Adam Deutschbauer (EGSB, ENIGMA, LBNL). We thank all members of the Mukhopadhyay group and Jeffrey Cjazka (PNNL) for their constructive feedback and comments on this manuscript. CBORG, a language model-based tool (LBNL Information Technology Division) was used to identify and simplify nested-clause sentence structures and other complex phrasing drafted by the authors. All output was manually reviewed by authors for use in revision. Icons from BioRender were used in the generation of Figure 7. The development and creation of the CRISPRi/gRNA library (Award No. 503809 doi.org/10.46936/10.25585/60001169) conducted by the U.S. Department of Energy Joint Genome Institute (https://ror.org/04xm1d337), a DOE Office of Science User Facility, is supported by the Office of Science of the U.S. Department of Energy operated under Contract No. DE-AC02-05CH11231. This material was also based upon work supported by the Joint BioEnergy Institute, U.S. Department of Energy, Office of Science, Biological and Environmental Research Program under Award Number DE-AC02-05CH11231 with Lawrence Berkeley National Laboratory. The United States Government retains and the publisher, by accepting the article for publication, acknowledges that the United States Government retains a nonexclusive, paid-up, irrevocable, worldwide license to publish or reproduce the published form of this manuscript, or allow others to do so, for United States Government purposes. Any subjective views or opinions that might be expressed in the paper do not necessarily represent the views of the U.S. Department of Energy or the United States Government.

## CRediT Author Contribution Statement

**JM**: Conceptualization, Data curation, Formal analysis, Investigation, Methodology, Validation, Visualization, Writing – original draft, Writing – review & editing **SK**: Data curation, Formal analysis, Investigation, Methodology, Validation **YC**: Data curation, Formal analysis, Methodology, Validation **JWG**: Data curation, Formal analysis, Methodology, Validation **EAT**: Data curation, Formal analysis, Methodology, Validation **NRB**: Formal analysis, Methodology, Validation, Visualization, Writing – original draft, Writing – review & editing **MAA**: Data curation, Investigation, Validation **AR**: Data curation, Investigation, Validation **ISY**: Resources **MG**: Resources **CDS**: Formal analysis, Methodology, Validation, Writing – original draft, Writing - review and editing PDA: Resources, Formal analysis, Validation, Writing - review and editing **TSL**: Resources, Methodology, Writing - review & editing **IKB**: Resources, Methodology, Writing – review & editing **EEKB**: Data curation, Formal analysis, Methodology, Validation, Visualization, Writing - review and editing **CJP**: Data curation, Formal analysis, Methodology, Validation, Supervision, Writing - review and editing **TE**: Conceptualization, Data curation, Formal analysis, Funding acquisition, Investigation, Methodology, Project administration, Resources, Supervision, Validation, Visualization, Writing – original draft, Writing – review & editing **AM**: Conceptualization, Formal analysis, Funding acquisition, Project administration, Supervision, Resources, Visualization, Writing – original draft, Writing – review & editing

## METHODS

### Bacterial growth media and cultivation parameters

*Pseudomonas putida* KT2440 (and its derivative strains) were routinely cultured using standard laboratory conditions described previously (*38*). Bacterial strains were revived from cryostorage by streaking to singles on LB Miller agar plates and maintained for no longer than two weeks. LB Miller (Luria Bertani) medium (10 g/L tryptone, 5 g/L yeast extract, 10 g/L NaCl) was purchased from Becton Dickinson (BD Difco, Product No. 244620). When cells were cultured on petri dishes, LB media was supplemented with 2% (w/v) solid agar (Becton Dickinson, Bacto Agar) and incubated at 30°C. Cryostocks were prepared by diluting bacterial cultures grown to saturation overnight to 25% (w/v) glycerol before storage at -80°C. When strains were tested for isoprenol production, cells were back diluted from LB to M9 media and serially passaged two more times overnight and then induced at the time of back dilution in the third back dilution. At the 1X working concentration, M9 media contains 47.9 mM Na2HPO4, 22 mM KH2PO4, 8.56 mM NaCl, 2 mM MgSO4, 100 μM CaCl2 with 1X trace metals solution (Catalog Num. T1001, Teknova Inc, Hollister CA), 2% glucose, 70 mM (NH4)2SO4 and 30 mM of 3-(N-Morpholino)propanesulfonic acid (MOPS; (Sigma Catalog Num. M1254)) adjusted to a pH of 7.0. This formulation of M9 used for *P. putida* is sometimes referred to as “NREL M9” or “Modified M9” (*79*, *80*). Crystal violet (Sigma-Aldritch, Product No. 61135) was used at a concentration of 1,000 nM (1 µM) to induce production of the integrated isoprenol pathway. The plasmid-borne isoprenol pathway (pIY670) was induced at 0.2% w/v arabinose and 50 µg/mL kanamycin sulfate was added to the media for those specific experiments to maintain the plasmid.

### Nomenclature

When discussing histidine kinases, hybrid histidine kinases, and response regulators in the context of two component signaling systems, such as YiaZ and PP_2665, we use the “/” notation to indicate the functional relationship between the two proteins as is the convention for the field. For all other cases, we use the “/” to indicate common synonyms (usually gene names and gene IDs) used to refer to the same gene or protein in *Pseudomonas putida* KT2440 as annotated in NIH GenBank record AE015451.2.

### Molecular biology

All strains, plasmids, gRNA sequences, and recombineering oligos used in this study are described in **Supplementary Table 4, Supplementary Table 5, and Supplementary Table 6**, respectively. Cloning of synthetic DNA constructs was conducted using chemically competent DH10-β cells purchased from NEB (New England Biolabs (NEB, Ipswitch, MA) or XL-1 Blue (Agilent Technologies, Santa Clara, CA). The open reading frame for *pyrF* homolog *purF* (orotidine 5’phosphate decarboxylase, TERTU_1389, referred to simply as *pyrF*) and 300 bp downstream sequence from *Teredinibacter turnerae* T7901 was synthesized by Genewiz Ltd and assembled into a RSF1010 plasmid backbone immediately downstream of the pedF promoter sequence. Cells prepared for chemical competency were generated with the Inoue method by the UC Berkeley QB3 Core facility (Berkeley, CA). All DNA assemblies were designed using Snapgene (BioMatters Ltd). NEB OneTaq 2x PCR Master Mix was used for routine genotyping as described previously in (*48*) after recombineering, and NEB Q5 2x PCR master mix was used to amplify DNA fragments for isothermal HiFi assembly (NEB). PCR and isothermal assembly were conducted following manufacturer’s guidelines for extension time and annealing temperature. The annealing time was always set to 3 seconds (*81*). Plasmids were transformed into *P. putida* strains of the indicated genotype via electroporation (*82*) after 2 washes in 10% glycerol exactly as described. New Cpf1-gRNA targeting sequences were designed, selecting for gRNA spacer sequences 19-21 nt in length using a 5’-TTTN-3’ PAM sequence. Putative plasmid clones were verified by Sanger sequencing (Azenta Life Sciences, Burlington, MA). Alternatively, complex multi-part DNA assemblies for heterologous gene pathways were verified by whole plasmid sequencing conducted by Plasmidsaurus Inc (South San Francisco, CA). Kanamycin (50 µg/mL) or gentamicin (30 µg/mL) (Teknova Inc, Hollister, CA) was added to the appropriate medium as indicated for experiments requiring the selection of plasmids for both *E. coli* and *P. putida*. Isogenic deletion strains were generated using recombineering as described (*83*) but the induction time with 3-MB addition was increased to 90 minutes for isoprenol producing strains (ie, TEAM-2595) and derivatives. In all cases (specifically for the *yiaY* and *yiaZ) P. putida* genome sequences were inspected to ensure deletion of a targeted ORF did not impact putative coding sequences of adjacent genes and care was taken that the sequences removed would not inadvertently truncate other gene products. Additional genomic integration of *mvaS* homologs included the NADPH-dependent *Enterococcus faecalis mvaS*; a NADH-dependent *mvaS* from *S. pomeroyi* (*24, 50*); and NADPH-dependent *mvaS* from *S. aureus* (*84*). Integration of *mvaS* overexpression constructs and the second copy of the *pedF-RBS-mCherry* constructs was conducted using pSPIN recombinase-based plasmids modified to contain new integration sites and cargo using the protocol as described in (*85*).

### Isoprenol biosensor activity assay: Standard assay and stationary phase assay

Isoprenol induction assays were performed by picking a single colony from a strain transformed with pTE518 into 5 mL test tubes in LB medium supplemented with 30 µg/mL gentamicin 30°C overnight. Saturated cultures were diluted 1:10 into 2 mL of fresh M9 minimal medium with 2% glucose in 24-deep well plates, with or without isoprenol supplementation at the desired concentrations. After 24 and 48 hours, 100 μL samples were taken and transferred to black opaque 96-well plates for fluorescence measurements. mCherry fluorescence was measured at an excitation wavelength of 587 nm and an emission wavelength of 610 nm using a monochromator-based Spectramax M2e Microtiter plate reader (Molecular Devices Inc, San Jose, CA). For stationary phase assays, strains were grown in the 24 well deep well plate format for 24 hours as in the conventional assay described above. Isoprenol was then added to the desired concentrations, the plate resealed, and mCherry fluorescence was measured 24 hours later.

### Flow cytometry

High-throughput flow cytometry experiments were performed using the Accuri C6 flow cytometer equipped with a microtiter plate autosampler (BD). Cells were prepared for isoprenol induction assays and sampled at the indicated timepoints. Upon sampling, cells were diluted to OD600 0.1 in 500 μl of PBS medium. A total of 30,000 events were recorded at a flow rate of 66 μl/min, and a core size of 22 μm. mCherry was excited at 552 nm at 70 mW and emission detected at 610 nm with a 20nm bandbass. Data acquisition was performed as described in the Accuri C6 Sampler User’s Guide and analyzed with Treestar FloJo V10.1. No sample gates were applied during analysis.

### Metabolomics

Organic acids were quantified by liquid chromatography and mass spectrometry (LC-MS) using the method described previously (*86*). All other metabolites were quantified by LC-MS via an Acquity UPLC BEH Amide column (2.1 mm ID, 100 mm length, 1.7 µm stationary phase particle diameter; Waters, MA, USA) and the appropriate guard column (Waters, MA, USA) using exactly as described previously (*87*). Metabolites were quantified via external calibration curves or chemical standards (for relative quantification). Data acquisition was performed via the Agilent MassHunter Workstation software (version 8). Data processing and analysis were performed via Agilent MassHunter Qualitative Analysis (version 6) and Profinder (version 8) software, respectively.

### Shotgun proteomics analysis

TEAM-2595, TEAM-3175, and TEAM-3184 *P. putida* strains were grown for isoprenol production in deep well plates. The log phase samples were harvested when each strain reached an OD600 of 0.7 (roughly 8-14 hours post back dilution and induction) as monitored by a spectrophotometer. The production phase samples were harvested at the 24-hour timepoint. Washed cell pellets for each sample were stored at -80°C until sample preparation. Protein was extracted from cell pellets and tryptic peptides were prepared by following established proteomic sample preparation protocol(*88*). Briefly, cell pellets were resuspended in Qiagen P2 Lysis Buffer (Qiagen Sciences, Germantown, MD, Cat.#19052) for cell lysis. Proteins were precipitated with addition of 1 mM NaCl and 4 x volume acetone, followed by washing the protein precipitant with two additional washes of 80% acetone in water. The recovered protein pellet was homogenized by pipetting mixing with 100 mM ammonium bicarbonate in 20% methanol. Protein concentration was determined by the DC protein assay (BioRad Inc, Hercules, CA). Protein reduction was accomplished using 5 mM tris 2-(carboxyethyl)phosphine (TCEP) for 30 minutes at room temperature, and alkylation was performed with 10 mM iodoacetamide (IAM; final concentration) for 30 minutes at room temperature in the dark. Overnight digestion with trypsin was accomplished with a 1:50 trypsin:total protein ratio. The resulting peptide samples were analyzed on an Agilent 1290 UHPLC system coupled to a Thermo Scientific Orbitrap Exploris 480 mass spectrometer for discovery proteomics.(*89*) Briefly, peptide samples were loaded onto an Ascentis® ES-C18 Column (Sigma–Aldrich, St. Louis, MO) and separated with a 10 minute LC gradient (10% Buffer A (0.1% formic acid (FA) in water) – 35% Buffer B (0.1% FA in acetonitrile)). Eluting peptides were introduced to the mass spectrometer operating in positive-ion mode and were measured in data-independent acquisition (DIA) mode with a duty cycle of 3 survey scans from m/z 380 to m/z 985 and 45 MS2 scans with precursor isolation width of 13.5 m/z to cover the mass range.

For IMAC pull down assay samples, peptides were measured in data-dependent acquisition (DDA) mode with the following parameters: Full MS survey scans were acquired in the range of 300-1200 m/z at 60,000 resolution. The automatic gain control (AGC) target was set at 3e^6^ and the maximum injection time was set to 60 ms. Top 10 multiply charged precursor ions (2–5) were isolated for higher-energy collisional dissociation (HCD) MS/MS using a 1.6 m/z isolation window and were accumulated until they either reached an AGC target value of 1e^5^ or a maximum injection time of 50 ms. MS/MS data were generated with a normalized collision energy (NCE) of 30, at a resolution of 15,000. Upon fragmentation precursor ions were dynamically excluded for 10 s after the first fragmentation event. DIA raw data files were analyzed by an integrated software suite DIA-NN.(*90*) The database used in the DIA-NN search (library-free mode) was the latest Uniprot *P. putida* KT2440 proteome FASTA sequence plus the protein sequences of heterogeneous pathway genes and common proteomic contaminants. DIA-NN determines mass tolerances automatically based on first pass analysis of the samples with automated determination of optimal mass accuracies. The retention time extraction window was determined individually for all MS runs analyzed via the automated optimization procedure implemented in DIA-NN. Protein inference was enabled, and the quantification strategy was set to Robust LC = High Accuracy. Output main DIA-NN reports were filtered with a global FDR = 0.01 on both the precursor level and protein group level. A jupyter notebook written in Python executed label-free quantification (LFQ) data analysis on the DIA-NN peptide quantification report, and the details of the analysis were described in the established protocol.(*91*). DDA raw data files were converted to mgf files and searched against the same database described above with Mascot search engine version 2.3.02 (Matrix Science). The resulting search results were filtered and analyzed by Scaffold v 5.0 (Proteome Software Inc.). The exclusive spectra count of identified proteins were exported for relative quantitative analysis between experimental groups.

### Overexpression and purification of YiaZ/PP_2683 and YiaY/PP_2682

The oligomeric states of the PP_2683 PAS domain (corresponding to amino acids 40-164) and full length YiaY, YiaZ (PP_2682, PP_2683) were analyzed by size exclusion chromatography. The requisite DNA sequences for either protein coding sequence were amplified from *P. putida* genomic DNA and subcloned using HiFi assembly into a modified pET-28b (pSKB3) expression vector to generate a N-terminal 6HIS-TEV-epitope tagged fusion protein constructs under transcriptional control of an inducible T7 promoter and verified by whole plasmid sequencing. Protein expression and purification was conducted essentially as described previously (*92*). Briefly, plasmids were transformed into chemically competent *E. coli* BL21 (DE3) (New England Biolabs) and selected on solid LB agar medium supplemented with 50 µg/mL kanamycin. A single colony was picked to inoculate a 50 mL LB culture supplemented with kanamycin (50 µg/mL) grown overnight. The next morning, saturated culture was diluted 100-fold into a 2L baffled shake flask containing 1 L of LB medium supplemented with 50 μg/mL kanamycin and grown at 37 °C with constant shaking at 190 rpm. When cells reached OD600 of ∼0.7, the culture was induced with 0.5 mM IPTG (final concentration). Approximately 30 min after induction, the cells were transferred to a 18 °C incubator and cells were allowed to grow overnight for ∼18 h at the same shaking speed. Samples were harvested and flash frozen with liquid nitrogen in approximately 20 g wet cell pellet aliquots. When ready for processing, cell pellets were thawed on ice and resuspended in 80 mL Lysis Buffer (25mM HEPES, pH 7.4, 250mM NaCl, 2mM MgCl2, 0.1mg/mL lysozyme, 3ug/mL DNaseI, 1x Protease Inhibitor Cocktail EDTA Free (CalBiochem). Cells were fully lysed by an Avestin Emulsifex Cell Disruptor (ATA Scientific, Caringbah, Australia) following the manufacturer’s protocol. The clarified lysate was obtained after centrifugation at 40,000 x*g* for 45 minutes. Imidizole (20 mM final concentration) was added to the clarified lysate before loading on a FPLC (AKTA Pure) 5 ml His Trap HP column (Cytiva) pre equilibrated with Buffer A (25 mM Hepes (pH 7.5), 250 mM NaCl, 20 mM imidazole, and 1 mM DTT). The column was then washed with five Column Volumes (CV) of buffer A to remove unbound proteins and eluted using a linear gradient formed by mixing buffer A with buffer B which contained 25 mM Hepes (pH 7.5), 250 mM NaCl, 500 mM imidazole, and 1 mM DTT. High purity eluate fractions were combined, treated with TEV protease (1:75 mass ratio, 4°C overnight) to remove the 6HIS tag, and re-fractionated on a second HIStrap column. TEV cleavage was verified by coomassie staining after SDS-PAGE of eluate fractions. The final purified protein from the cleanest fractions were then pooled, concentrated, and analyzed by size exclusion chromatography on a HiLoad 16/600 Superdex 200pg (Cytiva) size exclusion column. Relative protein species size was estimated by overlaying a chromatogram of Gel Filtration standards, (Bio-Rad Inc, Product No. 1511901).

### Co-elution analysis of PP_2664/YiaZ by immobilized metal affinity chromatography (IMAC) - mass spectrometry

To detect potential physical interactions between PP_2664 and YiaZ, we generated two constructs: one construct contained both 6HIS-PP_2664 and *3FLAG*-*yiaZ*, and the second contained only YiaZ. Constructs were generated by HiFi assembly where an in-frame 6HIS epitope tag was placed at the N-terminus upstream of the PP_2664 coding sequence. A *3FLAG-yiaZ* sequence was the second gene in this operon. A similar construct containing only *3FLAG*-*yiaZ* for overexpression was generated for use as a control. All constructs were verified with whole plasmid sequencing as described earlier. Both constructs were transformed into *P. putida* KT2440 via electroporation and a single colony from each transformant was used to inoculate a 25 mL LB culture grown at 30°C supplemented with gentamicin at 30 µg/mL for plasmid selection. The next morning, the culture was backdiluted to a starting OD600 of 0.1 and induced with 0.3% arabinose for 6 hours to express the constructs. Cell pellets with containing roughly 5 OD equivalents were harvested and flash frozen in liquid nitrogen before storage at -80°C. For affinity chromatography, cell pellets were thawed on ice and lysed in 1mL B-PER Buffer (Pierce Scientific) supplemented with 5% glycerol, protease inhibitor (Pierce, Product A32953), 1 mg/mL lysozyme (NEB), and 2.5mg/mL DNase (NEB) for 1 hour with gentle agitation at 30°C. 40 µL of Ni-IMAC MagBeads (ThermoScientific, Product A50589) were prepared for affinity chromatography following the manufacturer’s protocol. 400 µL of crude cell lysate was incubated with MagBeads for 3.5 hours before 3 washes and final elution. The wash buffer contained 1X PBS (Gibco Product 10010049), 1X protease inhibitor, 25 mM imidazole, and 1% Triton-X 100 and the imidazole concentration was raised to 150 mM in the final 2 washes. Proteins were eluted from the collected beads with incubation in 500 mM imidazole for 1 hour at room temperature. Eluate of the same volume was precipitated by acetone and digested with trypsin for shotgun proteomics analysis as described earlier.

### Isoprenol production assays

For each experiment, control strains were revived from cryostorage concurrently with test strains. Overnight LB cultures were prepared in test tubes (5 mL) from single colonies and incubated at 30 °C with 200 rpm orbital shaking and 40% humidity. Strains were adapted to M9 minimal media through two serial transfers. The first adaptation involved inoculating 200 μL of overnight LB culture into 5 mL of M9 minimal media in test tubes. The second adaptation was performed the following day, where 100 μL of saturated culture was inoculated into 1.5 mL of fresh M9 medium in 24-deep well plates. These plates were sealed with a gas-permeable film and incubated at 30°C with 1,000 rpm linear shaking and 70% humidity. After 24 hours, strains were inoculated into production media at a starting OD600 of 0.15. Production runs were performed in 24-deep well plates (1.5 mL) with a gas-permeable film and M9 minimal media supplemented with 1 μM crystal violet (CV) as an inducer. To minimize evaporation, the gas-permeable film was additionally sealed with tape that covered half of each square well. All production runs were performed in quadruplicate, with two biological replicates prepared for each strain when multiple clones were tested. M9 minimal medium was freshly prepared for each production run and discarded when precipitation occurred. When supplemented with amino acids, 100 mM stock solutions of molecular grade chemicals (Sigma) were prepared and filter-sterilized before use. To measure isoprenol titers, samples (200 μL) were collected after 24 and 48 hours and stored at -80°C for analysis. For strains harboring overexpression plasmids, gentamycin (30 µg/mL) was added to maintain plasmid stability, and arabinose was supplemented to a final concentration of 1% (w/v) to induce the isoprenol pathway on pIY670, unless otherwise indicated. Isoprenol titers reported at the 48 hour timepoint post induction were corrected for evaporation in the deep well plate format with additional tape coverage accounting for loss of the authentic standard as previously described (*24*). A similar 43% isoprenol loss was observed for all tested concentrations ranging from 1 g/L down to 62 mg/L. The formula applied for the 48 hour timepoint was ([detected isoprenol 24h] x 0.43) + [detected isoprenol 48h]). Glucose consumption was measured by HPLC exactly as previously described (*48*). Glassware was triple washed with ddH20 and ethanol (*48*) as previously described.

### Quantification of isoprenol by gas chromatography

Isoprenol quantification was performed using the gas chromatography-flame ionization detection (GC-FID 8890, Agilent Technologies, USA) equipped with a DB-WAX column (15 m, 0.32 mm inner diameter, 0.25 μm film thickness, Agilent, USA) as previously described (*15*). Briefly, the GC oven was programmed as follows: 40 °C to 100 °C at 15 °C/min, followed by 100 °C to 230 °C at 40 °C/min, and finally, a hold oven step at 230 °C for 2 min. The inlet temperature was set to 200 °C.Samples harvested for isoprenol analysis were thawed from cold storage on ice mixed with equal volume ethyl acetate (Fisher Scientific, No. E196SK) with 100 mg/L of n-butanol (Sigma Aldrich, No. 281549) as the internal standard as described previously (*93*).

### Optimizing RBS variants for *PpedF-pyrF, PpobR-mCherry*, and PJ23119-PP_1697

Candidate RBS sequences for the *PpedF-pyrF* design were calculated using the de novo DNA RBS calculator (“RBS Calculator Design Mode) available at www.denovodna.com (*94*) and a degenerate oligo with ∼40 potential RBS designs was selected. The degenerate oligo was synthesized with 3’ and 5’ overhangs compatible for ssDNA HiFi assembly as described previously (*83*). Approximately 0.25 µL to 1 µL of the HiFi assembly reaction was directly used for transformation via electroporation into a *P. putida* strain without outgrowth and isolation in *E. coli*. For growth-based selection assays, single colonies were picked into wells of a 96 well microtiter plate and incubated with exogenous isoprenol as indicated when strains contained the isoprenol production pathway, crystal violet was added to the media at a concentration of 1µM. Microtiter dishes were incubated at 30 C for 2-6 days to assess growth under varied isoprenol concentration. Clones with the desired behavior were isolated, plasmid DNA was extracted, and sequenced to identify RBS sequences and any other mutations. For the case of PP_1697 promoter modification, there was no statistical increase in isoprenol titer between RBS-PP_1697 clones screened compared to the RBS-control, and so we used the RBS-design in this study.

### Guide RNA library design, construction and validation

3 guide RNAs targeting upstream of each of the 5591 coding sequences in the *P. putida* KT2440 were designed using gRNASeqRET (*95*) (**Supplemental dataset 1-2**). The 20mer guide sequences were flanked with linkers containing AarI sites yielding 5’ AGAT and 3’ TCAA overhangs, and ordered as a pool of oligonucleotides (Twist Bioscience, CA, USA). The pool was PCR amplified for 9 cycles (Kapa polymerase (Roche)) and Golden Gate assembled into the AarI sites of pTE219_dCpf1_AarI, modified to include *oriT* for downstream conjugation. To help ensure all designed guide species were represented in the library, > 160k (10x the number of variants) *Escherichia coli* colonies were picked. The library was validated by PCR amplification (oligonucleotides 5’-CCACGTGACACACTcacctgc-3’ and 5’-GTTAGCCGTCTCCAcacctgc-3’), and the resultant amplicon library sequenced on the Illumina MiSeq platform, and analyzed using custom pipelines (**Supplemental Dataset 1-4, 1-5**).

### Enrichment of Guide RNAs from Library Under *PpedF-pyrF* Selection

*P. putida* Δ*pyrF* strains transformed with a *PpedF-pyrF* plasmid were subject to triparental conjugation with the gRNA library harbored in *E. coli* DH10 with a *E. coli* pRK2013 *tra+* helper strain. The three strains were spotted onto solid LB agar media and allowed to incubate overnight at 30°C. The next day, a small amount of biomass from the conjugation was isolated with a sterile toothpick and used to innoculate 1.5 mL M9 medium kanamycin with or without 1µM CV in 24-deep well plates with 4 replicates from each conjugation. Samples were grown for 24 hours at which point we examined the cultures for growth. If the cultures showed turbidity or in the best case saturation after 24h post-inoculation, 30 uL of the culture was prepared for colony PCR and the gRNA sequences present on the dCpf1/CRISPRi plasmid were amplified by OneTaq PCR. gRNA amplicons were amplified using oligos TEAM-1174 (5’-gaccagttgcgcctgtcggtgttcagtg-3’) and TEAM-644 (5’-gatcttccccatcggtgatgtcg-3’). Biomass from the LB conjugation spots pre-selection and the *E. coli* DH10 strain harboring the library was also amplified and sequenced to verify the diversity of the initial distribution of gRNAs. gRNAs were sequenced using the Oxford Nanopore linear DNA amplicon service by Plasmidsaurus Inc (South San Francisco, CA). The rapid ONT sequencing service was chosen over other sequencing platforms and providers since it provided gRNA sequencing results as quickly as within 24 hours from sample submission, enabling rapid data analysis for future experimental planning. Raw reads were mapped to the pTE219 reference gRNA plasmid map using Geneious Prime and the aligned gRNA sequences downstream of the 5-’TTTN-3’ PAM sequence were extracted as a CSV file. Targeting spacer sequences were filtered to remove sequences that were 19 bases or fewer. Sequences were compared to the known gRNA targeting sequences (Supplementary Data 4) and implicated genes were selected based on the following criteria: (1) if a particular gRNAs was enriched (>5 reads in one biological replicate) (2) there are multiple gRNAs targeting the same gene (3) gRNAs target genes functionally related (ie, generation of a specific process) or targets in the same operon (4) the repeated occurrence of gRNAs or gene targets across multiple replicates. All selected targets from both rounds are described in **Supplementary Tables 2 and 3**. Verification of gRNA knockdown on isoprenol titers was analyzed in isogenetic deletion strains to both reveal a fully penetrant phenotype and eliminate gene perturbations from potential off-target gRNA repression that would complicate interpretation of changes to isoprenol titers (*96*). Candidate genes from the gRNA enrichment were first grouped by function using HMMer and COG to identify non-redundant cellular processes. At random, we picked several from each category to design new gRNA plasmids and recombineering oligos, choosing 28 targets for the first enrichment screen and 30 for the second screen.

### Computational Structure Predictions

To identify potential interaction domains between PP_2664, YiaY, and YiaZ, we used AlphaFold (*97*), AlphaFill, and AlphaFold3 to model protein structures in the absence of evidence from protein crystallization studies. Protein sequences were identified from Uniprot (PP_2664: Q88JI5; YiaY: Q88JG7; Q88JG6). AlphaFold was run on a LBNL server described in (*98*). AlphaFill was performed using a public webserver as described in (*99*). All structures were reanalyzed in AlphaFold3 (at alphafoldserver.com) as described in (*39*) and prepared for inclusion as figure panels using the ChimeraX software package (*100*).

### Illumina Whole Genome Resequencing and SNP analysis

*P. putida* genomic DNA of the indicated genotypes was prepared for whole genome sequencing and Illumina short read assembly as described previously (*48*). Briefly, cryostocks were revived and struck to single colonies on LB agar medium. A single well formed colony was used to inoculate liquid 5 mL LB cultures and incubated overnight at 30°C (200 rpm shaking) until saturated. Approximately 1.5 mL of cells were harvested, spun down (5,000 x *g*, 3 minutes), flash frozen and lysed (50 U/mL units RNAse, 0.1% w/v SDS, 200mM NaCl, pH 8) for genomic DNA extraction using 1.5 mL of cell culture with a phenol chloroform extraction and Phase Lock sample tubes (Qiagen Sciences, Germantown, MD) to facilitate phase separation. Following isopropanol + NaOAc precipitation, the DNA pellet was washed twice with 70% ethanol and air dried before resuspension. Approximately 2 µg of DNA was used for Illumina sequencing, conducted by SeqCenter Inc. (Pittsburgh, PA). Illumina sequencing libraries were prepared using the tagmentation-based and PCR-based Illumina DNA Prep kit and custom IDT 10bp unique dual indices (UDI) with a target insert size of 280 bp. No additional DNA fragmentation or size selection steps were performed. Illumina sequencing was performed on an Illumina NovaSeq X Plus sequencer in one or more multiplexed shared-flow-cell runs, producing 2x151bp paired-end reads. Demultiplexing, quality control and adapter trimming was performed with bcl-convert (v4.2.4). Variant calling was carried out using BreSeq (*101*) under default settings. Pre-existing mutations in the TEAM-2595 parental strain were not counted as new mutations in the enhanced isoprenol production strains.

### Techno-Economic Analysis

We update our previous biomass sorghum-to-isoprenol production model (*60*), specifically focusing on the bioconversion stage by replacing *E. coli* with *P. putida*. Detailed methods and data sources are outlined in our prior study (*60*). The field-to-isoprenol production process model includes stages for biomass production, deconstruction, bioconversion, recovery and separation, wastewater treatment, and onsite energy and utilities. The biorefinery processes 2,000 bone-dry metric tons of sorghum per day, with 20% moisture content (*60*). Biomass sorghum is received in the form of bales, which is first milled and then pretreated with ionic liquid (5 wt% of cholinium lysinate) at 140 °C for 3 hours (*102*). After pretreatment, an enzyme cocktail (29.4 mg protein/g glucan) is added and hydrolyzed for 72 hours, releasing glucose and xylose (*102*). Following hydrolysis, the solid fraction, primarily lignin, is separated and routed to the boiler for onsite heat and power generation, while the liquid fraction, mainly a sugar solution, is sent to the bioconversion reactor after recovery of ionic liquid. Ionic liquid is recovered using a pervaporation system (*103*).

In the bioconversion reactor, inoculum and nitrogen sources are added. The inoculum is prepared in a seed reactor, where low-cost nitrogen sources, including corn steep liquor and diammonium phosphate, are added (*104*). Ammonium sulfate is supplied to the main bioreactor as a nitrogen source for the microbe (*105*) and 0.5 μM crystal violet is added as an inducer (*47*). We developed three separate models: one without amino acid supplementation, one with leucine alone, and one with a combination of leucine and phenylalanine. The model captures changes in the bioconversion stage and their impact on other isoprenol production stages, including isoprenol recovery, wastewater treatment, and onsite energy and utilities. We assessed the impact of sourcing low-purity amino acids from the commodity market versus those derived from plant biomass.

In the baseline scenario, xylose remains unutilized and is sent to wastewater treatment along with the wastewater. Isoprenol is recovered and separated via a distillation-decantation system (*60*). Wastewater is treated using a combination of anaerobic followed by aerobic treatment processes (*106*). The treated water is used as process water.

Biogas generated from the anaerobic treatment, primarily from remaining sugars, and microbial biomass produced during both the aerobic and anaerobic processes, is routed to the boiler for onsite heat and power generation.The onsite energy generation system produces process steam, which is used to meet the facility’s steam requirements, with any remaining steam used to generate electricity. In cases where biogenic energy sources, including lignin, biogas, and cell mass, are insufficient to generate steam and electricity, natural gas is used in the boiler as a supplemental energy source. The utilities stage includes the makeup process water supply system, cooling tower and cooling water supply unit, chilled water supply unit, and clean-in-place system for providing sterilizing cleaning solutions (*106*). **Supplemental Table 1** summarizes major data inputs used to develop models for different scenarios considered in this study.

We used the standard discounted cash flow rate of return (DCFROR) analysis (*106*) in Excel to determine the minimum selling price of isoprenol based on material and energy balance as well as capital and operating cost data obtained from the process modeling software package-SuperPro Designer V13. We considered an internal rate of return of 10%, plant life of 30 years, annual operating hours of 7,920 hours (24 hours/day and 330 days/year) per year, and income tax rate of 21% (*60*, *104*, *106*). These economic evaluation parameters align with previous techno-economic studies (*60*, *104*, *106*) .

