## Supplementary material for "Biosensor-Driven Strain Engineering Reveals Key Cellular Processes for Maximizing Isoprenol Production in *Pseudomonas putida*": Menasalvas et al biosensor 2025 supplemental figures

**Supplementary Note 1: Initial development of a *p*-CA biosensor with optimized RBS.**
The biosensor pipeline leverages functional genomics information previously collected from diverse growth conditions with a 200,000-member *P. putida* KT2440 RB-TnSeq mutant library (Wetmore et al. 2015). Analysis of transposon-derived gene inactivations with severe fitness defects under specified growth conditions could hint at endogenous inducible regulatory cascades, serving as the basis for inducible biosensors (**Figure 1B**). The robustness of this method is derived from over 200 mutant co-fitness studies curated from diverse growth conditions accumulated from a decade of experimental research, all meeting stringent QC criteria. A co-fitness analysis is a linear regression for all fitness values for a given gene or condition. This correlation determines which other mutants have a similar trend among all tested conditions. Co-fitness values approaching 1 would indicate a perfect correlation between two transposon mutants.

There are several cases where *cis*-acting DNA sequences are activated by simple transcription factors, by either filtering the data for transposon mutants with clear fitness defects with specific conditions, or using known enzymes part of well described catabolic pathways and in turn, their cofit regulators. We also developed a biosensor for *para*-coumarate generalizing this method not just for isoprenol but a second case.

*Para*-coumarate is an aromatic compound derived from lignin (Linger et al. 2014) that can be catabolized by *P. putida* but not other bioconversion hosts like *Escherichia coli* (Rodriguez et al. 2017). The concentration of *p*-CA released from pretreatment varies (Park et al. 2020; Timokhin et al. 2020), and quantification requires analytical methods like HPLC. Developing a biosensor for *p*-CA would facilitate rapid, high throughput characterization of biomass deconstruction methods.

Initially, *p*-CA is catabolized by the enzyme PP_3356/Fcs, producing *p*-coumaryl-coA (Johnson et al. 2017). However, analysis of the upstream DNA sequence revealed no transcriptional regulators with high cofitness values. Based on this analysis the PP_3356 DNA promoter sequence was not an ideal candidate for a biosensor.

In contrast, filtering the RB-Tnseq database for transcriptional regulators with low fitness values for *p*-CA identified PP_3538/*pobR* as a candidate ligand-inducible transcription factor. PP_3538 was strongly cofit with the adjacent gene PP_3537/*pobA*, a *p*-hydroxybenzoate hydroxylase, cofit value ~0.81 involved in the *p*-CA degradation pathway (Entsch and van Berkel 1995). This suggests that PP_3538/PobR autoinduces bidirectional expression from its shared promoter sequence, inducing PP_3537 gene expression in the presence of a compatible ligand. We cloned the promoter sequence upstream of a *mCherry* sequence and tested its expression to exogenously added *p*-CA in the mM concentration range.

Without tuning the RNA secondary structure of the $P_{PP\_3537}$-*mCherry* construct, the initial mCherry fluorescence had low signal to noise, poor linearity, and was not reproducible across day to day replicates. Augmenting our promoter-*mCherry* design with an RBS was key to building a robust biosensor. We screened a small, 40-member library of predicted candidate

RBS variants. One RBS variant enabled a 100-fold increase in dynamic range and also improved the linear dynamic range for the analyte. This linear dynamic range in the 1 g/L concentration is well matched for current *p*-CA concentrations from lignocellulosic biomass. A simpler application of this workflow lacking the RBS optimization step on AraC-family transcriptional regulators that included a *pobR-RFP* biosensor was described in an earlier report (Pearson et al. 2024). The pobR promoter sequence and active site was also modified using degenerate oligonucleotides (and not with the Salis lab RBS calculator tool) for expression *E. coli (Jha et al. 2018)*. Crucially, by excluding the RBS optimization step, several potential ligand-responsive biosensors in the previous study showed a limited or no predicted ligand dose response, indicating room for further analysis.

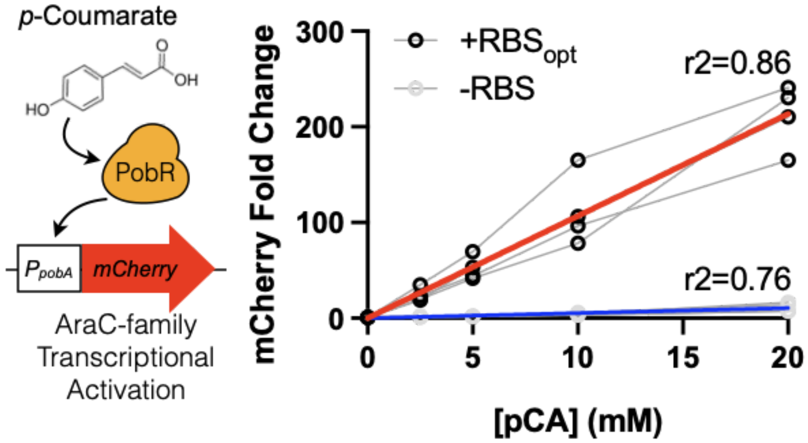


**Supplementary Figure 1. Development of a *p*-CA Biosensor.** Refer to Supplementary Note 1 for additional details on the use of RB-TnSeq data. Linear dynamic range of the optimized *p*-CA biosensor. The optimized *p*-CA biosensor was assayed in M9 minimal media with 2% of glucose and a range of *p*-CA concentrations (0 to 1 g/L). The fluorescent signal was measured 24 hours post *p*-CA addition and compared to a control sample grown in the absence of *p*-CA. A schematic of the proposed single-component signaling system describing how *p*-CA interacts with the signaling cascade is on the left. All datapoints are shown from 4 biological replicates.

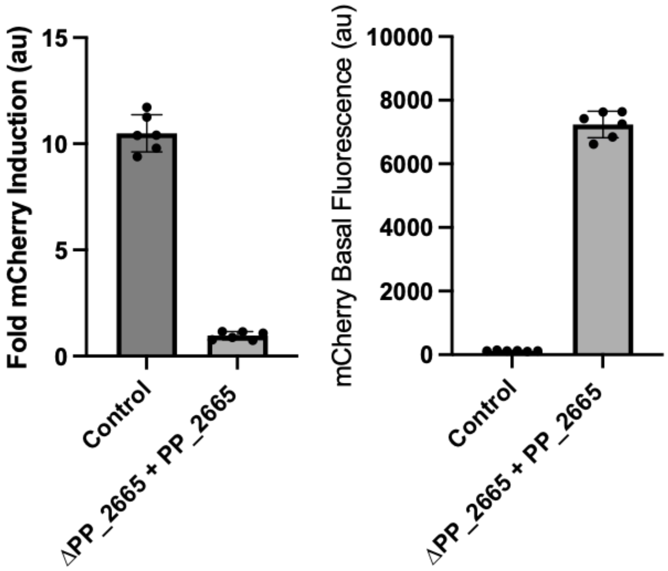


**Supplemental Figure 2. Complementation of ΔPP_2665 with $P_{BAD}$-PP_2665.** WT and mutant *P. putida* ΔPP_2665 strains were transformed with a plasmid expressing PP_2665 under the control of an inducible *BAD* promoter and prepared for an isoprenol biosensor activity assay with 1 g/L isoprenol. High basal mCherry biosensor activation was observed in the ΔPP_2665 $P_{BAD}$-PP_2665 strain. The control strain shows very low basal fluorescence in the absence of isoprenol, but the ΔPP_2665 $P_{BAD}$-PP_2665 shows ~7,000 au mCherry signal with and without exogenous isoprenol addition. All datapoints are shown and the error bar indicates standard deviation from the mean.

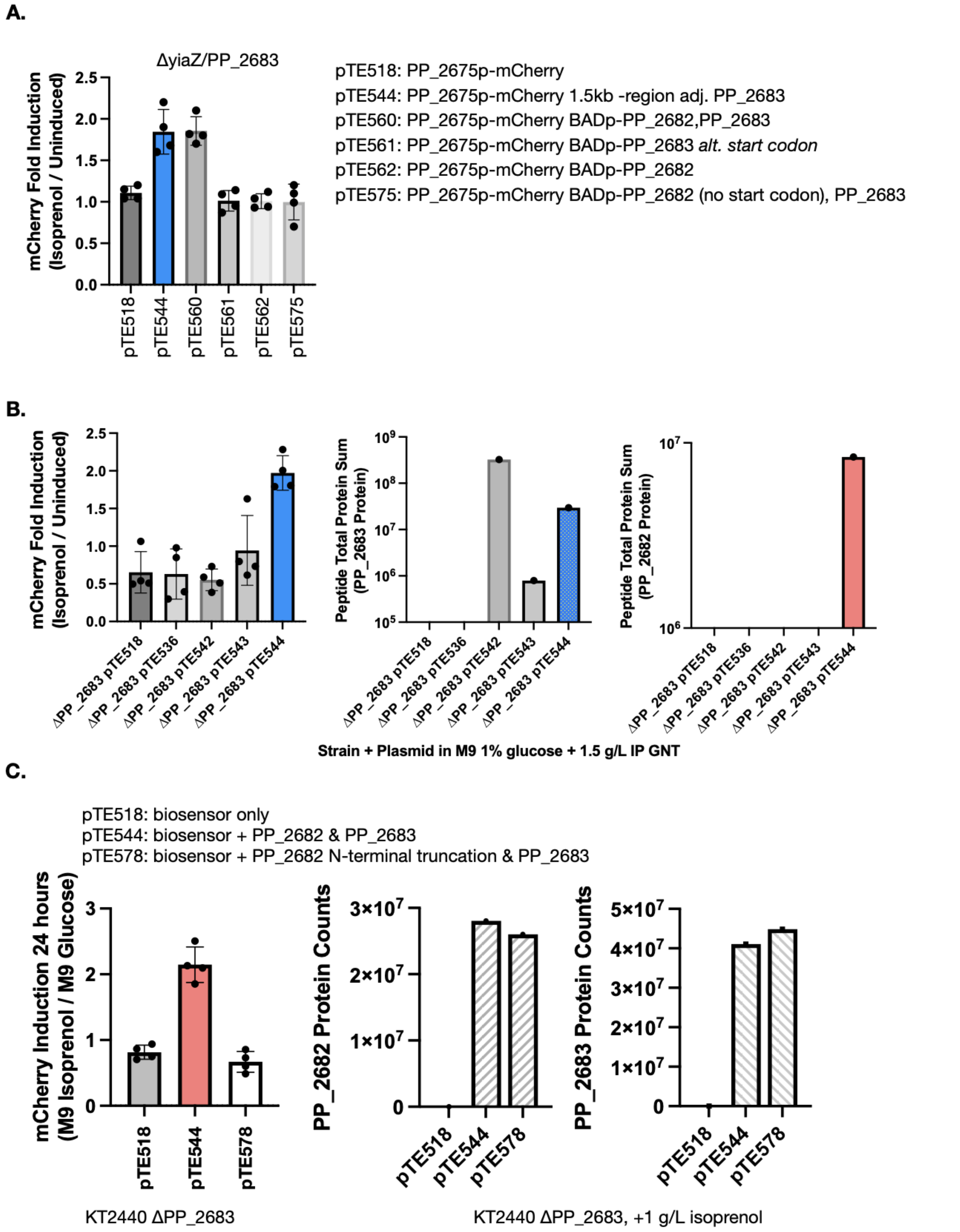


**Supplemental Figure 3. Complementation Analysis of PP_2683/YiaZ. (A)** The effect of different gene constructs on restoring biosensor activity in the Δ*yiaZ* strain. The x-axis shows the different constructs used for complementation: Empty Vector (EV), the 1.5kb region surrounding YiaZ, $P_{BAD}$-PP_2682,PP_2683, $P_{BAD}$-PP_2683 with an earlier start codon, $P_{BAD}$-PP_2682, or

$P_{BAD}$-PP_2682(no start codon),PP_2683. Fold signal from samples with or without isoprenol are plotted after ~24 hours incubation. All plasmids contain the $P_{pedF}$-RBS-mCherry biosensor sequence. **(B)** Proteomics analysis of PP_2683 plasmid constructs from isoprenol biosensor assay with 1.5g/L isoprenol and incubated for 24 hours. These samples were collected from new transformant plasmids on a separate day. pTE536: $P_{PP\_2682}$-PP_2683. pTE542: $P_{BAD}$-PP_2683. pTE543:$P_{J23119}$-PP_2683. pTE544: 1.5kb region genomic region containing PP_2681, yiaY, and yiaZ/PP_2683. All plasmids also contain the $P_{pedF}$-RBS-mCherry biosensor sequence. *mCherry* fluorescence is indicated on the left most panel. Shotgun proteomics counts for PP_2683 and PP_2682 are indicated in the middle and right hand panels. Complementation only occurs when both PP_2682 and PP_2683 are expressed at balanced protein levels. **(C)** Left. Isoprenol biosensor activity assay in ΔyiaZ/ΔPP_2683 strains with a YiaY N terminal truncation mutant. Right, analysis of protein expression levels. pTE578 contains a truncation of the first 14 amino acids at the N terminus of PP_2682 along with a full length *yiaZ/*PP_2683 gene. YiaY/PP_2682 protein expression is comparable to the WT YiaY/PP_2682 in the control complementation plasmid. The failure to complement when protein levels are the same implies a protein interaction is required for functionality. All datapoints are shown and the error bar (if used) indicates standard deviation from the mean.

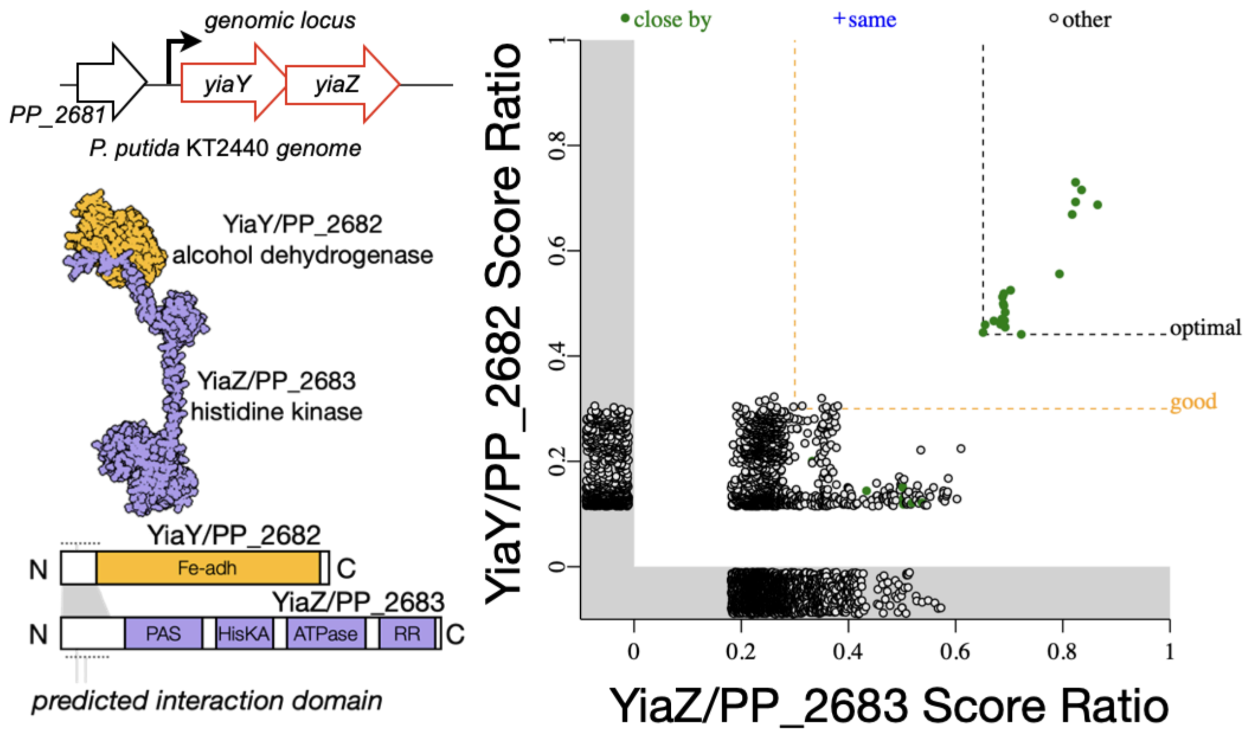


**Supplemental Figure 4. *Fast.genomics* Analysis of YiaY, YiaZ Operon Structure in Prokaryotic Genomes.** YiaY and YiaZ homologs in the same genome are plotted. In other *Pseudomonads*, YiaY is known as ErcA. The X and Y axis are the score ratio for homologs of either YiaY and YiaZ respectively, which is a value that compares the bit score of a protein's homolog to the maximum score. (Refer to (Price and Arkin 2024) for additional methodological details and **Supplemental Data 4** for output genomes). The green shaded circles indicate YiaY and YiaZ homologs that are within 5kb of each other in the analyzed genomes. Open circles are genomes where *yiaY* and *yiaZ* are not adjacent. A schematic on the left hand side of the scatter plot shows the overlapping coding sequences of *yiaY* and *yiaZ* in the *P. putida* KT2440 genome, alphafold multimer predicted interaction domains (also shown in Figure 2B) and predicted PFAM protein domains based on amino acid homology.

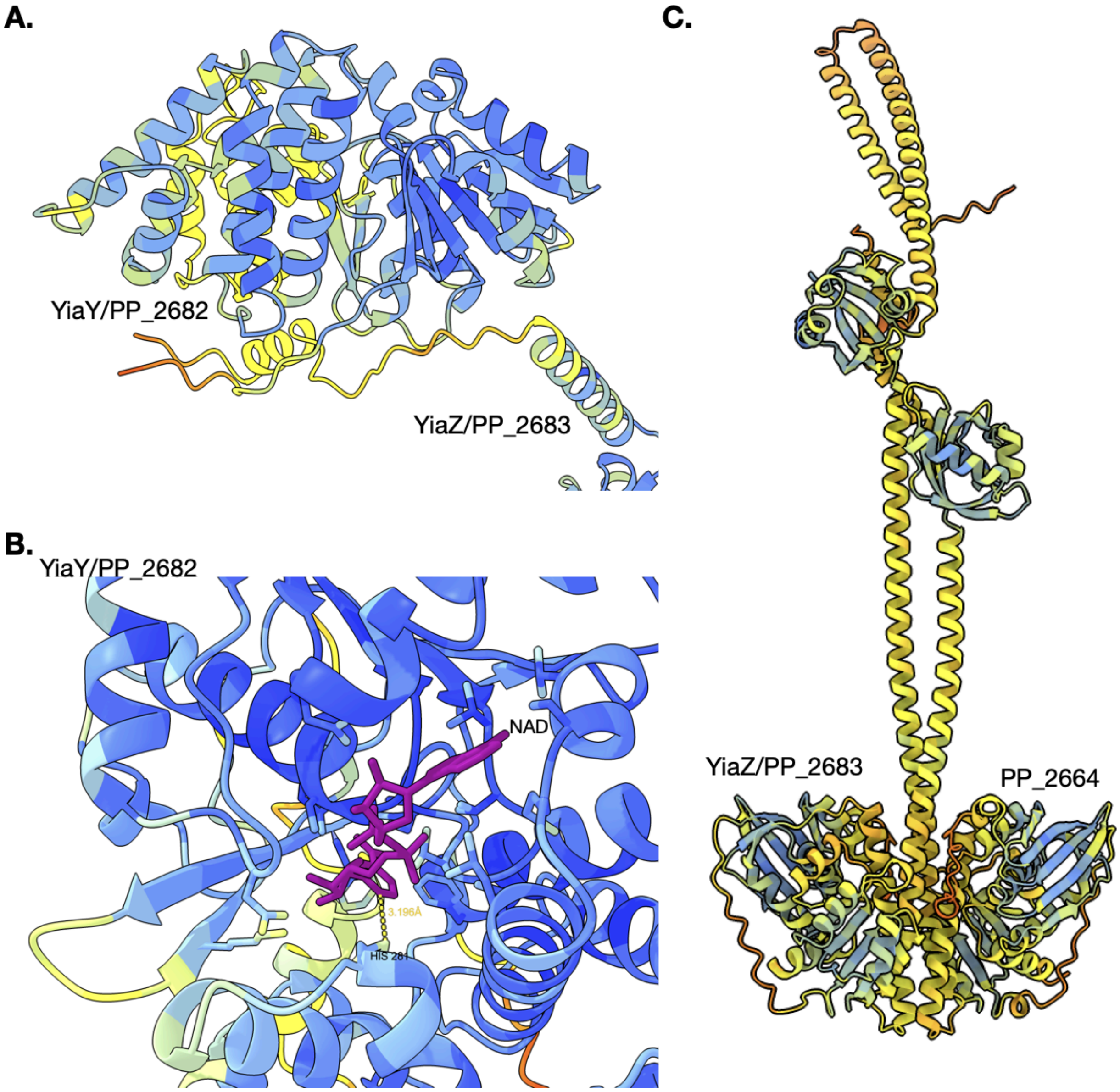


**Supplemental Figure 5. AlphaFold3 Predicted Multimer and NAD Ligand Analysis.** Refer to Supplemental Dataset 3 for full Alphafold3 structure prediction output in .CIF format. Confidence values per residue are supplied as .json files. Up to four other predicted structures are included and here a representative structure is shown. Residue confidence value colors are shown in a gradient displayed using ChimeraX and the output bfactor palette. **(A.)** YiaY and YiaZ Alphafold3 multimer analysis. The interaction domain for YiaY and YiaZ has low to poor confidence as indicated by the yellow to orange colors. **(B.)** Coordinating residues for the NAD cofactor in YiaY. Moderate to highly confident ranked residues surrounding the NAD are indicated by the light blue and dark blue colors. A potential salt bridge between the NAD and H281 with a distance of 3.196 Angstroms is highlighted. **(C.)** Alphafold3 Multimer analysis of YiaZ and PP_2664, showing interaction between their C-terminal response regulator domains at the bottom of the panel.

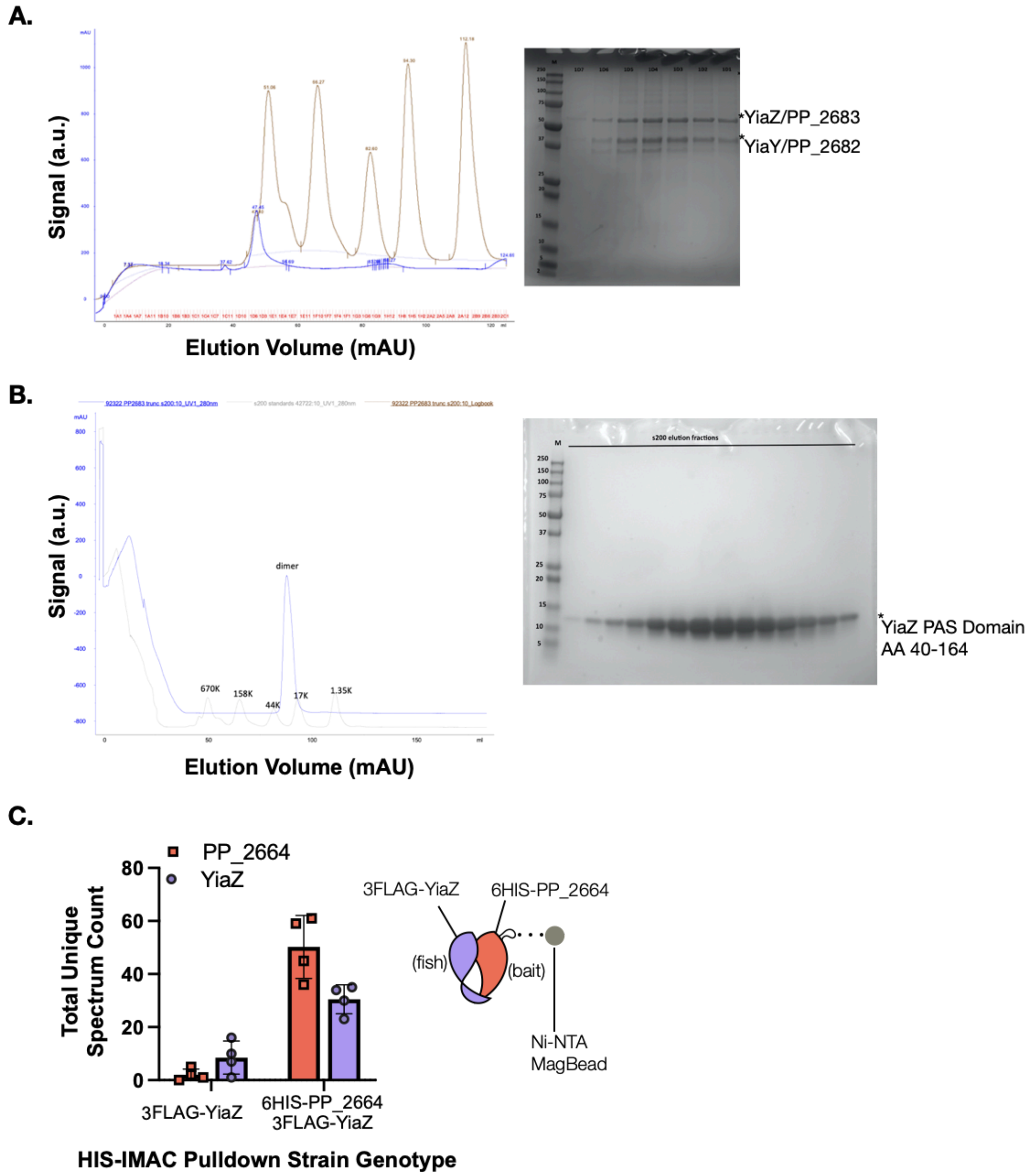


**Supplemental Figure 6. Purification and SEC analysis of YiaY, YiaZ (PP_2682/PP_2683) Co-Expressed in *E. coli* via Ion Exchange Chromatography.** (A) YiaY and YiaZ were co-expressed (pTE710) in *E. coli* and purified by a 6HIS tag and after tag removal, analyzed using ion exchange chromatography (see Materials and Methods). The purified proteins were then subjected to size exclusion chromatography to determine multimeric status. Figure A shows a very large multimer (>670kD) made of a 1:1 molar ratio of YiaZ and YiaY. Elution fractions from ion exchange chromatography are shown in the Coomassie gel on the right, and the relative mobility by size exclusion chromatography are on the left. (B) The YiaZ PAS domain (pTE588) was purified as in A. (C) HIS-IMAC affinity purification of 6HIS-PP_2664 and 3FLAG-YiaZ (pTE908, pTE906, see Materials and Methods). Background binding of 3FLAG-YiaZ eluting with the Ni-NTA beads after wash buffers containing imidazole and Triton-X 100 in the absence of a 6HIS protein is accounted for with a control plasmid expressing only 3FLAG-YiaZ. All datapoints are shown and the error bar indicates standard deviation from the mean.

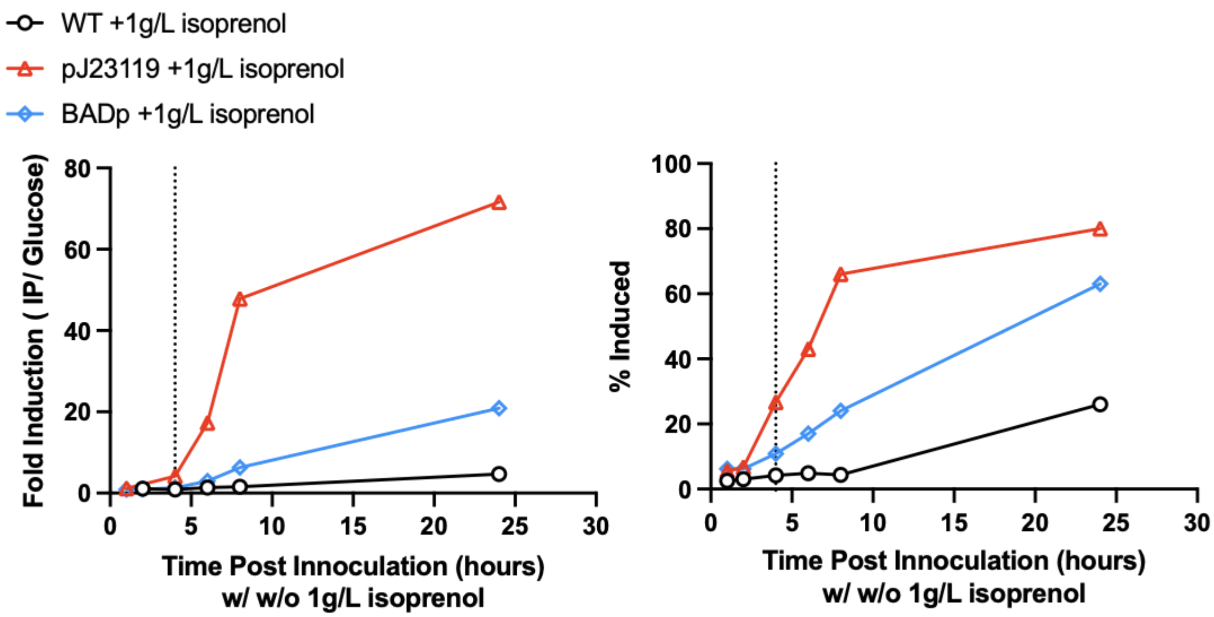


**Supplemental Figure 7. Induction Kinetics and mCherry Population Distributions by Flow Cytometry.** Refer to Figure 2D. Events collected from the 0 hour timepoint for each strain as indicated in the figure legend were used to set mCherry fluorescent thresholds, and the mean fluorescent value of the calculated for the fluorescence from uninduced samples. When bimodal populations were observed, the mean fluorescent value of the second peak with increased fluorescence used to identify the induced population as the induced population. The fluorescent value from the 0 hour timepoint was used as the denominator (uninduced mCherry fluorescence value) to calculate the fold induction. The % induced values were calculated by determining how many events (out of 30,000 total) were present above the fluorescence threshold versus events that remained under the uninduced peak identified in the 0 hour timepoint. 30,000 events (cells) were analyzed at each indicated timepoint.

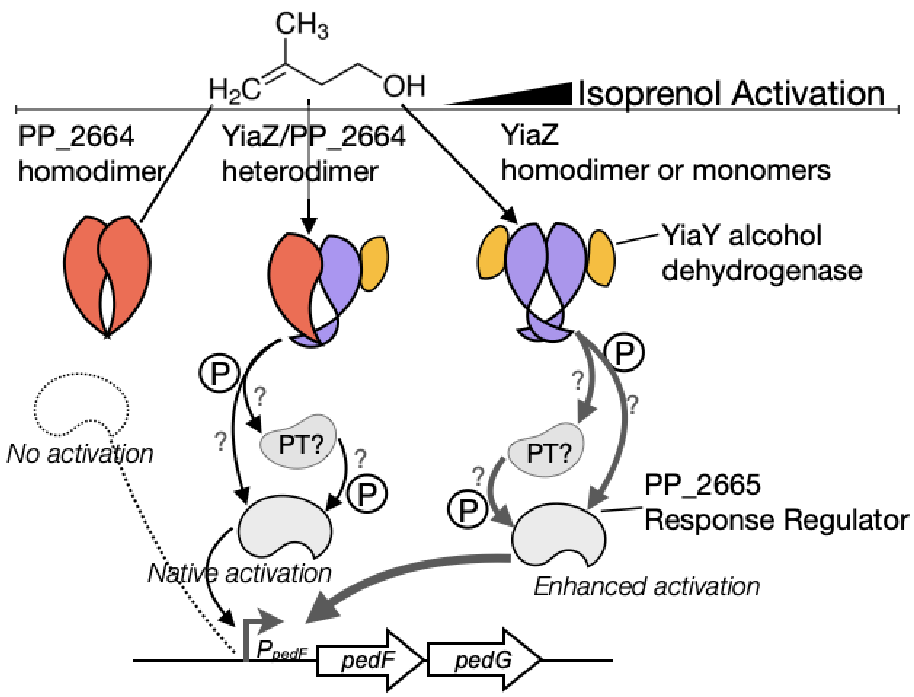


**Supplementary Figure 8. Model of the Isoprenol-Induced Signaling Cascade at the pedF promoter.** In this model, YiaY enzymatically reduces isoprenol to an aldehyde (e.g., 3-methyl-3-butenal). YiaY may also physically stabilize the modified ligand for recognition by YiaZ. Activated YiaZ (through autophosphorylation or trans-phosphorylation) initiates the phosphorelay using an unknown Hpt, leading to phosphorylation of PP_2665. This in turn enables binding to downstream DNA sequences to activate transcription in concert with a sigma factor. The extent of isoprenol pathway activation varies within a cell population due to stochastic differences arising from expression levels and the ratio of YiaZ homodimers, YiaZ/PP_2664 heterodimers, and PP_2664 homodimers. PP_2664 homodimers are not activated by isoprenol. Cells expressing only YiaZ homodimers show an improved response to isoprenol. Different ligands (short chain alcohols, diols) will have different preferences for homodimer vs heterodimer-mediated signaling cascade activation (refer to **Figure 3**). We do not know the kinetics or stability of the monomer to dimer formation and cannot exclude that monomers are active *in vivo*.

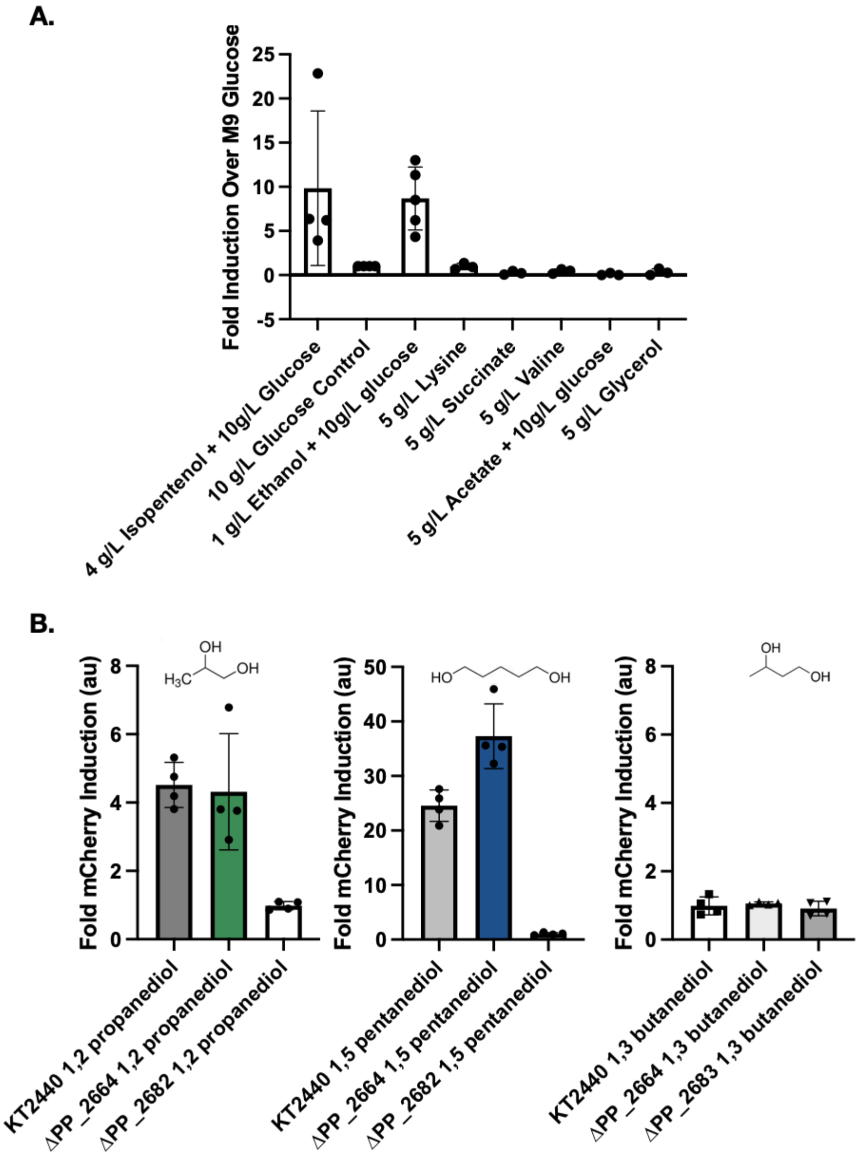


**Supplemental Figure 9. Additional Ligands for the *pedF-RBS-mCherry* Biosensor.** Refer to Figure 3. **(A)** Isoprenol biosensor activity assay with the analytes and concentrations indicated in WT P. putida with the plasmid born system. **(B)** Diol fold cherry activation of the biosensor are shown with a 11.6 mM concentration of each analyte in WT, ∆PP_2664, and ∆yiaY/∆PP_2682 (left and middle plots) and ∆yiaZ/∆PP_2683 (right plot). All samples were measured 28 hours after addition of analyte. All datapoints are shown and the error bar (if used) indicates standard deviation from the mean.

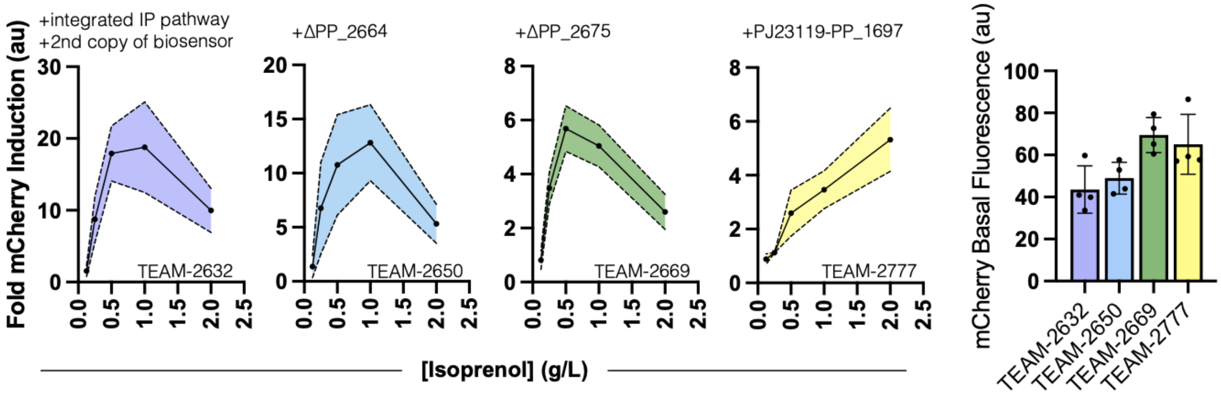


**Supplemental Figure 10. Characterization of Intermediate Strains Generated Towards a Genomically Integrated Isoprenol Biosensor.** Refer to Figure 4. Intermediate unified isoprenol production and biosensor strains were tested for biosensor activation with the indicated concentrations of isoprenol. Background fluorescence levels for each strain in the control condition are indicated on the far right panel. The linear dynamic range changes with each modification. Samples were measured 48 hours after induction. For the fold mCherry samples, the solid black line indicates the mean value from 4 biological replicates and the shaded colored area indicates the standard deviation. For the basal fluorescence readings, all datapoints are shown and the error bar indicates standard deviation from the mean.

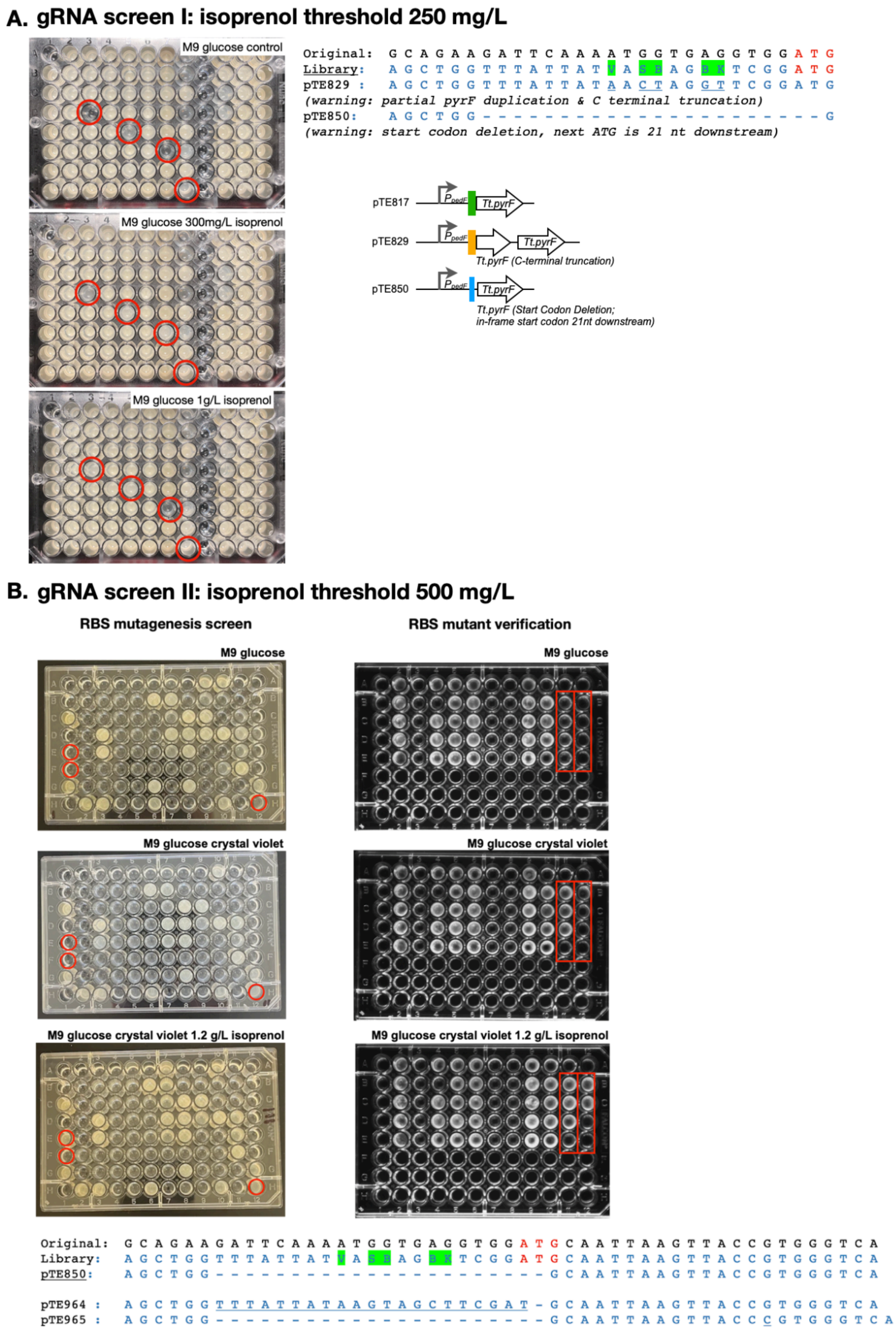


**Supplemental Figure 11. Representative High Throughput pedF-RBS-pyrF Screening Workflow for the gRNA growth screening.** Round 1 is described in **(A)** and round 2 is in **(B).** A representative screening plate for RBS mutants is shown with additional verification of the 2nd round verification plasmid screen on the right hand side of B. RBS mutant sequences are diagrammed comparing the original RBS sequence to the isolated plasmid.

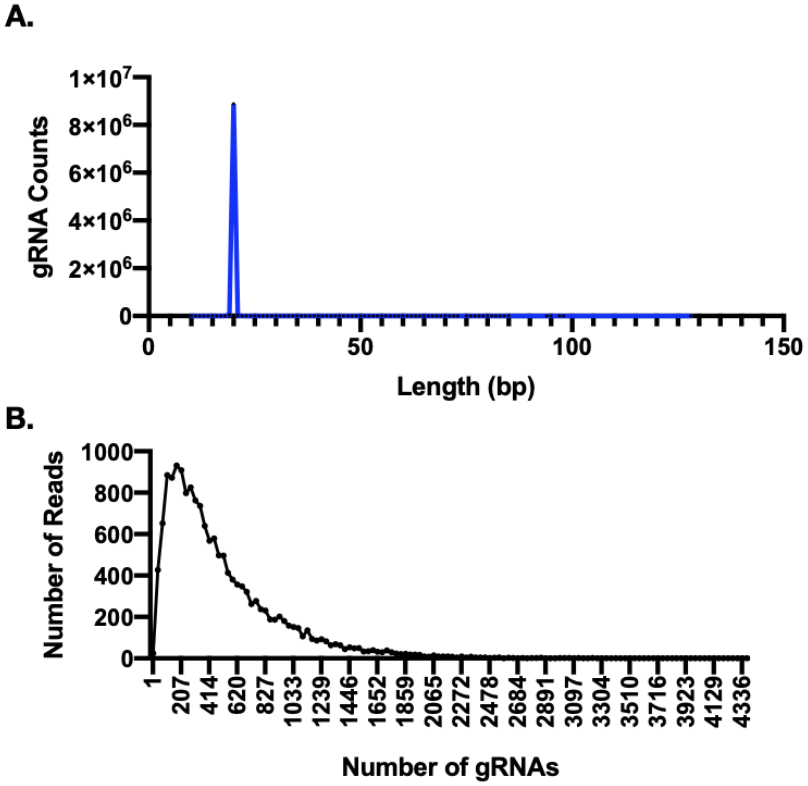


**Supplemental Figure 12. Characterization of Pooled gRNA Sequence Distribution After HiFi Assembly in *E. coli*.** Distribution of pooled gRNA sequences after HiFi assembly and transformation and extraction from *E. coli*. The data illustrates the representation and diversity of the gRNA library used for downstream CRISPRi screening in *P. putida*. (A.) The x-axis shows the count of gRNA sequences with the sequence size identified of the gRNA in the plasmid. (B) Histogram of gRNAs binned by number of reads, ranging from a low of 1 reads up to a maximum of 4,336 reads. This indicates the relative abundance or frequency of each gRNA in the library. The data was obtained by next-generation Illumina sequencing (NGS) of the plasmid library.

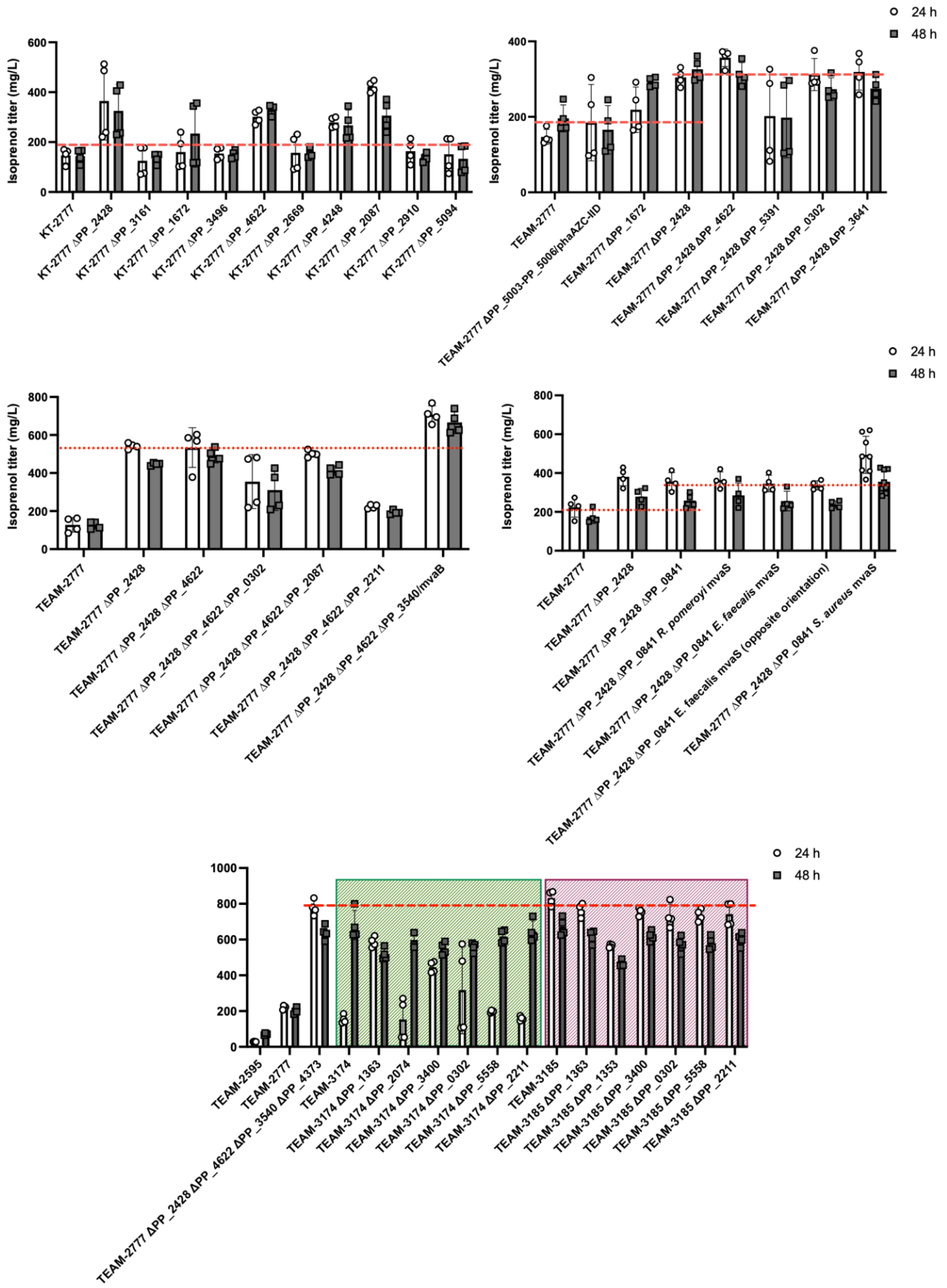


**Supplemental Figure 13. Representative isoprenol titers in deletion strains.** All strains shown on the same graph were assayed in parallel in the same M9 medium 2% glucose and GC analysis. The strain genotypes are indicated along the X axis and the analyzed timepoints for isoprenol quantification are indicated in the legend. All datapoints are shown and the error bars indicate standard deviation from the mean.

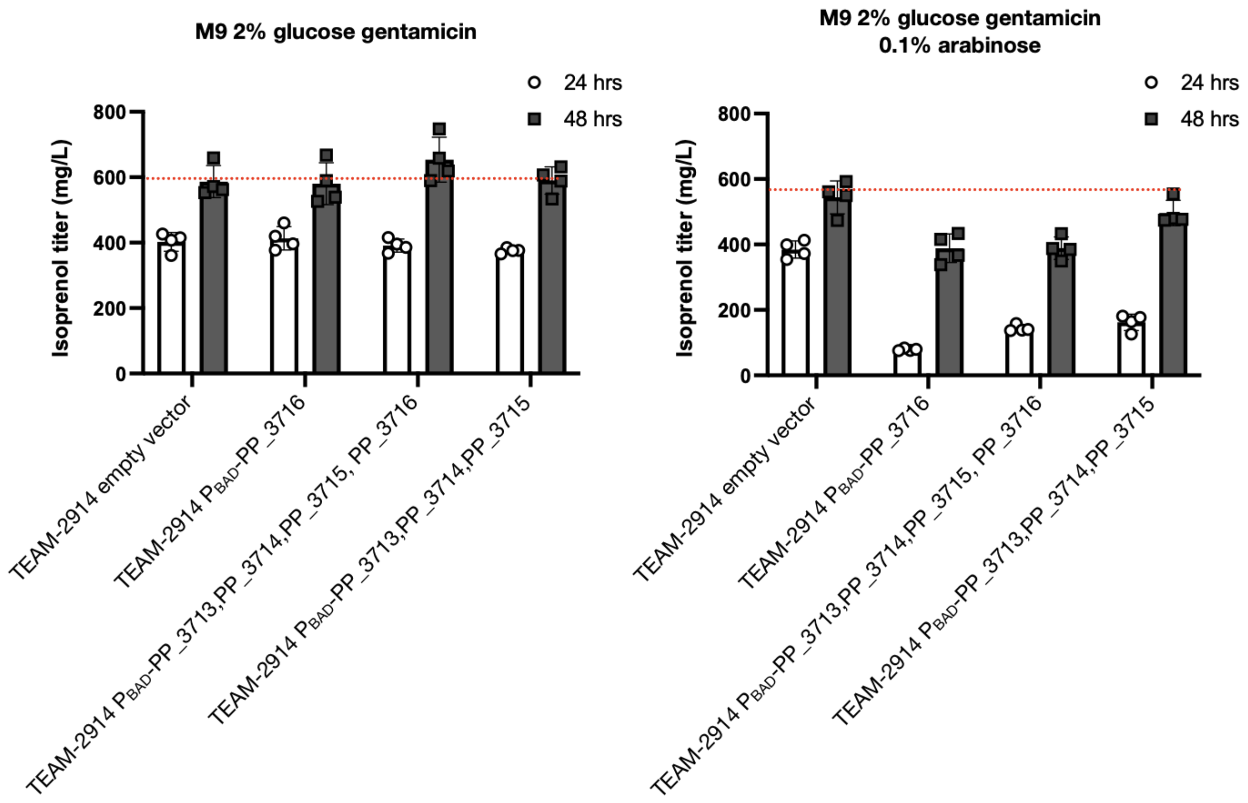


**Supplemental Figure 14. *catCBA-1* Overexpression Does Not Improve Titer.** Isoprenol production in strain TEAM-2914 harboring overexpressing constructs encoding the aromatic-related catabolism operon *catCBA-1* were compared to an empty vector control. Strains were cultivated in M9 2% glucose minimal media with 1μM CV added at the start of the time course to induce the isoprenol pathway and compared in parallel to samples additionally induced with arabinose (0.1% w/v). Samples were harvested and prepared for isoprenol quantification at the indicated timepoints. No improvements to isoprenol titer were observed with or without the inducer compared to the empty vector (pTE408) control. All datapoints are shown and the error bars indicate standard deviation from the mean.

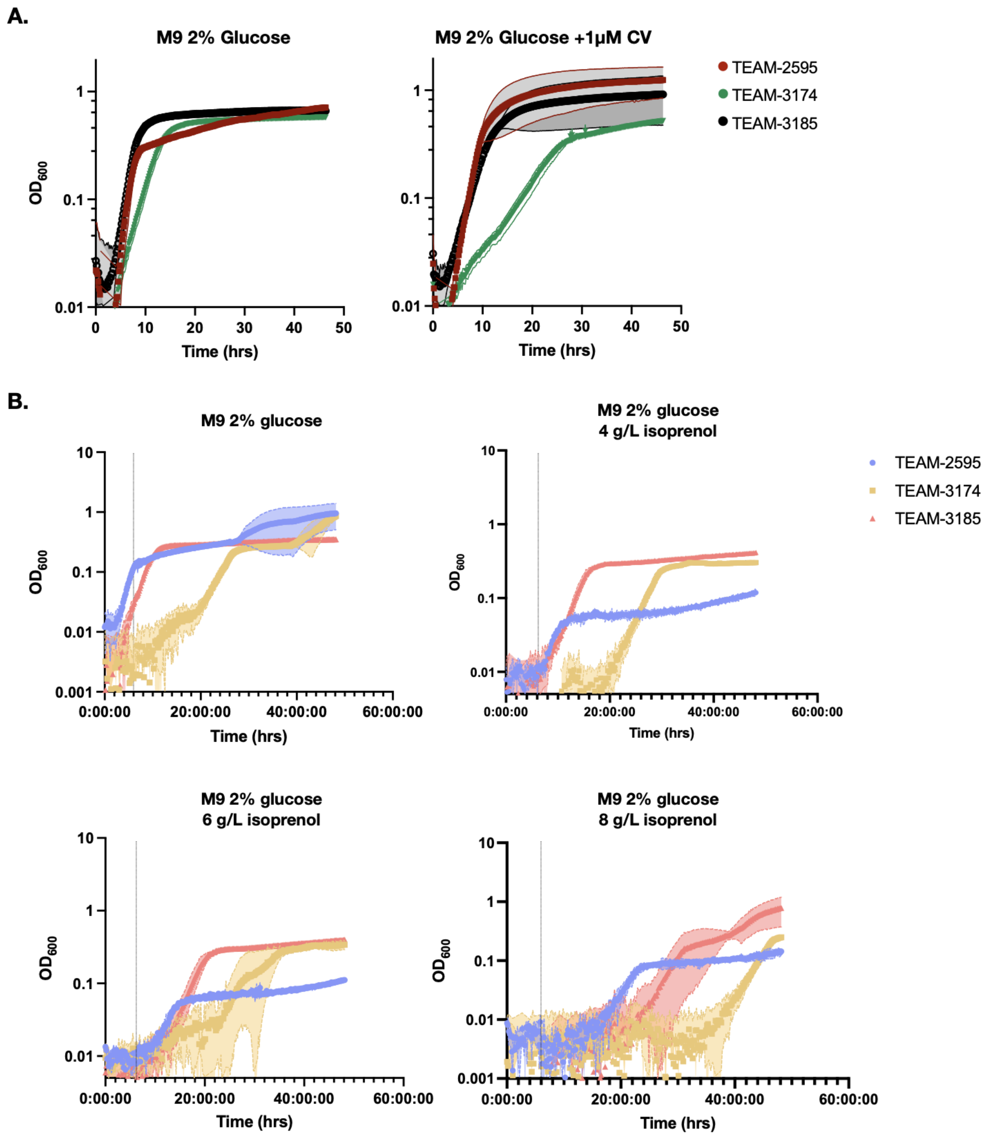


**Supplemental Figure 15. Kinetic growth analysis of high producer strains. (A.)** Growth curves in the absence and presence of crystal violet. **(B.)** Isoprenol tolerance growth curves in M9 minimal media supplemented with the indicated concentration of isoprenol. Samples in triplicate were read every 15 minutes with a microtiter dish plate reader at $OD_{600}$. The shaded colored area fill surrounding the solid color line indicates the standard deviation from the mean of measurements between 4 biological replicates. Strains are indicated in the sample legend to the right. A dotted black line indicates the 3 hour timepoint on the X axis to facilitate comparison of samples.

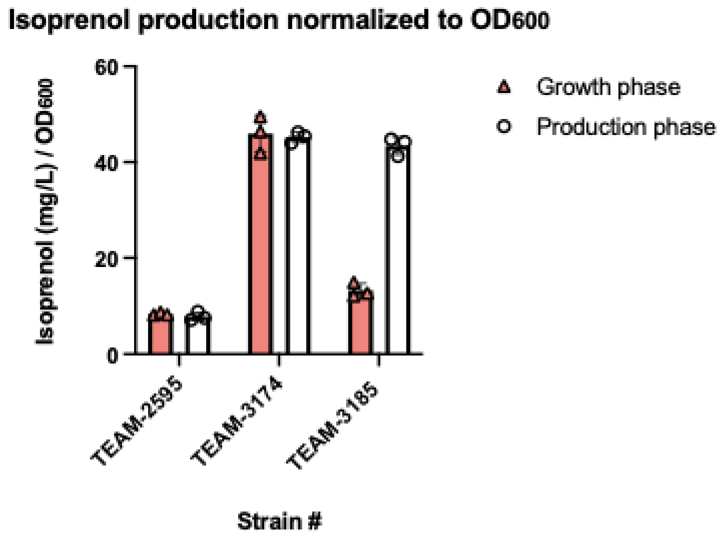


**Supplemental Figure 16. Specific isoprenol titer.** Refer to Figure 6A, 6C. Strains of the indicated genotypes were normalized to $OD_{600}$ for samples harvested in growth phase and the 24 hour production phase timepoint used for proteomics and metabolomics analysis. Analysis of glucose consumption indicated that all strains fully consumed glucose by the 24 hour timepoint. All datapoints are shown and the error bars indicate standard deviation from the mean.

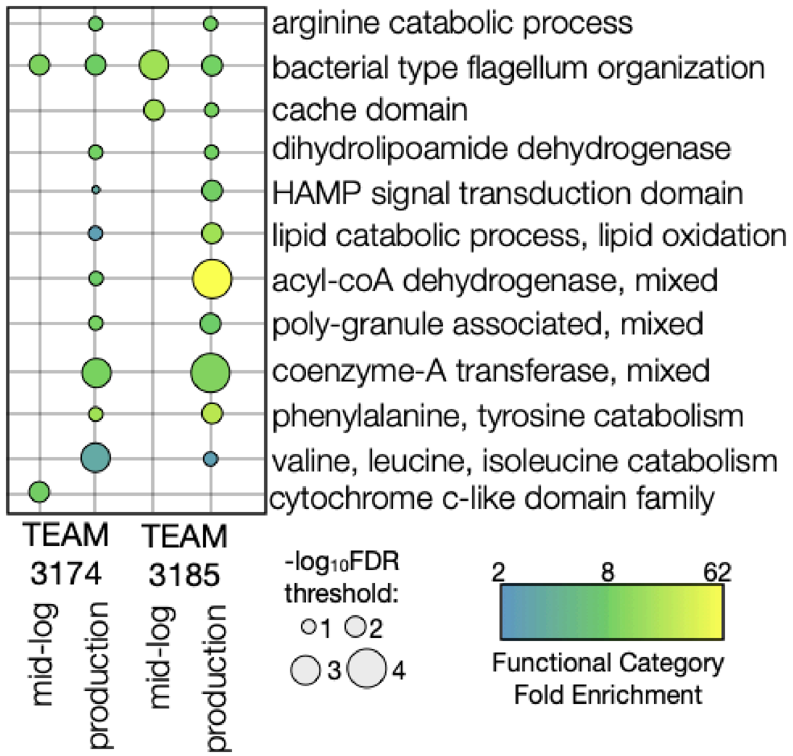


**Supplemental Figure 17. Analysis of Proteomics Enriched Pathways Using GO-term Enrichment With SHINY-GO.** Gene Ontology (GO) term enrichment analysis of differentially expressed proteins from the proteomics data, conducted using ShinyGO (Ge et al. 2020). The analysis identifies biological pathways and functional categories that are significantly overrepresented among the proteins showing altered expression levels from TEAM-3174 and TEAM-3185 each compared to TEAM-2595. The size and color of the circles correspond to the statistical significance of the enrichment, with larger circles indicating more statistical likelihood of significance with a lower false discovery rate while the color indicates the fold enrichment of the number of genes over the total number of genes in any gene category. The labels indicate the specific GO terms, providing an overview of the enriched pathways and functional categories. A selection of GO terms from non-duplicate categories are shown refer to **Supplementary Data 1-6** for the full SHINY-GO analysis.

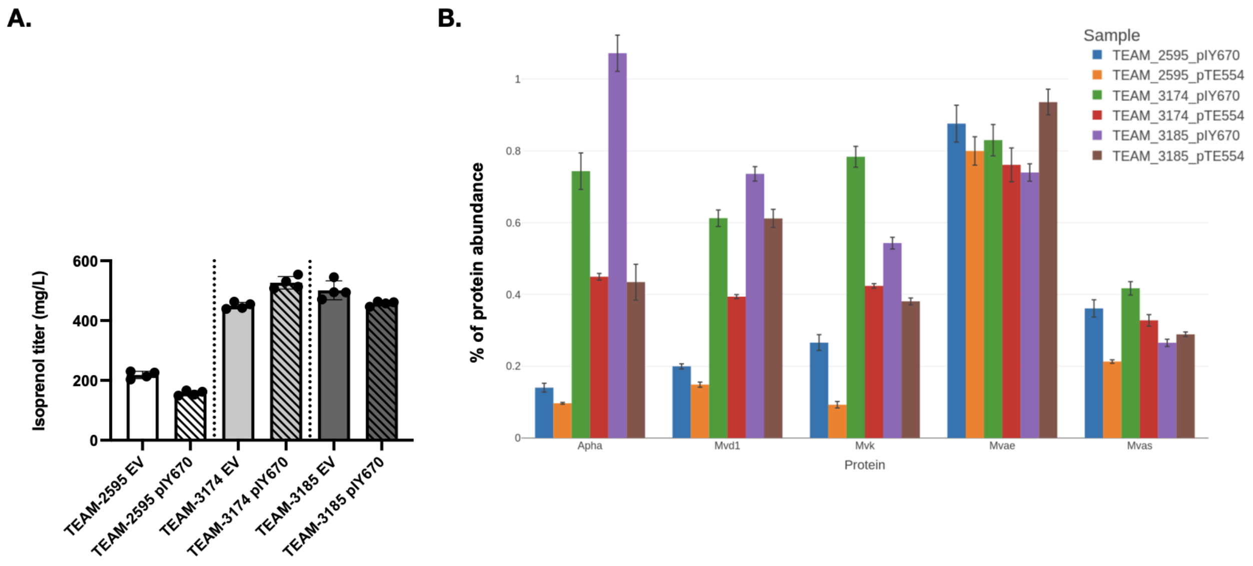


**Supplemental Figure 18. pIY670 plasmid pathway overexpression.** A. Isoprenol production was assayed in TEAM-2595, TEAM-3174 and TEAM-3185 harboring an additional copy of the isoprenol pathway with the plasmid-based construct pIY670 compared to an empty vector control (EV, “pTE554”). The production run was performed in M9 minimal media with 2% of glucose using 2% of arabinose as the inducer and the isoprenol titers displayed were obtained at 24 hrs. B. Comparison of the isoprenol pathway protein abundance in the strains harboring the pIY670 or the EV (pTE554) constructs by proteomics 24hr post-induction. In panel A, all datapoints are shown and the error bars indicate standard deviation from the mean. In panel B, the mean value is plotted and the error bar indicates standard deviation from the mean.

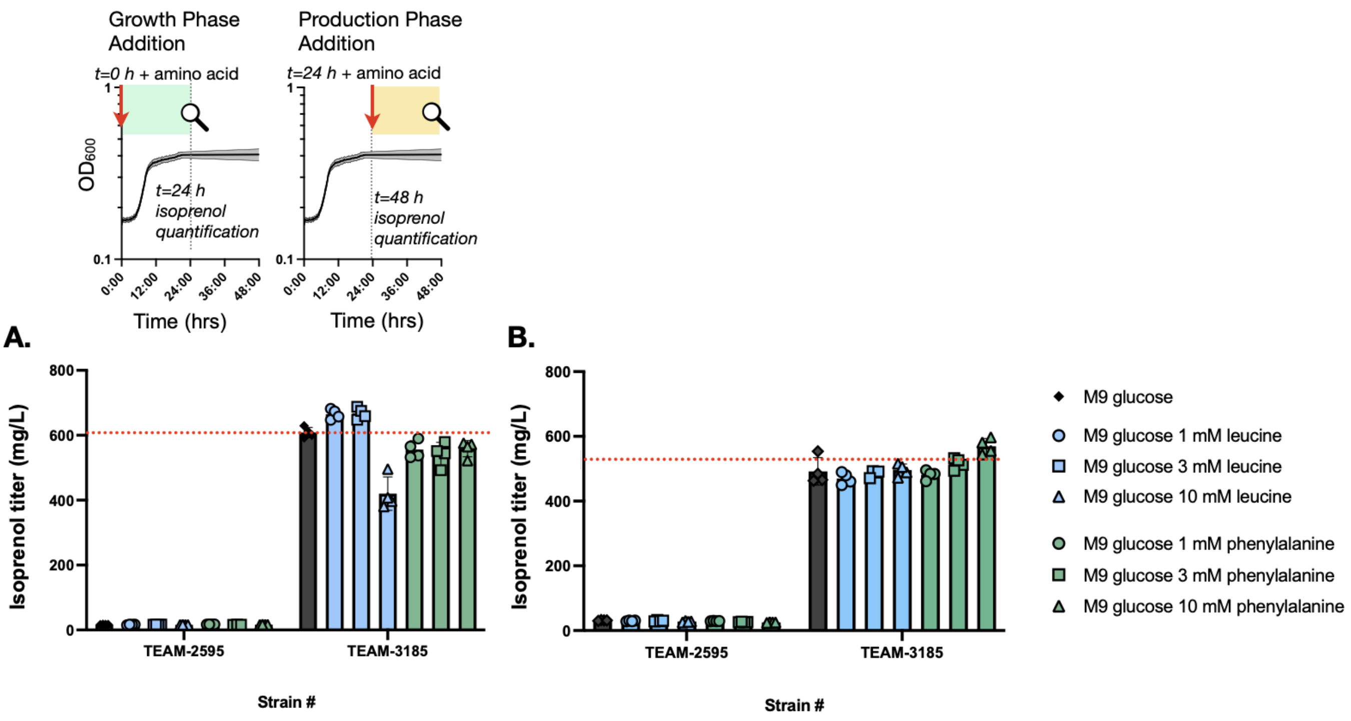


**Supplemental Figure 19. Amino acid supplementation during different growth phases modulates isoprenol titer.** Amino acid supplementation to isoprenol production cultures at the beginning of the production run (A, growth phase cultures) and 24 hr after the beginning of the production run (B, production phase cultures). See experimental workflow on top. In B, samples were then harvested at the 48 hour timepoint (24 hours after amino acid addition). Isoprenol producer strains TEAM-2595 and TEAM-3185 were tested with the concentrations of leucine and phenylalanine indicated in the figure legend to the right. All datapoints are shown and the error bars indicate standard deviation from the mean.

#### Supplemental Table 1. Major input parameters used to develop techno-economic analysis model in this study

| Parameter | Unit | Current state of technology scenario | | | | Optimal future case |
|---|---|---|---|---|---|---|
| | | Without amino acids | | With leucine alone | With leucine and phenylalanine | With leucine and phenylalanine |
| Biorefinery size (Baral et al. 2019) | bone-dry metric ton (bdt)/day | 2000 | | 2000 | 2000 | 2000 |
| Biomass sorghum feedstock cost (Baral et al. 2025, 2020) | $/bdt | 118.17 | | 118.17 | 118.17 | 87.48 |
| Moisture | % | 20 | | 20 | 20 | 20 |
| **Biomass composition (Baral et al. 2019; Yang et al. 2021; Magurudeniya et al. 2021)** | | | | | | |
| Acetate | wt% | 2.2 | | 2.2 | 2.2 | 0.9 |
| Ash | wt% | 4 | | 4 | 4 | 2.2 |
| Cellulose | wt% | 35.4 | | 35.4 | 35.4 | 40 |
| Extractives | wt% | 12.31 | | 12.31 | 12.31 | 12.06 |
| Hemicellulose | wt% | 20.7 | | 20.7 | 20.7 | 29.79 |
| Lignin | wt% | 21 | | 21 | 21 | 9.89 |
| Protein | wt% | 4.39 | | 4.39 | 4.39 | 5.16 |
| **Biomass deconstruction (Magurudeniya et al. 2021)** | | | | | | |
| Solid loading rate | wt% | 30 | | 30 | 30 | 40 |
| Ionic liquid (IL) loading rate | wt% | 5 | | 5 | 5 | 5 |
| IL cost | $/kg | 2 | | 2 | 2 | 1 |
| Pretreatment time | h | 3 | | 3 | 3 | 1 |
| Protein to amino acids | % | 70 | | 70 | 70 | 90 |
| Pretreatment temperature | % | 140 | | 140 | 140 | 140 |
| Acetate to acetic acid | wt% | 95 | | 95 | 95 | 100 |
| Sulfuric acid loading | kg/kg-IL | 0.15 | | 0.15 | 0.15 | 0.1 |
| Sulfuric acid cost | $/kg | 0.14 | | 0.14 | 0.14 | 0.14 |
| Enzyme loading rate | mg/g-glucan | 29.4 | | 29.4 | 29.4 | 10 |
| Solid loading for hydrolysis | wt% | 20 | | 20 | 20 | 25 |

| | | | | | | |
|---|---|---|---|---|---|---|
| Cellulose to glucose | wt% | 75.86 | | 75.86 | 75.86 | 95 |
| Xylan to xylose | wt% | 60.76 | | 60.76 | 60.76 | 90 |
| Hydrolysis time | h | 72 | | 72 | 72 | 48 |
| Enzyme price | $/kg-protein | 5 | | 5 | 5 | 5 |
| Hydrolysis temperature | °C | 50 | | 50 | 50 | 50 |
| **Bioconversion** | | | | | | |
| Inoculum cost (Baral et al. 2025) | $/kg | 0.004 | | 0.004 | 0.004 | 0.004 |
| Inoculum loading rate (Baral et al. 2025) | % | 10 | | 10 | 10 | 5 |
| Corn steep liquor (CSL) cost (Humbird et al. 2011; Davis et al. 2018) | $/kg | 0.07 | | 0.07 | 0.07 | 0.07 |
| Diammonium phosphate (DAP) cost (Humbird et al. 2011; Davis et al. 2018) | $/kg | 0.36 | | 0.36 | 0.36 | 0.36 |
| CSL loading (Humbird et al. 2011; Davis et al. 2018) | wt% | 0.25 | | 0.25 | 0.25 | 0.25 |
| DAP loading (Humbird et al. 2011; Davis et al. 2018) | g/L | 0.33 | | 0.33 | 0.33 | 0.33 |
| Ammonium sulphate loading[δ] | g/L | 9.3 | | 9.3 | 9.3 | 9.3 |
| Ammonium sulphate cost (Baral et al. 2025) | $/kg | 0.18 | | 0.18 | 0.18 | 0.16 |
| Air supply[ϕ] | vvm | 0.5 | | 0.5 | 0.5 | 0.2 |
| Bioconversion time[δ] | h | 24 | | 24 | 24 | 24 |
| Glucose-to-isoprenol yield[δ] | % by mass | 0.040 | | 0.048 | 0.056 | 0.31 |
| Xylose-to-isoprenol yield[θ] | % by mass | - | | - | - | 0.31 |

| | | | | | | |
|---|---|---|---|---|---|---|
| Leucine loading[δ] | g/L | - | | 1.31 | 1.31 | 1.31 |
| Leucine cost[□] | $/kg | - | | 3 | 3 | 3 |
| Phenylalanine loading[δ] | g/L | - | | - | 1.65 | 1.65 |
| Phenylalanine cost[ϒ] | $/kg | - | | - | 4 | 4 |
| Leucine-to-isoprenol yield[θ] | wt% | - | | - | - | 0.6 |
| Phenylalanine-to-isoprenol yield[θ] | wt% | - | | - | - | 0.67 |
| **Recovery and separation (Baral et al. 2019)** | | | | | | |
| Isoprenol recovery | % | 95 | | 95 | 95 | 98 |
| IL-recovery | % | 95 | | 95 | 95 | 99 |
| **Wastewater treatment (Humbird et al. 2011; Davis et al. 2018)** | | | | | | |
| Organic matter to biogas | wt% | 86 | | 86 | 86 | 86 |
| Nutrient cost | $/kg | 0.7 | | 0.7 | 0.7 | 0.45 |
| Nutrients loading for anaerobic digester | wt% | 0.05 | | 0.05 | 0.05 | 0.02 |
| **Onsite energy and utilities (Humbird et al. 2011; Davis et al. 2018)** | | | | | | |
| Boiler chemicals cost | $/kg | 5 | | 5 | 5 | 5 |
| Natural gas cost | $/kg | 0.22 | | 0.22 | 0.22 | 0.22 |
| Process water cost | $/kg | 0.00022 | | 0.00022 | 0.00022 | 0.00022 |

[δ]Based on the experimental data described in Figure 6 and Supplemental Figure 18 in this study.
[ϕ]Considered in this study to meet sufficient oxygen for cell redox balancing.
[θ]90% of the theoretical yield determined in this study.
[□]Globalsources. Available at: https://www.globalsources.com/Organic-chemical/Chemical-1201037043p.htm?source=Search_Product_to_PP Tianjin Saiyite Import and Export Co., Ltd, China. Accessed March 4, 2025. $0.9-$5 USD / kg leucine, depending on volume.
[ϒ]Globalsources. Available at: https://www.globalsources.com/Organic-chemical/Chemical-1201037043p.htm?source=Search_Product_to_PP Jbs Sa Decalcio Ltd, South Africa. Accessed March 4 2025. $3-$5 USD / kg, depending on volume.

**Supplemental Table 2. Lower pedF-RBS-pyrF Isoprenol Threshold: Enriched gRNAs.**

| Gene | Function | Functional Category |
|---|---|---|
| phaAZC-II (PP_5003-PP_5005) | Pha granule formulation genes | carbon storage |
| PP_5007 | Pha granule protein regulator | carbon storage |
| alkB (PP_3400) | Alkylated DNA repair protein | DNA repair |
| PP_2428 (SotB) | MFS transporter, DHA1 family, purine ribonucleoside efflux pump | efflux pump |
| PP_3168 | Benzoate-specific porin | efflux pump |
| PP_3641 | Cytosine/purine/uracil/thiamine/allantoin permease family protein | efflux pump |
| PP_4842 (urtB) | Branched-chain amino acid transport system permease protein | efflux pump |
| mvaB / PP_3540 | Hydroxymethylglutaryl-CoA lyase | enzyme |
| PP_0302 | L-carnitine 3-dehydrogenase or benzoate-related degradation | enzyme |
| PP_1409 | ribosomal small subunit pseudouridine synthase A; pyrimidine metabolism | enzyme |
| PP_1672 | Coenzyme B12 biosynthesis protein | enzyme |
| PP_2145 | beta-N-acetylglucosaminidase nucleotide sugar metabolism | enzyme |
| PP_2910 | L-2-hydroxyglutarate oxidase (EC 1.1.3.15) | enzyme |
| PP_3161 | Benzoate 1,2-dioxygenase subunit alpha benzoate degradation | enzyme |
| PP_5032 | Histidine ammonia-lyase | enzyme |
| PP_5094 | Enzyme, alanine racemace used for cell wall biosynthesis | enzyme |
| PP_0841 | Rrf2 family transcriptional regulator, iron-sulfur cluster assembly transcription factor | signal transduction |
| PP_2211 | Transcriptional regulator, araC family | signal transduction |
| PP_2739 | Histidine kinase | signal transduction |
| PP_3496 | Putative translation initiation inhibitor, yjgF family | signal transduction |
| PP_4197 | Amino acid related transcriptional regulator | signal transduction |
| PP_4373 fleQ | Flagella related regulator | signal transduction |
| PP_4622 | Transcriptional regulator, IclR family | signal transduction |

| Gene | Function | Functional Category |
|---|---|---|
| PP_5244 | Transcriptional regulator | signal transduction |
| PP_2087 (cmpx) | Unknown transmembrane protein | unknown |
| PP_2669 | Unknown outer membrane protein | unknown |
| PP_3883 | Phage holin protein | unknown |
| PP_5391 | Conserved hypothetical protein | unknown |

**Supplemental Table 3. Second Round pedF-RBS-pyrF with Higher Isoprenol Threshold: Enriched gRNAs.**

| Gene | Function | Functional Category |
|---|---|---|
| PP_1656 relA | ppGpp metabolism regulator | carbon storage |
| PP_4214 | Pyoverdin biosynthesis protein PvdN | carbon storage |
| ogt (PP_3017) | DNA methyltransferase | DNA repair |
| PP_2311 | TatD DNase family protein | DNA repair |
| hisQ (PP_4485) | Arginine/ornithine transport system permease protein | efflux pump |
| PP_0101 | Transmembrane Sulfate transporter family protein in cluster with carbonic anhydrase | efflux pump |
| PP_1363 | Phosphate transporter periplasmic component | efflux pump |
| PP_2667 | ABC transporter protein | efflux pump |
| PP_2974 | Transmembrane protein, unknown | efflux pump |
| PP_3221 | ABC transporter, possibly peptides | efflux pump |
| PP_3408 cbtA | Transmembrane protein cobalt transporter | efflux pump |
| PP_3480 | Transmembrane protein related to secretion | efflux pump |
| glgP (PP_5041) | Glycogen phosphorylase (related to degradative process) | enzyme |
| PP_0056 | Choline dehydrogenase, related to glycine serine threonine metabolism | enzyme |
| PP_1412 | Related to central metabolism acetyl-coA processing | enzyme |

| Gene | Function | Functional Category |
|---|---|---|
| PP_2869 | Oxidoreductase, FMN-binding | enzyme |
| PP_4322 | Cytochrome c-type biogenesis protein CcmF | enzyme |
| PP_4592 | Transmembrane metal dependent hydrolase | enzyme |
| PP_0055 | Transcriptional regulator | signal transduction |
| PP_0191 | Sigma-D family regulator, transcriptional regulatory protein AlgQ | signal transduction |
| PP_2074 | Transcriptional regulator LysR family | signal transduction |
| PP_2940 | YoeB-YefM toxin-antitoxin pair and DNA binding transcriptional repressor | signal transduction |
| PP_4462 | RNA processing related; putative 4-hydroxy-4-methyl-2-oxoglutarate aldolase | signal transduction |
| PP_4902 | Oligoribonuclease; RNA processing and degradation | signal transduction |
| PP_5558 | Transcriptional regulator | signal transduction |
| PP_0131 | Unknown transmembrane protein | unknown |
| PP_1353 | Membrane protein unknown function | unknown |
| PP_2483 | Conserved protein of unknown function | unknown |
| PP_2710 | Conserved protein of unknown function | unknown |
| PP_4706 | Unknown protein; monooxygenase domain | unknown |

**Supplemental Table 4. Strains Used in This Study.**

| Accession No. | TEAM ID# | Genotype | Reference |
|---|---|---|---|
| JBEI-13809 | TEAM-911 | *Pseudomonas putida* KT2440 | ATCC-47054, Nieto et al 1990 |
| | TEAM-1510 | *P. putida* KT2440 ΔPP_ 2675 | This study |
| JBEI-266166 | TEAM-2056 | Pp KT2440 ΔPP_ 2664 | This study |
| JBEI-266167 | TEAM-2052 | Pp KT2440 ΔPP_ 2665 | This study |
| JBEI-266168 | TEAM-2054 | Pp KT2440 ΔPP_ 2671 | This study |
| JBEI-266169 | TEAM-2012 | Pp KT2440 ΔPP_ 2683 (*yiaZ*) | This study |
| JBEI-266170 | TEAM-2176 | Pp KT2440 ΔPP_ 2682 (*yiaY*) | This study |
| JBEI-266171 | TEAM-2186 | Pp KT2440 pJ23119-yiaY-yiaZ | This study |
| JBEI-266172 | TEAM-2188 | Pp KT2440 $P_{BAD}$-yiaY-yiaZ | This study |
| JBEI-266173 | TEAM-2251 | Pp KT2440 ΔPP_ 2664 $P_{J23119}$-yiaY-yiaZ | This study |
| JBEI-266174 | TEAM-2431 | Pp KT2440 $P_{J23100}$-PP_2666,PP_2665 | This study |
| JBEI-266175 | TEAM-2555 | Pp KT2440 $P_{J23100}$-PP_2666,PP_2665 PP_5402intergenic::$P_{pedF}$-RBS-mCherry | This study |
| JBEI-266176 | TEAM-2583 | Pp KT2440 $P_{J23100}$-PP_2666,PP_2665 PP_5402intergenic::$P_{pedF}$-RBS-mCherry PP_5322intergenic::$P_{CV}$-mvaS,mvaE | This study |
| JBEI-251169 | TEAM-2595 | Pp KT2440 $P_{J23100}$-PP_2666,PP_2665 PP_5402intergenic::$P_{pedF}$-RBS-mCherry PP_5322intergenic::$P_{CV}$-mvaS,mvaE PP_0871intergenic::$P_{trc}$-MKmm,PMDHKQ,ahpA | This study |
| JBEI-266178 | TEAM-2632 | Pp KT2440 $P_{J23100}$-PP_2666,PP_2665 PP_5402intergenic::$P_{pedF}$-RBS-mCherry PP_5322intergenic::$P_{CV}$-mvaS,mvaE PP_0871intergenic::$P_{trc}$-MKmm,PMDHKQ,ahpA PP_3159intergenic::$P_{pedF}$-RBS-mCherry | This study |
| JBEI-266179 | TEAM-2650 | Pp KT2440 $P_{J23100}$-PP_2666,PP_2665 PP_5402intergenic::$P_{pedF}$-RBS-mCherry PP_5322intergenic::$P_{CV}$-mvaS,mvaE PP_0871intergenic::$P_{trc}$-MKmm,PMDHKQ,ahpA PP_3159intergenic::$P_{pedF}$-RBS-mCherry ΔPP_ 2664 | This study |
| JBEI-266180 | TEAM-2669 | Pp KT2440 $P_{J23100}$-PP_2666,PP_2665 PP_5402intergenic::$P_{pedF}$-RBS-mCherry PP_5322intergenic::$P_{CV}$-mvaS,mvaE | This study |

| | | | |
|---|---|---|---|
| | | PP_0871intergenic::$P_{trc}$-MKmm,PMDHKQ,ahpA<br>PP_3159intergenic::$P_{pedF}$-RBS-mCherry ΔPP_ 2664 ΔPP_2675 | |
| JBEI-266181 | TEAM-2777 | Pp KT2440 $P_{J23100}$-PP_2666,PP_2665 PP_5402intergenic::$P_{pedF}$-RBS-mCherry<br>PP_5322intergenic::$P_{CV}$-mvaS,mvaE<br>PP_0871intergenic::$P_{trc}$-MKmm,PMDHKQ,ahpA<br>PP_3159intergenic::$P_{pedF}$-RBS-mCherry ΔPP_ 2664 ΔPP_2675<br>$P_{J23119}$-PP_1697 | This study |
| JBEI-266182 | TEAM-862 | Pp KT2440 ΔPP_1815/*pyrF* | This study |
| | TEAM-2847 | Pp TEAM-2777 ΔPP_ 2087 | This study |
| JBEI-266183 | TEAM-2822 | Pp TEAM-2777 ΔPP_ 2428 | This study |
| | TEAM-2906 | Pp TEAM-2777 ΔPP_ 2211 | This study |
| | TEAM-2910 | Pp TEAM-2777 ΔPP_ 1409 | This study |
| | TEAM-2853 | Pp TEAM-2777 ΔPP_ 5003-5006 | This study |
| | TEAM-2861 | Pp TEAM-2777 ΔPP_ 5007 | This study |
| JBEI-266184 | TEAM-2885 | Pp TEAM-2777 ΔPP_ 2428 ΔPP_ 4622 | This study |
| | TEAM-2896 | Pp TEAM-2777 ΔPP_ 2428 ΔPP_ 0841 | This study |
| | TEAM-2877 | Pp TEAM-2777 ΔPP_ 2428 ΔPP_ 2087 | This study |
| | TEAM-2918 | Pp TEAM-2777 ΔPP_ 2428 ΔPP_ 4622 ΔPP_ 2087 | This study |
| | TEAM-2919 | Pp TEAM-2777 ΔPP_ 2428 ΔPP_ 4622 ΔPP_ 2211 | This study |
| JBEI-266185 | TEAM-2914 | Pp TEAM-2777 ΔPP_ 2428 ΔPP_ 4622 ΔPP_ 3540 | This study |
| | TEAM-2930 | Pp TEAM-2777 ΔPP_ 2428 ΔPP_ 0841 ΔPP_2211 | This study |
| | TEAM-2946 | Pp TEAM-2777 ΔPP_ 2428 ΔPP_ 0841 + *E. faecalis* mvaS | This study |
| | TEAM-2949 | Pp TEAM-2777 ΔPP_ 2428 ΔPP_ 0841 + *S. aureus* mvaS | This study |
| | TEAM-3084 | Pp TEAM-2777 ΔPP_ 2428 ΔPP_ 4622 ΔPP_ 3540 ΔPP_ 4214 | This study |
| | TEAM-3089 | Pp TEAM-2777 ΔPP_ 2428 ΔPP_ 4622 ΔPP_ 3540 ΔPP_1656 | This study |
| | TEAM-3086 | Pp TEAM-2777 ΔPP_ 2428 ΔPP_ 4622 ΔPP_ 3540 ΔPP_2710 | This study |
| | TEAM-3053 | Pp TEAM-2777 ΔPP_ 2428 ΔPP_ 4622 ΔPP_ 3540 ΔPP_2974 | This study |
| | TEAM-3080 | Pp TEAM-2777 ΔPP_ 2428 ΔPP_ 4622 ΔPP_ 3540 ΔPP_3400 | This study |
| JBEI-266186 | TEAM-3032 | Pp TEAM-2777 ΔPP_ 2428 ΔPP_ 4622 ΔPP_ 3540 ΔPP_4373 | This study |

| | | | |
|---|---|---|---|
| JBEI-266187 | TEAM-3134 | Pp TEAM-2777 ΔPP_ 2428 ΔPP_ 4622 ΔPP_ 3540 ΔPP_4373 ΔPP_2710 | This study |
| JBEI-264843 | TEAM-3185 | Pp TEAM-2777 ΔPP_ 2428 ΔPP_ 4622 ΔPP_ 3540 ΔPP_4373 ΔPP_2074 | This study |
| | TEAM-3115 | Pp TEAM-2777 ΔPP_ 2428 ΔPP_ 4622 ΔPP_ 3540 ΔPP_4373 + PP_3159intergenic::$P_{pedF}$-*S.pomeroyi*.mvaS | This study |
| | TEAM-3117 | Pp TEAM-2777 ΔPP_ 2428 ΔPP_ 4622 ΔPP_ 3540 ΔPP_4373 + PP_3159intergenic::$P_{pedF}$-*S. aureus*.mvaS | This study |
| | TEAM-3119 | Pp TEAM-2777 ΔPP_ 2428 ΔPP_ 4622 ΔPP_ 3540 ΔPP_4373 + PP_3159intergenic::$P_{pedF}$-*E. faecalis* mvaS | This study |
| JBEI-264842 | TEAM-3174 | Pp TEAM-2777 ΔPP_ 2428 ΔPP_ 4622 ΔPP_ 3540 ΔPP_4373 ΔPP_2710 + PP_1117intergenic::$P_{CV}$-*E. faecalis*.mvaS PP_5464intergenic::$P_{CV}$-*E. faecalis*.mvaS | This study |
| | TEAM-3168 | Pp TEAM-2777 ΔPP_ 2428 ΔPP_ 4622 ΔPP_ 3540 ΔPP_4373 ΔPP_2710 + PP_1117intergenic::$P_{CV}$-*S. pomeroyi* mvaS PP_5464intergenic::$P_{CV}$-*S. pomeroyi* mvaS | This study |

**Supplemental Table 5. Plasmids Used in This Study.** For all aCpf1 plasmids derived from pTE433β, see Supplemental Table 6. For the pooled dCpf1 library derived from pTE219, its storage in 1mL single use *E. coli* cryostocks is not compatible with our standard archival format and does not have an archival number.

| JBEI ID | TEAM # | Strain | Plasmid # | Description | Reference |
|---|---|---|---|---|---|
| JBEI-001791 | | *E. coli* | pRK2013 | tra+ mob+ (RK2) Km::Tn7 ColEl origin, helper plasmid, kanR *use for triparental conjugation* | (Figurski and Helinski 1979) |
| | TEAM-865 | *E. coli* | pTE219 | $P_{LacM}$-Fn.Cpf1-D917A PJ23119-PmeI-gRNA BBR1 oriT KanR (dCpf1) | (Banerjee et al. 2020) |
| | TEAM-1507 | *E. coli* | pTE365 | $P_{pobR}$-unoptimized-RBS-mCherry, PP_3538 $P_{clpB}$-mCitrine kanR BBR1 | this study |
| | TEAM-2448 | *E. coli* | pTE705 | $P_{pobR}$-optimizedRBS-mCherry, PP_3538 $P_{clpB}$-mCitrine kanR BBR1 | this study |
| JBEI-144837 | TEAM-1196 | *E. coli* | pTE317β | $P_{pedF}$-sfGFP kanR BBR1 *(no RBS optimization)* | this study |
| JBEI-266190 | TEAM-1959 | *E. coli* | pTE518 | $P_{pedF}$-RBS-mCherry gntR BBR1 | this study |
| | TEAM-2068 | *E. coli* | pTE542 | $P_{BAD}$-PP_2683/yiaZ $P_{pedF}$-RBS-mCherry gntR BBR1 | this study |

| JBEI ID | TEAM # | Strain | Plasmid # | Description | Reference |
|---|---|---|---|---|---|
| | TEAM-2092 | *E. coli* | pTE543 | $P_{J23119}$-PP_2683 $P_{pedF}$-RBS-mCherry gntR BBR1 | this study |
| | TEAM-2087 | *E. coli* | pTE544 | 1.5kb region genomic region containing PP_2681, yiaY, and yiaZ/PP_2683 $P_{pedF}$-RBS-mCherry gntR BBR1 | this study |
| | TEAM-2114 | *E. coli* | pTE556 | PP_2682,PP_2683 $P_{pedF}$-RBS-mCherry *gntR sacB (integration allelic exchange vector)* | this study |
| | TEAM-2130 | *E. coli* | pTE560 | $P_{BAD}$-PP_2682/yiaY,PP_2683/yiaZ $P_{pedF}$-RBS-mCherry gntR BBR1 | this study |
| | TEAM-2131 | *E. coli* | pTE561 | $P_{BAD}$-PP_2683+42aa $P_{pedF}$-RBS-mCherry gntR BBR1 *(assumes PP_2683 translation starts at a start codon 42 codons earlier in the PP_2682 reading frame)* | this study |
| | TEAM-2132 | *E. coli* | pTE562 | $P_{BAD}$-PP_2682/yiaY pedF-RBS-mCherry gntR BBR1 | this study |
| | TEAM-2149 | *E. coli* | pTE575 | 1.5kb region genomic region containing PP_2681, yiaY/PP_2682-G3C, and yiaZ/PP_2683 $P_{pedF}$-RBS-mCherry gntR BBR1 (introduces stop codon at start codon creating PP_2682-M1*) | this study |
| | TEAM-2172 | *P. putida* | pTE578 | 1.5kb region genomic region containing PP_2681, yiaY/PP_2682-Δ14, and yiaZ/PP_2683 pedF-RBS-mCherry gntR BBR1 (truncates the first 14 amino acids from YiaY/PP_2682) | this study |
| | TEAM-2206 | *E. coli* | pTE588 | T7p-6HIS-40-164aaPP_2683 kanR ColE1 | this study |
| | TEAM-3351 | *E. coli* | pTE710 | T7p-6HIS-TEV-PP_2682,PP_2683 kanR ColE1 | this study |
| | TEAM-2782 | *E. coli* | pTE906 | $P_{BAD}$-6HIS-PP_2664, 3FLAG-PP_2683 BBR1 gntR | |
| | TEAM-2800 | *E. coli* | pTE908 | $P_{BAD}$-3FLAG-TEV-PP_2683 BBR1 gntR | |
| | TEAM-2967 | *E. coli* | pTE959 | $P_{BAD}$-PP_2682 (yiaY)-H281A kanR | this study |
| | TEAM-1994 | *E. coli* | pTE520 | PP_5402intergenic::$P_{pedF}$-RBS-mCherry kanR sacB | this study |
| | TEAM-2477 | *E. coli* | pTE709 | PP_3159intergenic::$P_{pedF}$-RBS-mCherry kanR (*integrase plasmid derived from pSPIN-family backbone)* | this study |
| | TEAM-2662 | *P. putida* | pTE817 | $P_{pedF}$-Tt.pyrF RSF1010 gntR | this study |
| JBEI-266192 | TEAM-2732 | *P. putida* | pTE829 | $P_{pedF}$-RBSopt-D3-Tt.pyrF RSF1010 gntR "RBS-mut1" | this study |
| JBEI-266193 | TEAM-2740 | *P. putida* | pTE850 | $P_{pedF}$-RBSopt-P4-H5-Tt_pyrF RSF1010 gntR "RBS-mut2" | this study |
| JBEI-266194 | TEAM-3015 | *P. putida* | pTE964 | $P_{pedF}$-RBSmutH12-P4-H5-Tt_pyrF RSF1010 gntR | this study |
| JBEI-266195 | TEAM-3014 | *P. putida* | pTE965 | $P_{pedF}$-RBSmut1F-P4-H5-Tt_pyrF RSF1010 gntR | this study |
| JBEI-264945 | TEAM-2058 | *E. coli* | pIY670 | araC-pBAD-mvaS,mvaE ptrc-MKmm,PMDHKQ,aphA RK2 kanR | (Banerjee et al. |

| JBEI ID | TEAM # | Strain | Plasmid # | Description | Reference |
|---|---|---|---|---|---|
| | | | | | 2024) |
| | TEAM-2535 | *E. coli* | pTE744 | PP_5322intergenic::$P_{CV}$-mvaS,mvaE kanR sacB *(integration allelic exchange vector)* | this study |
| | TEAM-2536 | *E. coli* | pTE745 | PP_0871intergenic::$P_{trc}$-MKmm,PMDHKQ,aphA *gntR sacB (integration allelic exchange vector)* | this study |
| | TEAM-2783 | *E. coli* | pTE408 | $P_{BAD}$ BBR1 gntR *(empty vector)* | this study |
| | TEAM-2693 | *E. coli* | pTE847 | $P_{BAD}$-PP_1697 BBR1 gntR | this study |
| | TEAM-2692 | *E. coli* | pTE846 | $P_{BAD}$-PP_2211 BBR1 gntR | this study |
| | TEAM-2694 | *E. coli* | pTE848 | $P_{BAD}$-PP_4197 BBR1 gntR | this study |
| JBEI-236684 | TEAM-1673 | *E. coli* | pTE433β | $P_{LacM}$-RBSopt-Fn.Cpf1 PJ23119-PmeI-gRNA oriT BBR1 LacI- KanR | (Czajka et al. 2022) |
| | TEAM-2867 | *E. coli* | pTE924 | $P_{J23119}$-aCpf1-PP_2087 BBR1 kanR | this study |
| | TEAM-2814 | *E. coli* | pTE922 | $P_{J23119}$-aCpf1-PP_2428 BBR1 kanR | this study |
| | TEAM-2692 | *E. coli* | pTE846 | $P_{J23119}$-aCpf1-PP_2211 BBR1 kanR | this study |
| | TEAM-2803 | *E. coli* | pTE912 | $P_{J23119}$-aCpf1-PP_1409 BBR1 kanR | this study |
| JBEI-204825 | TEAM-1900 | *E. coli* | pTE504 | $P_{J23119}$-aCpf1-PP_5003 BBR1 kanR | (Czajka et al. 2022) |
| | TEAM-2809 | *E. coli* | pTE918 | $P_{J23119}$-aCpf1-PP_5007 BBR1 kanR | this study |
| | TEAM-2826 | *E. coli* | pTE930 | $P_{J23119}$-aCpf1-PP_4622 BBR1kanR | this study |
| | TEAM-3348 | *E. coli* | pTE934 | $P_{J23119}$-aCpf1-PP_0841 BBR1 kanR | this study |
| | TEAM-2571 | *E. coli* | pTE769 | $P_{J23119}$-aCpf1-PP_3540 BBR1 kanR | this study |
| | TEAM-3043 | *E. coli* | pTE996 | $P_{J23119}$-aCpf1-PP_4214 BBR1 kanR | this study |
| | TEAM-3023 | *E. coli* | pTE970 | $P_{J23119}$-aCpf1-PP_1656 BBR1 kanR | this study |
| | TEAM-3328 | *E. coli* | pTE993 | $P_{J23119}$-aCpf1-PP_2710 BBR1 kanR | this study |
| | TEAM-2314 | *E. coli* | pTE609 | $P_{J23119}$-aCpf1-PP_4373 BBR1 kanR | this study |
| | TEAM-3350 | *E. coli* | pTE978 | $P_{J23119}$-aCpf1-PP_2974 BBR1 kanR | this study |
| | TEAM-3039 | *E. coli* | pTE990 | $P_{J23119}$-aCpf1-PP_3400 BBR1 kanR | this study |
| | TEAM-3349 | *E. coli* | pTE991 | $P_{J23119}$-aCpf1-PP_2074 BBR1 kanR | this study |
| | TEAM-2923 | *E. coli* | pTE954 | $P_{pedF}$-RBS-*E. faecalis* mvaE kanR (*integrase plasmid derived from pSPIN-family* | this study |

| JBEI ID | TEAM # | Strain | Plasmid # | Description | Reference |
|---|---|---|---|---|---|
| | | | | *backbone)* | |
| | TEAM-2922 | *E. coli* | pTE952 | $P_{pedF}$-RBS-*S. aureus* mvaE kanR (*integrase plasmid derived from pSPIN-family backbone)* | this study |
| | TEAM-2921 | *E. coli* | pTE951 | $P_{pedF}$-RBS-*S. pomeroyi* mvaE kanR (*integrase plasmid derived from pSPIN-family backbone)* | this study |
| | TEAM-3109 | *E. coli* | pTE1007 | $P_{CV}$-RBS-*E. faecalis* mvaE kanR (*integrase plasmid derived from pSPIN-family backbone)* | this study |
| | TEAM-3107 | *E. coli* | pTE1005 | $P_{CV}$-RBS-*S. pomeroyi* mvaE kanR (*integrase plasmid derived from pSPIN-family backbone)* | this study |
| | TEAM-3108 | *E. coli* | pTE1006 | $P_{CV}$-RBS-*S. aureus* mvaE kanR (*integrase plasmid derived from pSPIN-family backbone)* | this study |
| | TEAM-2105 | *E. coli* | pTE554 (EV) | RK2 kanR *(empty vector)* | this study |

**Supplemental Table 6. Cpf1-gRNA Sequences Used in Recombineering to Make Deletion Mutants.** A 5’-TTTN-3’ PAM sequence was always used to identify compatible gRNA targeting sequences for assembly into plasmid pTE433β. Recombineering oligos were ordered without any basepair modifications or additional purifications.

| Gene target | Plasmid No. | gRNA Targeting Sequence 5'-3' | Recombineering Oligo 5'-3' |
|---|---|---|---|
| PP_0055 | pTE1004 | CTCGAGTTGCAGCGCAGCGGC | GCATGCCCGTTAAGAGGCCGGAACGAACTGCCATAAAGGACAAGCCTAGCCGAGtagAGCC<br>TTGCCGGCCAACAACAGGGCAGCGGCCCA |
| PP_0056 | pTE977 | GGTTATCCAAGGGGCAAGGTG | ACTCACTAAGATGGATCGGGACAAGAATAAAAAACAGGCGCGAGGCCCGGGTGCGTGCAGT<br>ACCGGCGGCTTGCGGTCAGAAGTCGCCGA |
| PP_0101 | pTE1012 | CCGGCGGCATGCCAGAAGCGA | GGCCGTGCGACTAGAGAAAGGAGAAACACGGTGAACATTACACAGGAGCGGGTTTACCCGC<br>GAATACGGTAGCGGCGGCAACGGTGATCG |
| PP_0131 | pTE1010 | CCGTCCCTGGATGCTGGCCAC | GAGGGTGATTATTGATTCCGTTGCTATACCTTGGTTATCAACTGGGCGCGTCGAAGCGATC<br>AGACCGGCAACGCCATGTAGAACTGGGTC |
| PP_0191 | pTE1001 | CGCCCTTCTGGTCGACTACGT | TGGTGCGAGGTGCAGCGGCATCTACCCAGAAGGGAAGAGATCGCC<br>GGTTGGCGTCAGTCGCCAACCGCCAGCAGTTCGATCTCGAATACC |
| PP_0841 | pTE933 | GCCGATATTTCCGAGCGCCAG | CGCATACTCGCACACATCCTGAAACTCCCGTGGTACCCAATAGCCCCCCCTGGGACGACGA<br>GCGACGCCACCGCCTGATAGGAGAGACAA |
| PP_1353 | pTE1018 | CCAGTTCCAGCAACACATTGC | ACGCGCGGCCTGTTCAACCGTGTACTTGCCGCTCGGCATCAGGAG<br>TGCCGCGTTCAAGCCGGGGCGCAAAGCGGCCCCAGCTTTTTCCGA |
| PP_1363 | pTE1003 | CAGGCAAACGCCGATACACTG | GGCATTTGAACGGGGCAGGGTATCCTTGCAAAAGGGTGTGAAGATCACAAAGGTGTGGTTT<br>ATTTCACGAGATTTGCTGCGCGTGCTGCG |
| PP_1409 | pTE912 | tattttgccttcgaagacctC | CCCCCTGGGACGACGAGCGACGCCACCGCCTGATAGGAGAGACAAGCCGTGACAGACAGCC<br>CGTGTCGTCATACGACCGCTCAGCGACAA |
| PP_1412 | pTE990 | CCCTGCACACCTACCCCAAAC | GGCCAGGCAAGCCATACTTGAGCGATGACTTGCGAGAGGTAGTCC<br>CCTGCTGGTCCGGGGGCTGCTTTTCAGCCCCCGCCTCCGCTCAGC |
| PP_1656/relA | pTE970 | CTCTTGAGGTAGAGAAAAAGG | GCCAGCACACTCGTATGGTGTGCTGCATGGAAAGGTAAAACAAAG<br>GACCGGGGCGTCTGCTTCGCGGGTAAACCCGCTCCTACAGGATGG |
| PP_1672 | pTE919 | TGTGCGCGGATACCAGCCTG | CCTGTCGCCGCGCATTGCCGTGGTGCCAGCCAGCGGAGAACCCTGGtgaGTCGATCGCGCC<br>ACTGCCCCGCCGTGTTGATCGCGGCGCCG |

| Gene target | Plasmid No. | gRNA Targeting Sequence 5'-3' | Recombineering Oligo 5'-3' |
|---|---|---|---|
| PP_1697 | pTE901 | TATAACAAGACGAGAGCGAGA | GGGCGACAGGCGGAGGATTGACAGCCTGTCAGGCTCAGGGAATAC TTGACAGCTAGCTCAGTCCTAGGTATAATGCTAGCatgCACCCATTGACCGGTGATGCTCG ACTGCCCCGCTACCAACAA |
| PP_1815/pyrF | pTE427 | CCTACCCGTGAGGCCGCCCT | GGTTCGGCACAGGCCCTTGCTGGACAGCCGCAGGTTAACAGGGCAGGGTCTCTTGGCAAGT CGAAAACGGCGCGCATTGTAAACGAAGT |
| PP_2074 | pTE991 | GACCCGCTGACCAGCAGGATG | ATGCTGGAATTGAACCCGAACAACAAGCACGGCCCGGGAGCACCC TGGATTTGCATCGTGCTGCAACTGGCTGCCGATGAAGTGCGCCGG |
| PP_2087/cmpx | pTE924 | GACCAAGGTAGCGAACTTCA | CCGCAACGCTGCCAATGGCAGCAAAGCAGATAAGGCCGATTGAATCCGCACAAAAGCTGTT AATGTATGCCGCCGCGAAATTCGACCCAC |
| PP_2145 | pTE927 | CGCGAACCGATTATCGCCAG | CTGCCCCTCGGTGGGCCTCTCTCTGTCTTCAATGAAGGATTACCCCTGCGCTGGCCTGTTC GTTCCCACAAGTCGGGTTTACCTGCGAAT |
| PP_2311 | pTE980 | CACGACCAACAGGCCGCAATC | CGCGGTTTTTGCTATGCTGGCCGCCCCGCGGCACGAGATAGCACCGTCTCGCGTGTTGTTG CGGGGCGGGTGAACACAGTTGATATGCAT |
| PP_2428/sotB | pTE922 | CACCCATGAACGGACCCATAC | ACGTTGGCCCCCTGCAAGCGGGTGGCTGGCGTGCGACCTGTTGCCGCCGAGGCCGCTCAGG CCCCTTGTCGGTCAAGTAGAGGCTCAGGC |
| PP_2483 | pTE1014 | TCGCTGCCGAAGCGCTGCCCG | TCTTCTACGGCCAACCTTCTTTAGGGGGCAGCATAGAATTTCAAC TCAGCGCGCTGCCGGTAAACGTAGCGTCCTGCTTGTCCGCCAGCG |
| PP_2667 | pTE1015 | CCTGCTGCAAGTCTATGCCTT | TTTGCCCGCCTGACCAACGCCCCCTCAGGAGCCAACGCCCGatgAAAACCTGGCGTGTAAC CCGTGTGGGAGCGGCCTTGTGTCGCGATT |
| PP_2669 | pTE913 | AGCTGCTTCGCATCGATGAC | TGTAGCTACCCCTGACCCGACAACCACAACAATAAGGAATCCGCCAtgaACGCCCTCGACG TCAGTGACGTGAGCTTTGCCTATGGCCAG |
| PP_2669 | pTE913 | AGCTGCTTCGCATCGATGAC | TGTAGCTACCCCTGACCCGACAACCACAACAATAAGGAATCCGCCAtgaACGCCCTCGACG TCAGTGACGTGAGCTTTGCCTATGGCCAG |
| PP_2710 | pTE993 | GACACCCTGCAAGTACAAACC | GCGTTACCACACCCTGTTGTCCGACATCTCGTTGTGAGGTTCACA AGATGTTCCAGGGCTTGCCCGCGTGCCGGGCAGCAAGAGGTGCACC |
| PP_2739 | pTE926 | TTCAGGGCGAGGATCACGACC | CTGCAAGGAAGCCTTTGGCGCACATCGCATACGAAATGCCTGTAGCAAAAAACCTATACAA GACCGTTGACTTGTATAGATAAAGGCTTA |

| Gene target | Plasmid No. | gRNA Targeting Sequence 5'-3' | Recombineering Oligo 5'-3' |
|---|---|---|---|
| PP_2869 | pTE994 | ACCCTGGGCGGGAGATCATGG | ACACTTTTATAAGTAAAAATAACAACACCGCACCGGAGCCGCCTG<br>GGCTGTACATACTGCGGCTTTTGCGATGGACCGACGCGCATGAGC |
| PP_2910 | pTE768 | CCTGTTCCTGCAGGCCGACC | GCTATGGTGAGCCTGCCAGGCAATGCCTGCGTGTGAGGTAAGGCTCAGCCGGGTAACACCT<br>TCTGCGGTTTCGTTTTACTCAACCCGTAC |
| PP_2940 | pTE976 | CCGGGACCATGAGCCCGCCGT | TCCAAAGAGCAACCCTTACCCGGGCAGAGCCTGGAGGATGGACAAAGCCCGCGACAAGGTA<br>AGCGTGAAGTTCACCAAGGAGGGCTGGGA |
| PP_2974 | pTE978 | ATCAGCGTCTCAACGCATTGG | TCCTGTTATAGCTCCCCATTCTTAAATTATAAAAGGCTACCCATTTCCCCCCTGAAATCAG<br>CCCTCGTCAGAAGTAGAATTTTCTCCTAA |
| PP_3017/ogt | pTE983 | CCGGTACTGAGTTCCAGCGCC | TGTGGATGCGTACAGTGGGCCGATCGCAACTTGCAGGAGCCCGCTGGCGTCAACCGAACCC<br>GCTGGCGATCAGGCAGGCCAGCACGGCCA |
| PP_3161 | pTE920 | ATCGCGCGCAACAAGGACGG | CCCTAACCTGACCAGGTCTAAACAATAACAAGGGAGACCCTGGCCAtgaGCCTGTATGACA<br>CCGTGCGCGACTTCCTGTACCGCGAAGCG |
| PP_3168 | pTE917 | CGACGAGCTCAAACTCACGC | TTCCCAACGCTTTACCCATAAAAACAAGATCAAGAGAGCTTCGGAGCCATCCAGGAGAATT<br>GCGCATCGGGGGTGTATGGGGGTAGCGTG |
| PP_3221 | pTE974 | CGCGCTGGCCCCGCACTGGCT | CCTACCTCGACCCGCGCATCACCCACACCGCGAAGGTGAACTGAGAtgaGCCAGTCACCCC<br>TGATCGAGGTCCGCGACCTAAGCGTCAGC |
| PP_3400/alkB | pTE999 | GGTCAGCCGATCGTGTCGGTG | TAGCTGTCGGGTACAATGGCAAACCGCTCGAAGATGTGCAGCCACACAGGGGCCGTCATCT<br>GCCACCCAGCGCAAAACAAGATGAACGCA |
| PP_3408/cbtA | pTE981 | GCGTTGCCCCAGGTCGGCGGC | GACACCCGCCACAGCGCCGCCTTCCCCTGCCACTGAGTACGCCTGTtgaTGCCTGCCTTCT<br>ATGCCGGTTTTGGCTGCCGCCGGGGTTGC |
| PP_3480 | pTE985 | TCAAGGGCCTGCTCAAGCAAC | TTCACCCTCACCGCCACCTGGGAGACCGGCCatgCGCGCCCTTGAaTGAACACACAACGTG<br>CAGTCATCTGGGCCGGCTTCCTCGGCGTG |
| PP_3496 | pTE929 | CGTGGTGACCAGCACGCCGT | TCGACCGGGCTGGTGTGCTCGCGTAGGCTTGCCGTTTTCCTTCCGGCTGCGCTTGTGGGGG<br>TGCGCAGCGCCCCGACATCGCTGATTGGT |
| PP_3641 | pTE937 | TCGAGAATCACACGGTCGATT | GGTTTCCTTACAAAGATAAAAACGACCTTTCGAGCTGAAAACC<br>CAACTGCAACATGGGGCTGCAAAGCAGCCCCGACTATGGCCGC |
| PP_3883 | pTE925 | GCTGCCTCCAGTCAGGCCTGA | GGCTACTCCGACCTGTTCTTGTCTCGTAAAGCGCAGGGGTGACTCAAGCCTCCGGGCGCTG |

| Gene target | Plasmid No. | gRNA Targeting Sequence 5'-3' | Recombineering Oligo 5'-3' |
| --- | --- | --- | --- |
| | | | CGGGGCCCCATGAAAATGGGTAGGGGGGT |
| PP_4214 | pTE996 | GCCACCACCATGACCCCAGGT | TGCACGCCCGACCGCCACCTCTTCCCGTTGAGCTGCGAGCCGCCC TTTTTCTTAGCCAACAGCTGACGGAATCTACTTCATGTCCGATGA |
| PP_4322 | pTE997 | CCGCCGACCCGTCGCGGGGTA | ACGCAGCACTGGTAGTCCCCGAACTGGGCCAGTTGGCGATGATTC AtgaAGCGTTGGATCATGGTAGTGCCCCTTGCAGTGTTCCTGCTG |
| PP_4373/fleQ | pTE609 | TTCGCCGAGGAAGTTCAGCAC | CACGCCCCTGCAGGGCGCCGATATGACTAGGGAAGTTGCTATTGC TGTTTGGGCTGGGTGTTCGCCTTGGGGGTTGCGTTTGGGCGGGGT |
| PP_4462 | pTE986 | GCGCTGCCTGCTCAAGACTCC | GCGCAAACAAATTCCGCTTGCTTCGCACAAGGTACTTCAGAAAtgTTGGAATTGACCCACC CTTCAAGCCTTGCTATGCGGGATGGAAAA |
| PP_4485/hisQ | pTE987 | GCGCCTACCTCTCGGAAACCT | CTGAGGTTTGCACCATTTTTCATTCTCTAGGTCGAGGACCTCATCAtgaTCTTCGACTACA ACGTCGTGTGGGAAGCGCTGCCGATGTAC |
| PP_4592 | pTE998 | CTGTGCTGATGCTCTTCGCGT | TGCCATCTGGCCTGGGGTGATCATCACTCTTGATAGCGAACGCTGAGCCCTGTGGGAGCGG GCACGCCCGCGAACCCGGGCAAACCCGGG |
| PP_4622 | pTE930 | CAACAAACCGCCAGTGACTA | CAAGCTATAAAGGCGACACGATAAATCCGGAACCTCCCCTCCC CAGAGCCTGCTCACGGCCGCACGAACCGTGAGCTGGCGCATGGG |
| PP_4706 | pTE1002 | tgTATAGCCTGTTCATAAAGA | TTTGTATACAATTCATTATTGAATTTAAAACAAAAGGAGAGATTTGCGCAAGAAAAAGCCC TGGCACGGGCCAGGGCTTCGATGAAAGTG |
| PP_4842/urtB | pTE914 | TGGATGAACAGAATGATCAA | GGCTCACCCGCGAAGAATCCAACACCGATTTCCAGGATTGCCCGCaTGAACCAGCCACTGC TTGTCACTGCTACGCAAAAGGCCGGCCCA |
| PP_4902 | pTE969 | CTCTACCGCCACATGCGCAAC | CTTGGCCACTAGGCTAATATGACCGCCTATATGAGGAGCCCCTGC GTCGAAACGGCGCAGATATTGTATCTGCGCCCCCTTTTGGTGCCT |
| PP_5003* | pTE504 | CGAGGCCTGCCGCTGTAGCTC | CGCCACAGCAACCGGTACTCGTCTCAGGACAACGGAGCGTCGTAG GCCTGCTGGAGATGTAGTGTTGCAGCCGACGCATTCGCGGGTAAA |
| PP_5007 | pTE918 | GTCGCGCGCTGCAGCAACCAA | ATGAGCTGCTTGAGCGCGTCGCGCACAAACCATAAGGAGAGCAGGTACCGCGTCGCCTTCT TCGCGGGTAAACCCGCTCCTACAAGATGC |
| PP_5032 | pTE950 | GGCCTGCTGGCCTCGACCCGC | TACCCACAAGCGGGCCTGCGGCTCCACGCGATTTGGAGTAGTAACCGCCCCTGGGGCCGCT GGGCGGCCCTTCGCGGGCGCGCCCGCTCC |

| Gene target | Plasmid No. | gRNA Targeting Sequence 5'-3' | Recombineering Oligo 5'-3' |
|---|---|---|---|
| PP_5041/glgP | pTE982 | TCATCGGCCGCTTGCTGTACG | TCAACACTCTCCCTCCGCTTTCGTCAACTTGCCCGAGGATGCCGCGGTGAGGCCGTTCGCGGGCACGCCCGCTCCCACAGGTGTCTCGCA |
| PP_5094 | pTE923 | CGCGATTTCCGCCCGTATCG | AAGCAGGTAGAATGCCGCAGACCCCACAGCTGCCGGCATCGATCCTTCCATGCTTCACTCACTCTTTTCCACAAGGACCTGACATGAGCA |
| PP_5244 | pTE911 | TCCCGCGCGTTCAAAGTCAA | GCGCACAATTAACAACAATTCAGGTTTGGCTCAGGACTTCGTCATGCCAATGCTATCCACCGGTATAACCCCTGCGGACAGTGCCGGCCC |
| PP_5391 | pTE921 | taacagtgttaaggagatgat | AACGGGATCAGCTTCGAGCCGCCGAGTGCCCAATAGCCCGTTGATCCCACCTCCTTTCCCAGGAGTCGCTATGAACCCTCACTTCGATGA |

### The PP_1697 gRNA targets the upstream promoter sequence for a DSB, not the ORF.

*Previously described in (Czajka et al. 2022). Accession ID: JBEI-204,825
